# Extracellular matrix context shapes morphogenesis and lactation-associated states in human milk-derived mammary organoids

**DOI:** 10.64898/2026.08.23.746503

**Authors:** Amelia Hasenauer, Valerie Pascetta, Maxwell C. McCabe, Anthony J. Saviola, Simone Ponta, Mehmet Yilmaz, Verena Bossung, Christina L. Coelius, Thomas Biesgen, Kirk C. Hansen, Stefan Prekovic, Nicole Ochsenbein-Koelble, Marcy Zenobi-Wong

## Abstract

The mammary gland relies on reciprocal interactions between epithelial cells and their surrounding extracellular matrix (ECM) to form and maintain milk-producing tissue structures. Yet these processes remain difficult to study in human model systems. Mammary epithelial cells (MECs) can be isolated noninvasively from breast milk, but whether they generate three-dimensional organoids and respond to matrix cues has been unclear. Here, human milk-derived MECs (milk MECs) spontaneously form complex organoids, including polarized acinar and terminal duct lobular unit-like structures after isolation. To investigate how matrix composition shapes these organoids, milk MECs were cultured in decellularized mammary ECM (dECM), Matrigel, and collagen I. In dECM, milk MECs formed polarized branched networks with aligned actin organization along collagen fibrils, whereas in Matrigel they adopted a more lactation-associated state, marked by β-casein expression and milk fat globules. Together, these findings establish breast milk-derived MEC organoids as a human model to study how ECM context regulates mammary morphogenesis and lactation biology.

## Introduction

The human breast establishes and maintains a ductal network with secretory alveoli that are repeatedly remodeled across pregnancies to support lactation [1]. Disruption of mammary developmental programs or lactation-associated remodeling can not only compromise milk production but also bias the tissue toward cancer-associated phenotypes [2–4]. However, the mechanisms by which extracellular matrix (ECM) cues regulate human mammary tissue organization and function remain difficult to interrogate systematically. Most insight still derives from animal models, which do not fully recapitulate human physiology and in humans, studies of lactating mothers are typically observational and largely correlative [5, 6]. Primary human mammary epithelial cells (MECs) embedded in defined ECM niches could provide a controllable *in vitro* system to dissect how the ECM regulates tissue homeostasis and function [7].

The mammary epithelium develops within a dynamic stroma, whose biochemical composition and physical properties instruct morphogenesis [8]. Across puberty, pregnancy, and lactation, stage-specific hormones intersect with matrix-derived signals to coordinate epithelial fate decisions that bias development toward ductal or alveolar programs [9, 10]. Within the mammary niche, β1-integrins on the cell surface engage the basement membrane to establish apico-basal polarity and maintain ductal integrity, while fibrillar collagens (I/III) provide guidance cues for collective migration [11–13]. Concurrently, protease-mediated remodeling and enzymatic crosslinking reshape matrix stiffness and viscoelasticity, which the epithelium senses via mechanotransduction pathways to regulate tip motility, branching frequency, and lumenization [2, 9, 14]. Together, biochemical signaling, ECM remodeling, and mechanosensing pattern the mammary epithelium through principles that are broadly conserved across branching organs[15]. When dysregulated, ECM stiffening and remodeling can compromise epithelial integrity, promote cancer invasion, and contribute to therapy resistance [3, 16].

Early evidence that mammary epithelial behavior is instructed by ECM cues motivated the development of three-dimensional culture systems. Since the 1980s, MECs have been embedded in ECMs to generate organoids, self-organizing structures derived from stem or progenitor cells that reproduce key aspects of tissue architecture *in vitro* [17, 18]. Among the most complex examples, Yuan et al. reported an *in vitro* “mini” mouse mammary gland that recapitulates ductal-alveolar organization, cancer progression and functional features of the native tissue [19–21]. Despite substantial progress, important challenges remain in translating these systems to human mammary biology. State-of-the-art models remain mouse-derived and reflect a pubertal program in which terminal end buds drive ductal elongation [22, 23]. By contrast, the adult mammary gland culminates in terminal duct–lobular units (TDLUs), the functional epithelial units of the breast[22]. Mouse and human mammary glands also differ in developmental dynamics, epithelial organization, and stromal context. In mice, the mammary gland is embedded in a compliant, adipose-rich stroma, whereas the human TDLU niche is comparatively fibrous and collagen-dense [22, 24]. This stromal mismatch alters tissue mechanics and epithelial-stromal signaling in ways that are difficult to reproduce in standard organoid culture with mouse sarcoma-derived Matrigel [25]. These limitations underscore the need for human-relevant in vitro models of the adult breast microenvironment. To address this gap, we developed a human milk-derived epithelial organoid model in decellularized mammary matrices that incorporates both a human epithelial source and a native-like matrix context.

Human milk provides a noninvasive source of primary mammary epithelial cells (MECs). Because milk can be sampled longitudinally from the same donor, milk-derived MECs (milk MECs) enable biopsy-free, within-donor study designs that reduce inter-individual confounding[26]. Single-cell studies have identified lactocytes, basal/myoepithelial cells, and progenitor-like cell states that shift across lactation [27, 28]. Most work to date has emphasized isolation protocols and cell atlas efforts, with few examples of 3D-cultured milk MECs used to interrogate cell-ECM interactions [28, 29].

Prior work on 3D printed “ductal-alveolar” scaffolds for *in vitro* lactation studies showed that milk MECs form stable epithelial layers, exhibit lactation-associated outputs, and respond to matrix stiffness [26, 30]. However, scaffold-based systems often confine cells to predefined shapes and limit their capacity to interact with and remodel the surrounding microenvironment. In contrast, organoids permit branching and lumenization to emerge de novo, and enable readouts of how ECM cues guide morphogenesis toward ductal versus alveolar-like programs [23].

In parallel, advances in biomaterials now make it possible to reconstruct more human-relevant microenvironments. Decellularized ECM (dECM) hydrogels, produced by the removal of cells and nucleic acids from native tissue, retain tissue-specific ligands, fibrillar architecture, and physiologic mechanics [31, 32]. Across epithelial organs (e.g. lung, skin, and intestine) dECM can be formulated as self-gelling hydrogels to support long-term cell culture [33, 34]. Organ-matched dECM has been reported to better maintain epithelial differentiation and function than commercial matrices, which highlights the value of native ECM cues for maturation and phenotype stability [33]. In breast tissue engineering, decellularized materials are also emerging as practical, scalable alternatives to commercial gels for *in vitro* tissue models, drug screening, and GMP-aligned translational applications [35]. Mammary dECM can encode microenvironmental “state” information; for example, aged or irradiated breast dECM hydrogels have been shown to modulate cancer invasion and proliferation programs, which underscores that ECM history can be functionally instructive. In recent studies, mammary dECM has been formulated as bioink for 3D-printed tumor organoids, and as scaffolds for breast cancer models [35, 36]. Thus, mammary dECM offers a promising physiologically relevant matrix to study how human-relevant biochemical ligands and mechanical cues regulate MEC-ECM interactions and lactation *in vitro*.

In this study, we embedded milk MECs in self-gelling mammary dECM derived from human breast and bovine udder, alongside collagen I and Matrigel hydrogels, to establish a mammary tissue– informed organoid model. This platform enables controlled analysis of how matrix context shapes mammary epithelial architecture and lactation-associated function, with milk MECs as a practical primary human cell source. Together, this work provides a human-relevant organoid system with potential applications in disease modeling, biomaterials testing, and translational breast research.

## Results

### Breast milk–derived MECs generate complex polarized mammary organoids *in vitro*

Milk samples were collected from lactating donors to establish *in vitro* milk MEC organoids. The total number of cells per milliliter of milk was not associated with maternal BMI or gestational age at birth but was reduced at later postpartum timepoints (late infancy), in donors with two versus one child, and in afternoon versus morning samples (Fig. S1, S2A-E and Table S1). Isolated cells had high viability (>90%) and were either plated in 2D tissue culture flasks or encapsulated in 3D Matrigel to initiate primary milk cell cultures (Fig. 1A and Fig. S2F). Previous flow cytometric analysis of freshly isolated human milk cells showed that MECs comprised only ∼11% of the total population at isolation but were enriched during culture [26, 27]. The isolated epithelial cell fraction gave rise to colonies in 2D and organoids in 3D, and cultures were stained with cytokeratin 8 (CK8) and cytokeratin 14 (CK14) to validate the presence of luminal and basal mammary epithelial lineages (Fig. 1A). Epithelial cultures were successfully established from most samples (9 of 11) and were expandable in both 2D and 3D conditions, with higher cell yields following 2D (1 million cells) rather than 3D (0.4 million cells) expansion (Fig. 1B-C and Fig. S3).

**Figure 1.**
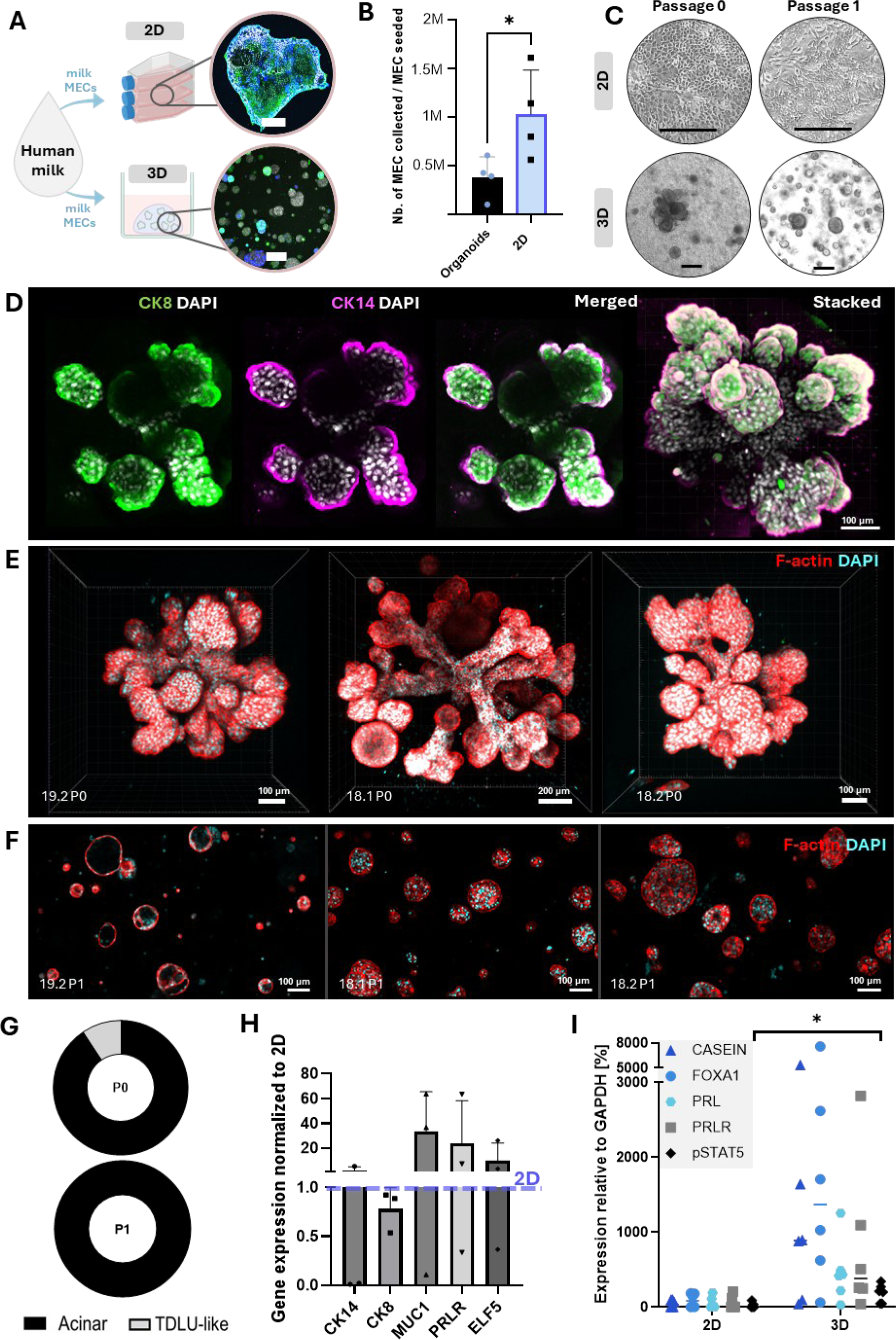
Establishment and characterization of human milk-derived mammary epithelial cultures (milk MECs) in 2D and 3D. **A)** Schematic overview of the workflow: human milk–derived cells are processed and cultured either as adherent 2D mammary epithelial cultures or embedded in Matrigel to generate 3D organoids. Representative fluorescence images of a 2D colony (top) and 3D organoids (bottom) are shown. **B)** Collection of mammary epithelial cells (MEC) expressed as the number of MEC collected relative to the number of MEC initially seeded for 3D organoids versus 2D culture (Mean ± SD, n = 4 biological replicates; *p < 0.05). **C)** Representative brightfield images of cultures at Passage 0 and Passage 1, showing 2D morphology and corresponding 3D organoid growth. Scale bars: 750 μm **D)** Immunofluorescence characterization of 3D organoids stained for cytokeratin 8 - CK8 (luminal marker; green), cytokeratin 14 - CK14 (basal marker; magenta), and DAPI (nuclei; white). Single channels, merged view, and a 3D stacked rendering are shown. Scale bar, 100 µm. **E)** Representative 3D reconstructions of TDLU-like organoids at P0 from independent samples stained for F-actin (red) and DAPI (cyan/white), illustrating complex organoid architecture. Scale bars as indicated. **F)** Representative organoids stained for F-actin (red) and DAPI (cyan/white), forming acinar structures at P1. Scale bar, 100 µm. **G)** Quantification of organoid morphologies at Passage 0 (P0) and Passage 1 (P1), classified as acinar or TDLU-like (terminal duct lobular unit-like). **H)** RT-qPCR analysis of lineage/lactation-associated genes (CK14, CK8, MUC1, PRLR, ELF5) shown as expression levels normalized to 2D (Mean ± SD, n = 3 biological replicates; *p < 0.05). **I)** Western Blot comparing 2D and 3D cultures from donors for lactation-related signaling/markers: pSTAT5, PRLR, FOXA1, PRL, and β-casein; GAPDH serves as a loading control.

In 2D, colonies displayed epithelial colonies with cobblestone, stratified, and refractive-edges morphologies, consistent with our previous study [26]. In 3D, milk MECs formed both acinar and branched structures within 14 to 20 days of culture post isolation (Fig. 1C). Branched morphologies were observed in 6 of 11 milk samples and retained apico-basal polarity, with CK8 localized to the luminal compartment and CK14 to the basal layer (Fig. 1D; Figs. S3–S4). F-actin staining further highlighted the overall architecture of these branched organoids, with duct-like extensions that terminated in budded tips (Fig. 1E). Across independent cultures, these TDLU-like structures comprised 2.8–41.7% of organoids (mean 17.1%) at P0 (Fig. 1E, G). With culture passages, however, this TDLU-like organization was lost, and organoids converged to exclusively acinar morphologies (Fig. 1F-G). Similar budding structures, and their progressive loss with passaging, have been reported in organoid cultures derived from tissue-derived MECs [37].

To assess the influence of 3D culture on milk MEC lineage state and differentiation, RT-qPCR markers associated with basal identity (CK14), luminal identity (CK8 and mucin 1; MUC1), and alveolar/lactation competence (PRLR and E74-like factor 5; ELF5) were examined. Expression of basal CK14 remained similar between 2D and 3D. In contrast, MUC1, PRLR, and ELF5 trended to be higher in 3D relative to 2D, indicating a shift toward an alveolar-like program, despite a slight decrease in CK8 expression (Fig. 1H).

Western blot analysis was performed on milk MEC cultures (P0-P1; n = 6 independent donor samples) to determine whether the observed transcriptional trends were accompanied by changes in differentiation- and lactation-associated protein expression (Fig. 1I). β-casein, a major milk protein and marker of lactational differentiation, forkhead box A1 (FOXA1), a transcription factor associated with luminal epithelial identity, PRL, a key lactogenic hormone, PRLR the corresponding receptor and phosphorylated signal transducer and activator of transcription 5 (pSTAT5), a downstream effector of PRL signaling, were all expressed at low levels in 2D cultures. In 3D organoid cultures, several donors showed higher abundance of these proteins, although the magnitude of the increase varied across donors. In particular, pSTAT5, PRLR and β-casein were higher in many 3D samples than in 2D cultures, consistent with the acquisition of lactation-associated features (Fig. 1I and Fig. S5).

### Self-gelling ECM for organoid culture can be extracted from mammary tissue

Physiologically relevant ECM microenvironments may preserve or reinstate TDLU-like architectural complexity, rather than driving a spherical (acinar) growth program *in vitro*[38]. Tissue-specific ECM hydrogels were generated from human breast and lactating bovine udders (Fig. 2A). Breast dECM enhances relevance to the adult human mammary niche, while lactating bovine udder provides an abundant, accessible source of lactation-stage matrix[26]. Native bovine and human mammary tissues exhibited dense cellularity, but cells were efficiently removed via decellularization (Fig. 2B). The mammary ECM-derived materials were benchmarked against standard substrates: Matrigel, a laminin/collagen IV-rich basement membrane mimic commonly used in standard organoid culture, and rat tail collagen I, which represents a fibrillar interstitial matrix (Fig. 2A)[23]. After enzymatic digestion the human and bovine decellularized matrices contained a broad range of protein bands distributed more similarly to collagen I than to Matrigel (Fig. 2C).

**Figure 2.**
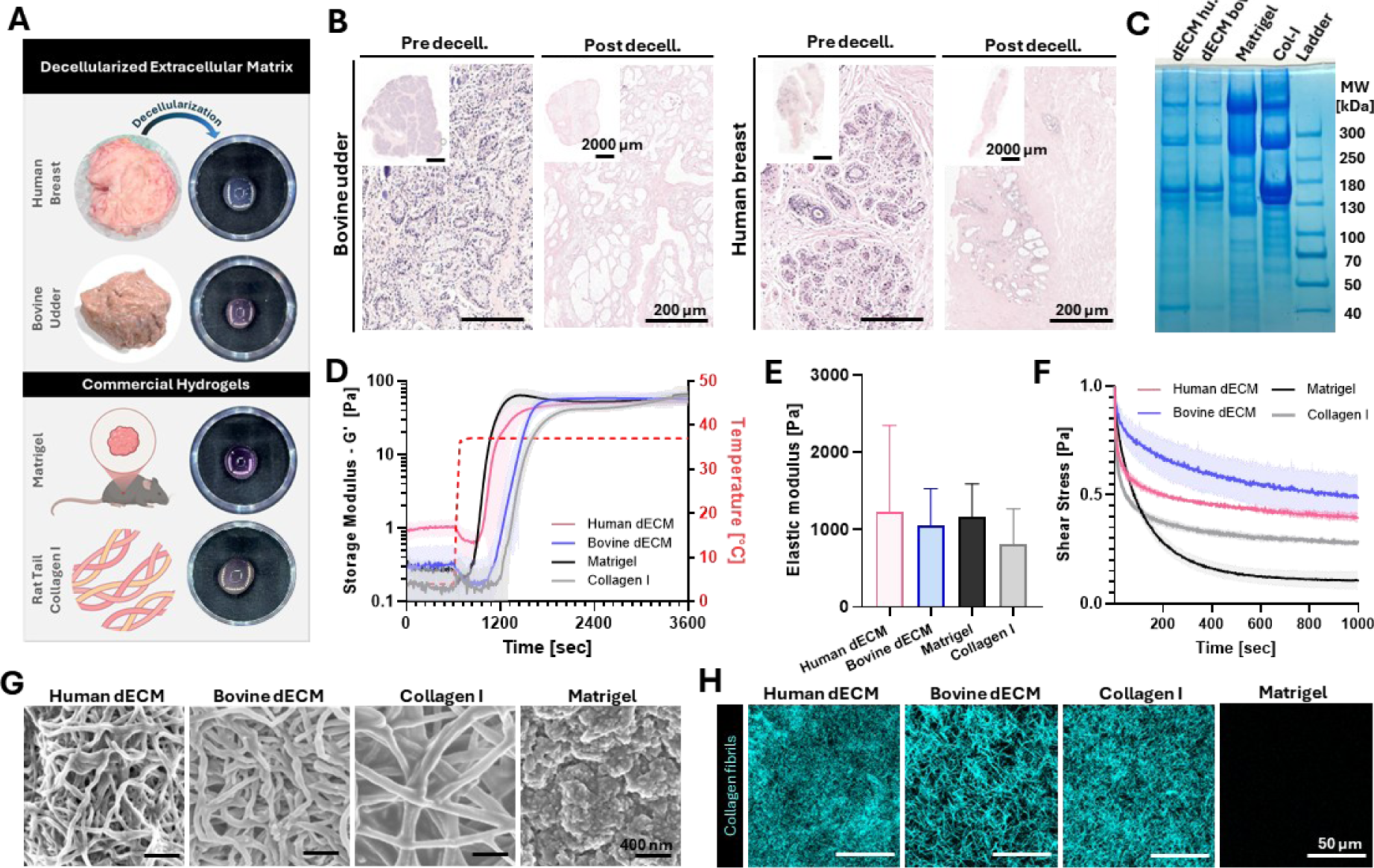
Extracellular matrix hydrogel characterization. **A)** dECM can be extracted from human and bovine mammary tissues and converted to translucent acellular hydrogels with similar self-gelling properties to commercial Matrigel and rat tail collagen I. **B)** Representative Hematoxylin and Eosin (H&E) staining of bovine udder (left) and human breast (right) tissue sections before and after decellularization, demonstrating removal of cellular components while preserving overall tissue architecture (scale bars as indicated; low magnification: 2000 µm; high magnification: 200 µm). **C)** SDS–PAGE protein profiles of human dECM, bovine dECM, Matrigel, and collagen I (ladder shown at right). **D)** Oscillatory rheology during a 4 → 37 °C ramp shows all hydrogel materials reach a comparable plateau storage moduli (G’) upon thermal gelation (means ± SD, n = 4 technical replicates). **E)** Compression testing of the four ECM hydrogel materials (means ± SD, n = 3 technical replicates). **F)** Stress-relaxation behavior under constant strain (means ± SD, n = 3 technical replicates). **G)** SEM reveals interconnected fibrous networks in dECM and collagen I materials, and an amorphous, non-fibrillar matrix in Matrigel. **H)** SHG images highlight differences in collagen fiber microstructure (dECM and collagen I) or lack thereof (Matrigel) between matrices (collagen fibers = cyan).

Because matrix mechanics strongly influence cell behavior, reproducible thermal gelation and mechanical characterization of the extracted hydrogels are essential for interpretable 3D culture experiments. dECM precursor solutions were centrifuged to remove undigested particles which yielded homogeneous hydrogel solutions (Fig. S6A–B). Upon warming to 37 °C, both human (6 mg/mL) and bovine (8 mg/mL) dECM rapidly increased in storage modulus (G′) within ∼20 min and reached plateau values matched to Matrigel (9 mg/mL) and collagen I (2.6 mg/mL), with G′ ∼ 90 Pa (Fig. 2D). Storage conditions influenced gelation kinetics, with frozen dECM preparations gelling more rapidly (∼20 min) than lyophilized/reconstituted material (∼25 min). However, both converged to similar plateau storage moduli and supported cell growth (Fig. S6C–E). Optimized human and bovine dECM hydrogels reached elastic moduli of ∼900–1100 Pa, with no significant differences compared with Matrigel and collagen I (Fig. 2E). Under constant strain, all hydrogels exhibited viscoelastic stress relaxation. Bovine dECM relaxed more slowly and maintained higher residual stress than the other matrices, whereas Matrigel relaxed most rapidly (Fig. 2F).

Epithelial cells are sensitive to not only mechanical properties, but also the matrix’s micro- and nanoscale structure, which shapes cell mechanotransduction, polarity, and migration[2, 9]. Analysis of hydrogel architecture showed that human and bovine dECM formed fibrillar networks similar to collagen I, whereas Matrigel exhibited a more amorphous microarchitecture (Fig. 2G and Fig. S7A). Accordingly, collagen fibrils were prominent in both dECM and collagen I gels, whereas no fibrillar collagen signal was detected in Matrigel, consistent with its composition as a basement membrane extract enriched in laminin and collagen IV (Fig. 2H and Fig. S7B).

### Human breast and bovine lactating udders have distinct proteomic and biochemical profiles

To determine whether the dECM materials retained distinct biochemical cues, decellularized human breast tissue was profiled by proteomics and integrated with previously acquired bovine udder datasets for direct cross-tissue comparison[26]. Proteomic analysis identified 201 soluble and insoluble proteins in decellularized breast, with 88 matrisome (ECM) and matrisome-associated (ECM-associated) components, 62 of which were shared across all three donors (Figs. S8-S9). Non-ECM proteins accounted for <5% of total summed intensities and were excluded to focus consequent analysis on cell-matrix-relevant cues (Figs. S9-S10). Fibrillar collagens (e.g., COL1A1/COL1A2) are crucial for maintaining tissue integrity, and dominated the ECM signal in both species, alongside additional stromal collagens (e.g., COL3A1, COL5A2, COL6A) (Figs. S10-S11). To resolve tissue-specific collagen composition, collagen subtype-profiling was performed (Figs. S10-S11). Human breast dECM was richer in collagen I (69.6% of collagen signal) relative to bovine mammary dECM (52.0%), whereas bovine dECM contained a markedly higher fraction of collagen II (15.1% vs 0.7%), with additional tissue-specific collagens detected (human: VII/XXI; bovine: XVII) (Fig. S12). Matrisome subcategory analysis further detailed basement membrane, elastic microfibril, and small leucine-rich proteoglycan components in human breast dECM (Figs. S13-S15).

To further dissect differences between bovine and human mammary matrices, unsupervised multivariate analysis was performed. Principal component analysis (PCA) of matrisome proteins separated lactating bovine udder and human breast dECM into two non-overlapping clusters (Fig. 3A). Principal component (PC) 1 explained 73.3% of the variance, consistent with strong tissue-specific differences in ECM composition, while PC 2 accounted for 11.6% and reflected within-group variability. This distinct grouping was also observed in PCA of specific ECM sub-compartments, such as collagens and basement membrane components (Fig. S16). Additionally, a heatmap of the 25 most abundant matrisome proteins revealed tissue-specific abundance patterns that were consistent across donors (Fig. 3B and Fig. S10). Laminins (e.g., LAMB1, LAMB3, LAMA5) and basement membrane-associated proteins (e.g., COL18A1, COL4A6, MATN2, HAPLN1, COMP) were among the strongest discriminators between matrices (Fig. 3C–D and Fig. S17). Together, these data indicate that the two decellularized matrices preserve distinct signatures rather than converging on a shared generic matrisome profile.

**Figure 3.**
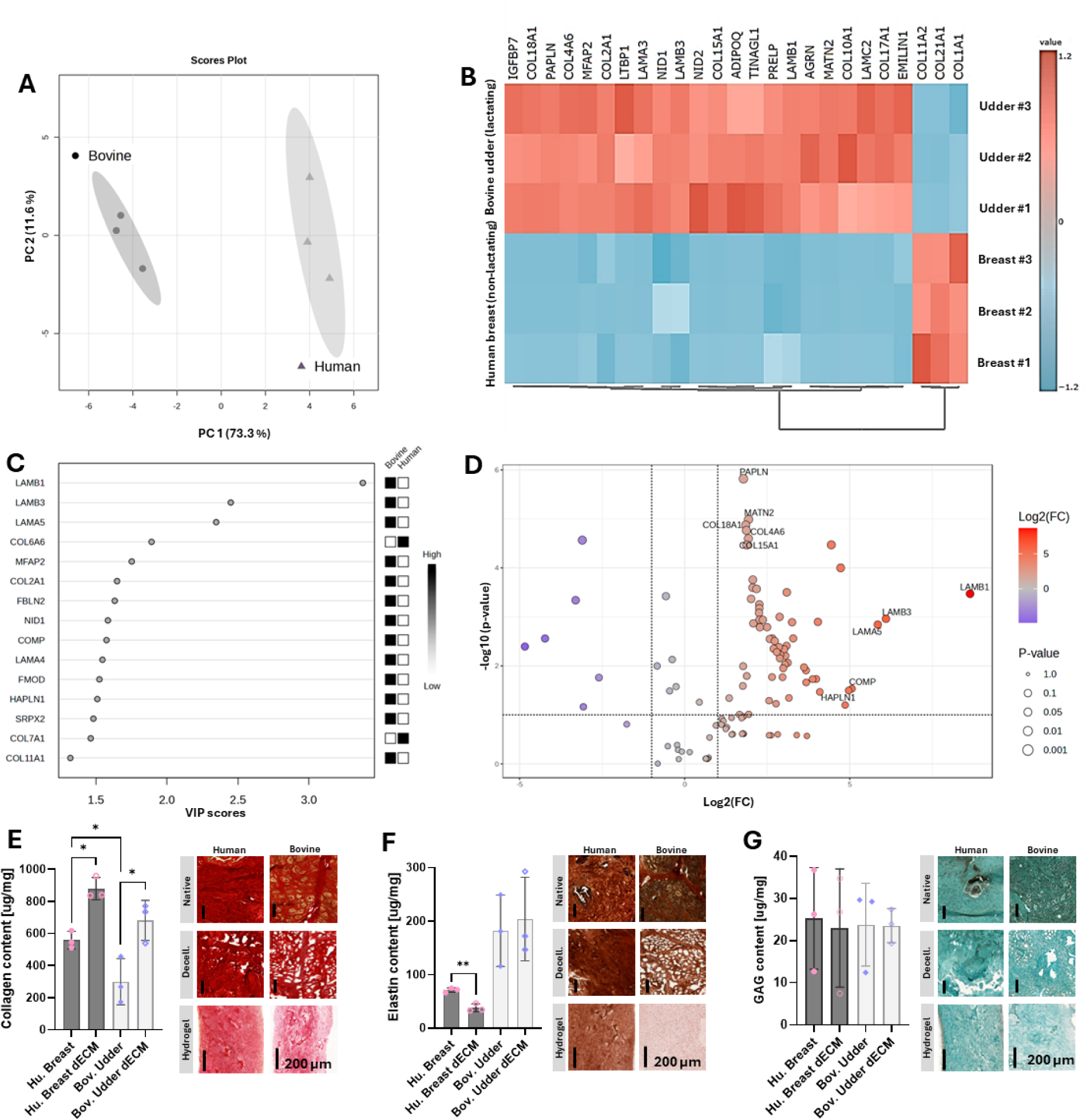
Human breast and lactating bovine udder dECM show distinct matrisome signatures and biochemical composition. **A)** Principal component analysis (PCA) of ECM-associated proteins from label-free proteomics separates human breast and lactating bovine udder dECM into non-overlapping clusters (percent variance explained indicated on axes; n = 3 per group); shaded ellipses denote 95% confidence interval for each group. **B)** Hierarchically clustered heatmap of the 25 most abundant ECM-associated proteins showing species-specific abundance patterns conserved across samples; values are row-scaled (z-scored) relative abundances (red, higher; blue, lower). **C)** Partial least squares–discriminant analysis (PLS-DA) variable importance in projection (VIP) scores highlighting top proteins driving separation between human and bovine dECM proteomes. D) Volcano plot of differential abundance (human vs bovine); points are colored by log2(FC) and sized by P value, and selected ECM proteins are annotated. Multiple ECM proteins are significantly enriched in one matrix versus the other, highlighting species-specific ECM cues. (E–G) Biochemical validation in native tissue, decellularized ECM, and ECM-derived hydrogels from human breast and bovine udder. Bulk collagen (E), elastin (F), and glycosaminoglycan (GAG) content (G) were normalized to dry weight (mean ± SD and n = 3 per group; *P < 0.05, **P < 0.01). Representative stains: Picrosirius Red (collagen), Verhoeff–Van Gieson (elastin), and Alcian Blue (GAG). Scale bars, 200 µm.

To validate these proteomics-defined differences, the bulk abundance of collagens, elastin, and glycosaminoglycans was quantified in native tissue and the decellularized extracellular matrices. In line with the proteomics results, collagen was abundant in both matrices and became enriched after decellularization, consistent with removal of cellular mass and concentration of matrix (Fig. 3E). Elastin content was tissue-dependent, and lactating bovine udder contained more elastin than human breast in both native and decellularized matrices (Fig. 3F), while total glycosaminoglycan content remained broadly comparable across groups (Fig. 3G). Together, these results establish our dECM materials as a mechanically stable and biochemically rich foundation for subsequent organoid experiments.

### Decellularized mammary ECM promotes polarized branching of human milk MEC

Milk MECs were first expanded to obtain sufficient cell numbers for subsequent cell-ECM interaction experiments. Cells were then cultured either on 2D substrates or embedded in non-floating hydrogels composed of human breast dECM, lactating bovine udder dECM, Matrigel, or collagen I [23]. MCF10A cells were included as a well-characterized, non-tumorigenic human MEC reference to validate matrix-dependent effects and to distinguish ECM-specific responses from donor-to-donor variability in milk MEC cultures. (Fig. S18)[39].

Consistent with our previous observations, milk MECs formed dense epithelial monolayers in 2D (Fig. 4A) (29). In 3D, morphology depended strongly on matrix context. Matrigel supported predominantly spherical organoids, whereas human breast dECM, bovine udder dECM, and collagen I promoted branched, frequently interconnected epithelial networks (Fig. 4A,C). Area-based morphometric analysis confirmed that spherical structures accounted for ∼100% of signal in Matrigel, compared with 9% in human breast dECM, 21% in bovine udder dECM, and 1% in collagen I. Accordingly, branched structures dominated epithelial coverage in the latter matrices (Fig. 4A,C). MCF10A cells showed the same matrix-dependent shift, with spherical organoids in Matrigel and branched morphologies in mammary dECM and collagen I (Figs. S19 and S20).

**Figure 4.**
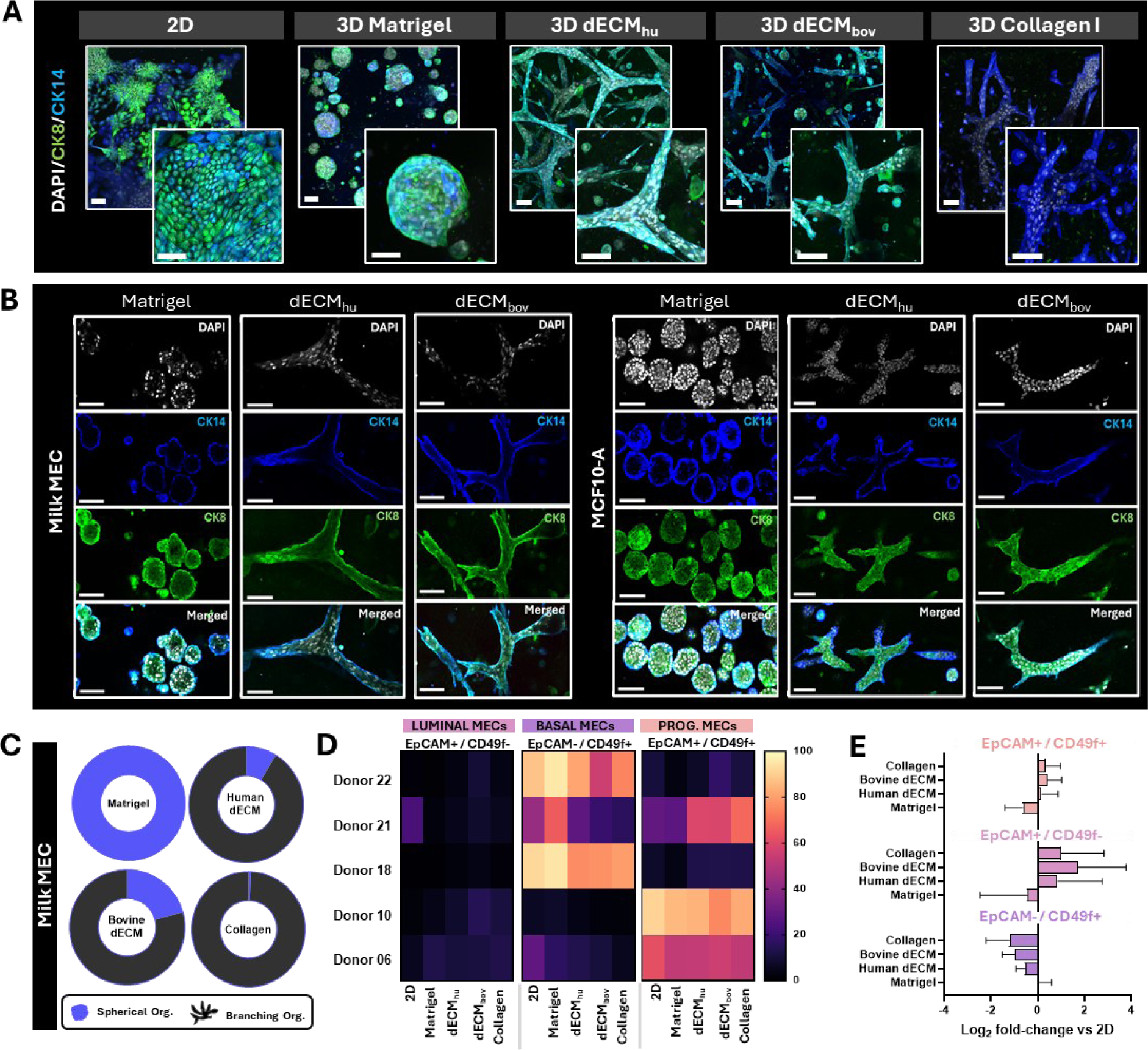
Matrix-dependent morphology and lineage composition of milk MEC. **A)** Representative maximum-intensity projections of confocal 3D z-stacks of milk MEC cultured in 2D or embedded in 3D Matrigel, 3D decellularized ECM from human mammary tissue (dECM_hu_), 3D decellularized ECM from bovine mammary tissue (dECM_bov_), or 3D collagen I. Nuclei are stained with DAPI (blue), CK8 marks luminal epithelial cells (green), and CK14 marks basal epithelial cells (cyan). **B)** Representative single-channel and merged single z-plane showing organoids generated from milk MEC (left) and MCF10A cells (right) cultured in Matrigel, dECM_hu_, or dECM_bov_. Shown are DAPI (gray), CK14 (blue), CK8 (green), and merged channels. Collagen I is not shown in panel B because the absence of CK8 signal limited interpretation of CK8/CK14 spatial organization in this condition. **C)** Quantification of organoid morphology for milk MEC cultures across matrices. Donut plots indicate the fraction of spherical organoids (blue) versus branching organoids (black) in Matrigel, human dECM, bovine dECM, and collagen I. **D)** Heatmaps summarizing the percentage of EpCAM/CD49f-defined subpopulations after culture under the indicated conditions (2D, Matrigel, dECM_hu_, dECM_bov_, collagen I) for each donor line (n = 5): EpCAM⁺/CD49f⁻ (luminal), EpCAM⁻/CD49f⁺ (basal), and EpCAM⁺/CD49f⁺ (luminal progenitor). **E)** Log₂ fold-change in abundance of each EpCAM/CD49f subpopulation in 3D matrices relative to matched 2D controls (Means ± SD, n = 5 donors). Scale bars: 100 μm.

To link gross morphology to epithelial organization, CK8 and CK14 staining was used to assess luminal and basal marker distribution, respectively (Fig. 4A,B and Fig. S19). In Matrigel, organoids showed CK8 enriched on the luminal-facing side, with CK14 restricted to the periphery. In mammary dECM, ductal-like branched structures retained a comparable spatial partitioning, with CK14 localized along the outer branch boundary and CK8 detected within the branches (Fig. 4B). This bilayer-like arrangement is consistent with an *in vivo*-like mammary epithelial organization and was further supported by staining of α-smooth muscle actin and vimentin in the outer, basal cell layer of milk MEC organoids in human dECM (Fig. S21). Despite robust branching in collagen I, CK8 remained absent, which indicated that collagen I-supported structures were morphologically ductal-like but did not recapitulate the lineage organization observed in mammary dECM (Fig. 4A). Consistent with this, both MCF10A and milk cells showed smaller, less interconnected branches and overall reduced growth in collagen I compared with mammary dECM (Figs. S19 and S20).

Imaging of luminal and basal cell localization within organoids was complemented with flow cytometry of epithelial cell adhesion molecule (EpCAM) / integrin alpha 6 (CD49f) markers to quantify epithelial subpopulations across donors (n = 5) and culture conditions (Fig. 4D). The luminal (EpCAM⁺/CD49f) cell fraction remained low in 2D and all 3D conditions (mean 4.1–9.5% across conditions; range 0.3–23.1%) (Fig. 4D). In contrast, basal (EpCAM⁻/CD49f⁺) and luminal progenitor (EpCAM⁺/CD49f⁺) populations were more prominent with 32.9-55.9% (range 1.8– 95.1%) and 34.5–45.3% (range 3.1–90.8%) respectively (Fig. 4D and Fig. S22-S23). The double-negative fraction (EpCAM⁻/CD49f⁻) increased in 3D, rising from 2.2% in 2D (0.33–5.30%) to 5.5% in Matrigel (1.21–17.4%) and ∼10–14% in ECM matrices (Fig. S24).

EpCAM/CD49f frequencies were normalized within each donor to the matched 2D condition to account for strong donor-to-donor variability (Fig. 4E). This paired normalization highlighted a consistent decrease of basal cells (EpCAM⁻/CD49f⁺) in 3D conditions, accompanied by relative increases in EpCAM⁺ luminal cells (Fig. S24). Mammary dECM showed the largest positive shifts for EpCAM⁺/CD49f⁻ and smaller shifts for EpCAM⁺/CD49f⁺, whereas Matrigel remained closer to the 2D baseline (Fig. 4E). Overall, 3D culture, most notably mammary dECM, shifted compositions away from basal dominance toward relatively more EpCAM⁺ luminal-like states, despite substantial donor-to-donor heterogeneity. This result is consistent with the increased expression of luminal/alveolar-associated markers in 3D relative to 2D reported earlier by RT-qPCR (Fig. 1H).

### Mammary dECM guides cytoskeletal organization in milk-derived MEC organoids

Branching morphogenesis is often accompanied by cytoskeletal remodeling, thus matrix-dependent organoid morphologies were next examined for changes in actin organization[19]. In spherical organoids (Matrigel) actin alignment was largely isotropic and MECs maintained E-cadherin–positive junctions. In human dECM, branched organoids exhibited actin alignment along the axis of branch extension, as shown by orientation maps, while E-cadherin expression was retained (Fig. 5A). For milk MECs, alignment increased from 26.2% in Matrigel to 61.5% in human dECM, 56.6% in bovine dECM, and 57.8% in collagen I (Fig. 5Bi). MCF10A cells showed the same shift (25.7% in Matrigel; 65.2% in human dECM; 62.5% in bovine dECM), with the highest alignment in collagen I (74.3%) (Fig. 5Bii).

**Figure 5.**
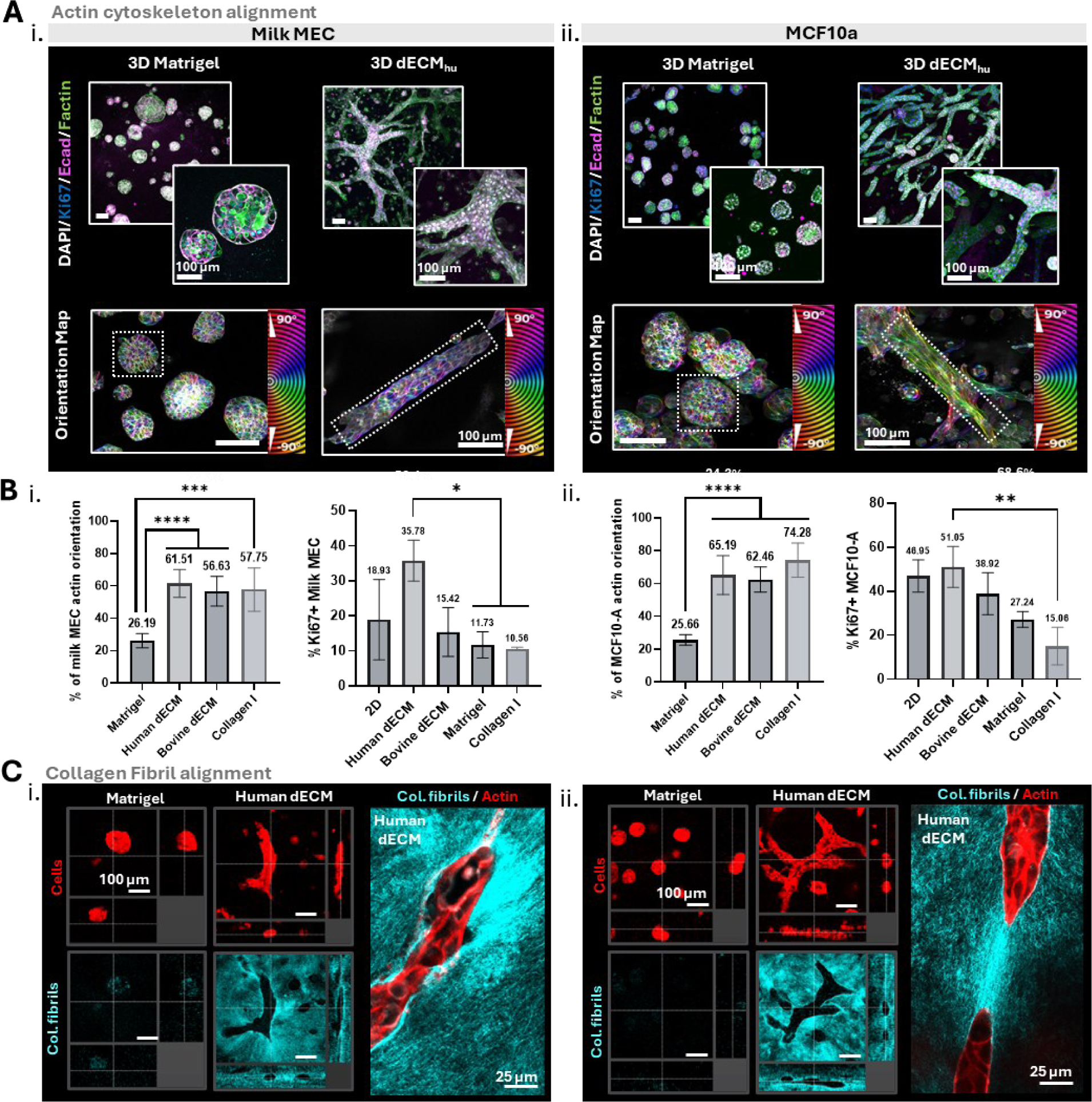
Actin alignment and ECM collagen fibril remodeling during branching morphogenesis. **A)** Actin alignment in 3D matrices. Representative confocal images of milk MEC (i) and MCF10A (ii) cultured in 3D Matrigel or human decellularized ECM (dECM_hu_). DAPI (white), Ki67 (cyan), E-cadherin (magenta), and F-actin (green). Orientation maps (color-coded by angle; color wheel). Scale bars, 100 μm. **B)** Quantification of actin alignment and proliferation. Actin alignment index (±15° criterion) and Ki67+ fraction for milk MEC and MCF10A cultured in Matrigel, human dECM, bovine dECM, or collagen I (n = 10 isolated branches/spheroids per condition). dECM and collagen I significantly increase actin alignment relative to Matrigel. Paired t-test; ***p < 0.001, ****p < 0.0001. **C)** Collagen fibril remodeling. Representative images of milk MEC (i) and MCF10A (ii) in Matrigel or human dECM, showing cells/actin (red) and collagen fibrils (cyan). Higher-magnification views highlight cell alignment along collagen fibrils in dECM. Scale bars, 100 μm and 25 μm.

Branching and actin anisotropy were restricted to fibrillar collagen-containing matrices (dECM and collagen I), and cell-ECM interactions were assessed by second-harmonic generation (SHG) imaging. SHG detected a strong collagen fibril signal in human dECM, with collagen enrichment along branch boundaries and fibril alignment ahead of advancing branch tips, whereas Matrigel exhibited no detectable SHG signal (Fig. 2H, Fig. 5C and Fig. S25A). This indicated that the milk MECs not only respond to fibrillar collagen, but also actively align and remodel the surrounding ECM during branch extension, consistent with prior reports that aligned collagen provides contact-guidance cues for branching morphogenesis (Fig. S25B)[19]. To test whether organoids also contribute to matrix assembly, metabolic labeling of nascent secreted proteins and proteoglycans revealed de novo deposition at the organoid surface, which indicated local accumulation of cell-derived ECM (Fig. S26). We then asked whether the observed morphological differences were associated with matrix-dependent growth. Ki67 quantification showed that proliferation varied by matrix, with milk MECs proliferating most in human dECM (35.8%) and least in collagen I (10.6%). MCF10A organoids showed a similar pattern (Fig. 5B).

Because branching morphogenesis is often accompanied by changes in epithelial adhesion state, E-cadherin (CDH1) and N-cadherin (CDH2) transcript levels were quantified by RT-qPCR. E-cadherin is the principal adhesion molecule of epithelial junctions and helps maintain polarized epithelial architecture, whereas N-cadherin is often associated with motility and epithelial plasticity (Fig. S25C) [40, 41]. CDH1 levels were comparable across 3D matrices in both milk MEC and MCF10A cultures. CDH2 remained low in milk MECs across matrices, with a slight increase in collagen I relative to Matrigel and dECM. In contrast, MCF10A exhibited a significant increase in CDH2 in collagen I, compared to the other matrices (Fig. S25C). Together, these data indicate that collagen I selectively increased CDH2 in milk MEC and MCF10A without a corresponding loss of CDH1, consistent with a partial shift toward a more N-cadherin-expressing state. Based on these findings, subsequent hormonal-stimulation experiments focused on Matrigel and human mammary dECM, which represented the most distinct phenotypes: alveolar-like spherical organoids versus rapid, lineage-preserving branching morphogenesis.

### ECM composition differentially regulates lactation-associated states in milk MEC organoids

Milk MEC organoids were cultured in human mammary dECM and Matrigel and stimulated with 800 ng/mL PRL to evaluate differences in lactation-associated function. To exclude matrix-related variation in hormone transport, rhodamine-labeled PRL (23 kDa) diffusion was assessed in acellular crosslinked Matrigel and dECM droplets. PRL penetration was comparable in both matrices, indicating similar hormonal access to embedded cells (Fig. S27). Imaging showed lactation-associated markers in both matrices, such as milk fat globules (MFG) and β-casein, but signal intensity varied substantially between donors and conditions. Although signals appeared more consistent in Matrigel than in human dECM, no clear PRL-dependent increase was evident from imaging alone (Fig. 6A-B and Fig. S28).

**Figure 6.**
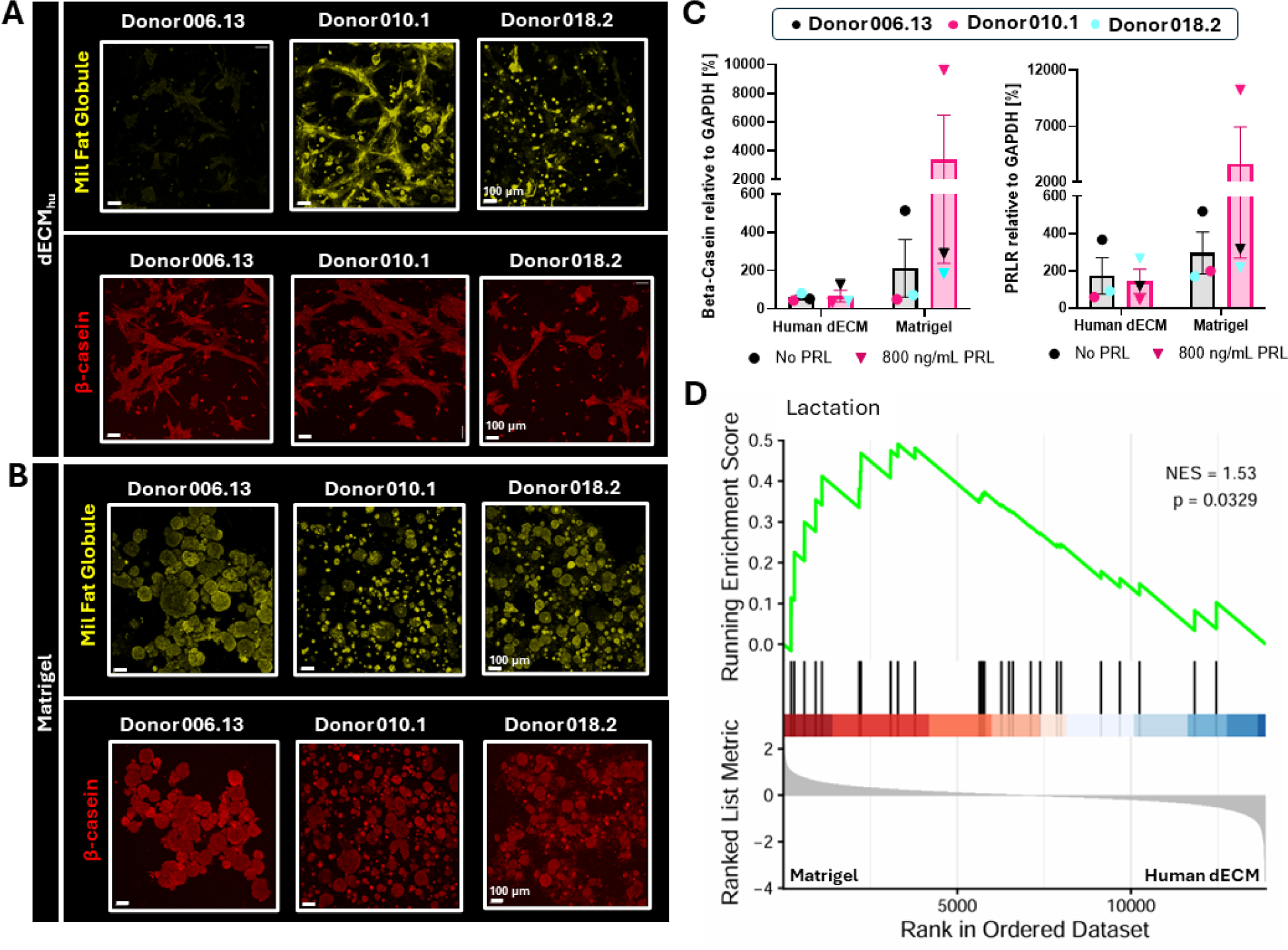
Human dECM and Matrigel differentially support lactation-associated marker expression in mammary organoids. **A-B)** Representative fluorescence images of milk MEC cultures embedded in human dECM **(A)** or Matrigel **(B)** showing milk fat globules (MFG; yellow) and β-casein (red); boxed insets show higher-magnification views. Scale bars, 100 µm**. C)** Quantification of β-casein (left) and PRLR (right) protein in primary milk mammary epithelial cells (milk MECs) from the indicated donor samples (006.13, 010.1, 018.2) cultured in human dECM or Matrigel ± 800 ng/mL PRL, normalized to GAPDH; colored symbols denote individual donors. **D)** Gene set enrichment analysis (GSEA) of the lactation pathway (GO:0007595) shows positive enrichment in Matrigel conditions, with leading-edge genes concentrated toward the top of the ranked gene list, indicating coordinated upregulation of lactation-associated genes (NES = 1.53, p = 0.0329, n = 3 donors).

To complement the imaging data, β-casein and PRLR protein expression were quantified by Western blot in organoids cultured in human dECM or Matrigel, with or without 800 ng/mL PRL. In milk MEC organoids, both markers were detectable even without exogenous PRL, but no consistent PRL-dependent increase was observed across samples (Fig. 6C and Fig. S29A). β-casein and PRLR levels trended higher in Matrigel than in human dECM, although these differences did not reach statistical significance in milk MECs. To assess matrix effects in a less variable epithelial model, the same analysis was performed in MCF10A organoids, where Matrigel supported significantly higher expression of lactation-associated markers than human dECM under the same conditions (Fig. S29B).

Next, bulk RNA barcoding and sequencing (BRB-seq) was performed on luminal/progenitor-like donors (n = 3) to determine whether ECM-dependent differences in lactation-related function could further be observed at the transcriptional level (Fig. S30A). Gene set enrichment analysis identified a significant lactation-associated signature in Matrigel relative to human dECM (Fig. 6D). Broader gene ontology analysis further showed that Matrigel and human dECM drove distinct biological programs, with Matrigel favoring lactation- and cell junction-related pathways, whereas human dECM was associated with estrogen receptor-related and cell cycle processes (Fig. S30B, C).

Despite PRL stimulation, milk MEC organoids in dECM or Matrigel did not exhibit PRL-induced morphological changes, and rounded alveolar-like structures at branch tips were not observed. To test whether organoid crowding and shared matrix remodeling obscured localized morphogenesis, PRL stimulation was repeated under reduced seeding density. Milk MEC were seeded at 200 cells per droplet and maintained for 14 days to preserve more isolated microenvironments compared with high-density cultures. Under these conditions, organoids adopted a more dispersed, stellate morphology, and rounded TDLU-like end buds remained absent (Fig. S31).

Together, these results show that matrix context supports distinct milk MEC states, with Matrigel favoring a more lactation-associated program and human dECM preserving a distinct matrix-dependent phenotype, that was not resolved into *in vivo*-like alveolar end buds by PRL stimulation alone.

## Discussion

Mammary tissue development and lactation emerge through dynamic interactions with the surrounding extracellular matrix, however these relationships remain difficult to model *in vitro* (23). In this study, primary human milk MEC organoids consistently assembled into complex 3D structures and remained responsive to matrix context. Rather than serving as a passive scaffold, the ECM biased milk MEC behavior toward distinct outcomes. Fibrillar collagen rich matrices supported branch formation together with collagen remodeling, whereas basement membrane-rich Matrigel favored a more lactation-associated state. Together, these findings show that ECM composition shapes how human milk MECs balance morphogenesis and functional differentiation *in vitro*, and highlight breast milk as a powerful noninvasive source of human mammary epithelium for mechanistic studies.

Milk MECs were expanded in 2D and 3D from unsorted milk cell populations, which were isolated based on centrifugation. Milk MECs readily formed monolayers and organoids, but TDLU-like structures in 3D (in 6 of 11 samples) were a transient phenotype, detectable only in the post-isolation culture (P0), and were not maintained with passaging under standard Matrigel conditions. Similar TDLU-like budding morphologies have also been reported for primary reduction-mammoplasty MECs [37, 38]. In these studies, cells were cultured directly from freshly isolated tissue-derived epithelial clusters (or, in some cases, single cells) and were evaluated at early time points without passaging[42]. Serial passaging can disrupt multicellular geometry and bias cultures toward acinar architectures over more complex morphological features [37, 43]. In contrast to breast tissue biopsies, human milk contains only a small epithelial fraction at isolation (∼11%), which is obtained as single cells rather than clusters [26, 27]. This creates a practical trade-off for milk as a MEC source, between retaining early-passage TDLU-formation potential and the expansion of sufficient cell numbers for mammary tissue engineering or comparative *in vitro* studies.

The breast is a structurally and compositionally complex organ in which epithelial form and function are regulated by a specialized microenvironment. The loss of *in vivo*-like structures *in vitro* during passaging could therefore also reflect progressive loss or dilution of native stromal niche signals. To better define the breast ECM microenvironment, we performed proteomic analysis on decellularized human and bovine mammary tissue. Human breast tissue provided the most relevant species context, whereas lactating bovine tissue enabled analysis of a lactation-state ECM that is rarely accessible in humans. Bovine tissue was also chosen for its accessibility and biological relevance, as its collagen-rich stromal architecture more closely resembles human breast than the predominantly adipose-rich rodent mammary gland [44]. Because our proteomic analysis is confounded by both species and physiological state (lactating bovine udder versus non-lactating adult human breast), any observed differences must be interpreted as reflecting the combined effects of species and physiological stage.

Global matrisome profiles of the human breast and lactating bovine udder dECM segregated robustly, and tissue-derived ECM hydrogels retained reproducible, source-specific biochemical signatures after decellularization. Both mammary ECMs were dominated by (fibrillar) collagens in line with previous studies which likely reflects conserved structural requirements between species and tissue state [45]. At the same time, human breast dECM was relatively enriched in collagen I, whereas bovine udder dECM contained a higher proportion of collagen II. Further, the observed relative enrichment of basement membrane-associated and ECM glycoprotein constituents in lactating udder dECM could reflect lactation state-dependent redistribution among ECM sub-compartments and may contribute to the higher proportion of spherical organoids in bovine compared to human dECM. Laminins and other basement membrane-associated proteins emerged among the strongest discriminators between matrices, which suggested that the differences were not limited to overall protein abundance but also involved distinct representation of specific ECM sub-compartments. These observations are supported by stage-resolved rodent mammary ECM proteomics from the Schedin laboratory, which underscored extensive matrisome remodeling across the reproductive cycle, with reduced fibrillar collagen during pregnancy and lactation and pronounced compositional shifts during involution, while basement membrane proteins remain a comparatively minor ECM fraction [45].

Mammary tissue-derived dECM hydrogels were generated to evaluate whether native stromal matrix cues could preserve or recover complex epithelial morphologies *in vitro*. Milk MECs adopted matrix-dependent morphologies and cell states. Basement membrane-rich Matrigel favored polarized, acinar-like structures. By contrast, in fibrillar matrices, dECM drove more extensive branching than purified collagen I across milk MEC and the MCF10A control. Matrix composition also shaped epithelial cell identity. In collagen I, branched organoids showed a shift away from luminal epithelial identity, with increased CDH2/N-cadherin expression, particularly in MCF10A, and reduced luminal-marker signal (CK8)[35, 46]. By contrast, dECM supported branching while preserving luminal features [35]. These findings are consistent with prior studies showing that fibrillar collagen can promote N-cadherin-associated migratory or EMT-like programs in mammary epithelial cells, whereas tissue-matched ECM better sustains epithelial differentiation and function. Collectively, these data indicate that dECM supports branch morphogenesis while maintaining luminal epithelial identity, and further demonstrate that milk MECs remain functionally responsive to ECM composition at the levels of tissue architecture and lineage-associated state.

Although dECM matrices robustly promoted branch formation, they did not restore budding in later passages, and additional tip-localized cues are required to generate and maintain TDLU morphogenesis. *In vivo*, formation of budded alveolar structures integrates immune and stromal cell support, hormones, and morphogen gradients [9]. Future experiments should therefore test whether adding other developmental cues, particularly hormonal signals beyond PRL such as estrogen and progesterone, can help establish budded tips for lactation studies [20].

In Matrigel, the expression of β-casein and PRLR in the absence of exogenous PRL further suggests that milk MEC organoids retain an intrinsic lactation baseline program which is gated by matrix context and is donor dependent. Donor-to-donor variability observed cell population distribution and hormonal response likely reflects biologically meaningful postpartum heterogeneity, such as variation in lactation stage, parity, endocrine milieu, metabolic status, and inflammatory exposures (e.g., subclinical mastitis)[27, 28]. Accordingly, donor-to-donor differences in lactation-associated signatures observed *in vitro* may reflect distinct functional states jointly shaped by ECM context and donor physiology. Thus, inter-individual variability is not merely a technical challenge, but an informative feature that may help explain differences in lactation performance across donors. Future work should link organoid phenotypes and *in vitro* lactation readouts to donor metadata and cell-state profiling by sequencing, to identify the molecular and physiological determinants that regulate lactational competence.

Our milk MEC-matrix platform provides a human-relevant framework for probing how postpartum perturbations reshape human mammary form and function. By enabling controlled interrogation of inflammatory, metabolic, and pharmacologic exposures within a physiologically informed ECM context, we provide a foundation for mechanistic studies of human lactation and branching that are difficult to address *in vivo* [47]. More broadly, ECM context is positioned as a central regulator of lactation biology and milk MECs provide a minimally invasive route toward defining cell-ECM mechanisms that govern milk supply robustness or mastitis susceptibility.

## Materials and Methods

### Milk-derived human mammary epithelial cells (MECs)

Human donors were recruited in line with Swiss Ethics guidelines, and written informed consent was obtained from all participants (Kantonale Ethikkomission, 2022-02012). Milk samples were obtained through recruitment conducted at ETH Zurich and through collaborating clinicians at University Hospital Zurich (USZ). Milk was collected by breast pump, transported on ice, diluted 1:1 in cold PBS, and centrifuged (780 RCF, 20 min, 4°C). The fat layer/supernatant was removed, and the pellet resuspended in 5 mL cold PBS, filtered (40 µm; VWR, 732-2757), and the original tube rinsed with an additional 5 mL PBS and filtered; filtrates were combined. Antibiotic–antimycotic (Gibco, 15249-062) was added to 2% (v/v) and cells washed by centrifugation (480 RCF, 5 min, room temperature) for three total wash cycles. Cells were resuspended in MEC isolation medium, counted on a Luna FX7 (Logos) using acridine orange/propidium iodide (F23001; cells:stain 9:1, v/v) (Table S1). Cultureware was coated with Matrigel diluted 1:26 (v/v) in cold MEC maintenance medium (Table S2) and polymerized at 37°C; cells were seeded at ≥120,000 cells/cm² and cultured at 37°C under hypoxia (5% CO₂, 5% O₂) for 5 days in MEC isolation medium to promote attachment, then switched to MEC expansion medium exchanged every 2 days for 12–15 days. Cells were passaged with TrypLE (Gibco, 12604021) at 37°C under hypoxia and cryopreserved or reseeded. Post-expansion, MECs were seeded at ≥8,000 cells/cm², maintained in MEC recovery medium (2 days), then in MEC maintenance medium exchanged every 2 days until >70% confluency (typically 7– 10 days) (Table S2).

### MCF10A

MCF10A cells (gift from the Bodenmiller group, ETH Zurich) were cultured on uncoated tissue-culture plastic at ≥4,000 cells/cm² in MCF10A medium at 37°C under normoxia (5% CO₂, 21% O₂) with medium exchange every 2 days; cells were counted and passaged using 0.25% trypsin-EDTA (Gibco, 25200056) (Table S2).

### Organoid culture

Milk MECs or MCF10A cells were pelleted (500 RCF, 5 min), resuspended in ice-cold hydrogel pre-gel, and mixed by pipetting; 30-µL droplets were dispensed into 24-well suspension plates (Greiner Bio-One, 662102), inverted, and polymerized (37°C, 45 min) under hypoxia (milk MEC) or normoxia (MCF10A). Droplets were overlaid with 1 mL MEC maintenance medium (milk MEC) or 500 µL MCF10A medium (MCF10A) and fed every 2 days. Prolactin stimulation used recombinant human prolactin (Thermo Fisher, 100-07; 800 ng/mL) added at each medium exchange. Unless noted, milk MEC organoids were cultured 7 days (hypoxia) and MCF10A organoids 5 days (normoxia); milk MEC passages 0–4 and MCF10A passages 10–18 were used. Seeding densities (cells/droplet) were: milk MEC donor samples 006.13, 7,000; donors 010.1/010.2, 10,000; donor 018.2, 10,000; MCF10A, 5,000 (Table S3).

### Morphology quantification

Morphology was quantified in Fiji (ImageJ) by area-based image analysis. Because branched epithelial structures were often interconnected, morphology was not assessed by counting individual organoids. Instead, confocal images were converted to 8-bit, manually thresholded, and used to determine total epithelial area. Spherical structures were identified using Fiji’s Analyze Particles function with defined circularity and size parameters (circularity, 0.4–1.0; size, 1900 µm² to infinity), branched/non-spherical area was calculated by subtraction from the total epithelial area. Data are reported as the percentage of total epithelial area occupied by spherical versus branched/non-spherical structures.

### Decellularized ECM (dECM) hydrogels

Ethics approval for the study was obtained (Kantonale Ethikkomission, 2022-01844), and human tissue samples were collected after written informed consent. Lyophilized human breast and lactating bovine udder dECM powders were prepared as described previously [26]. Powders were digested in pepsin (1 mg/mL; Sigma-Aldrich, P7012) in 0.1 M HCl (150 rpm, 48 h). Human dECM was digested at 10 mg/mL and bovine dECM at 20 mg/mL to obtain hydrogels of matched stiffness. Digests were neutralized on ice by gradual addition of 1 M NaOH (10% of final gel volume) under stirring (300 rpm), then adjusted to pH 7.0–7.4 with cold NaOH/HCl (pH paper). Neutralized solutions were supplemented with cold 10× DMEM/F-12 (1/9 final volume). Human dECM was clarified by centrifugation (3000 RCF, 10 min) and bovine dECM by centrifugation (10,000 RCF, 8 min); supernatants were collected. dECM pre-gels were stored at −20°C (short-term) or −80°C (long-term). Immediately before material testing/cell culture, human dECM was diluted 1:5 (v/v) with the appropriate culture medium to achieve the desired stiffness.

### Commercial hydrogels

Matrigel basement membrane matrix (Corning, 354234) was aliquoted upon receipt and stored at −20°C (short-term). Rat tail collagen I (4.79 mg/mL; Merck, 08-115) was neutralized on ice and diluted to 2.6 mg/mL by adding 0.5 M NaOH (1:200, v/v), then adding cold 10× DMEM/F-12 stepwise in three equal portions with mixing between additions to ensure uniform neutralization (color uniformity); pH was adjusted to 7.0–7.4 with cold NaOH/HCl (pH paper). Neutralized collagen pre-gels were aliquoted and stored at −20°C (short-term).

### Rheology

Rheology was performed on an Anton Paar MCR 302e (20-mm parallel plates, 0.2-mm gap). Pre-gels (76 µL) were loaded onto a pre-cooled (4°C) plate and evaporation was minimized with a damp tissue. Gelation kinetics and storage modulus (G′) were measured by oscillatory time sweeps (γ₀ = 1%, ω = 6 rad/s; sampling every 10 s): samples were equilibrated at 4°C for 10 min, heated to 37°C, and recorded for ∼1 h (n = 4 per material). Shear response was measured by time sweeps at 2% strain and 1 Hz at 25°C for 20 min and 37°C for 25 min, followed by a 37°C strain sweep (0–15% at 1 Hz over 45 min; sampling every 2 s); shear stress–time curves were obtained from the 37°C strain-sweep data (n = 3 per material).

### Compression

Compression testing used a TA.XT Texture Analyzer (Stable Micro Systems) with a 500 g load cell. Gels were cast by polymerizing 56 µL pre-gel in PDMS rings (Sylgard 184, Corning, 401986) to yield cylinders (6 mm diameter, 2 mm thickness) and incubated at 37°C for 45 min. Gels were removed, placed between plates, preloaded to 0.1 g, and compressed with a 15-mm cylindrical probe to 15% strain at 0.01 mm/s. Compressive modulus was obtained by linear fitting of the first 3% of the stress–strain curve (n = 4 per material).

### Second harmonic generation (SHG) imaging

SHG imaging used a Leica SP8 with a Mai Tai laser (Spectra-Physics) (excitation 900 nm; SHG collection 420 nm) and a 25× water objective (NA 1.05). Hydrogels (300 µL) were polymerized at 37°C for 1 h before imaging. Organoid-laden hydrogels were fixed in 4% PFA and stained with DAPI or phalloidin-iFluor 555 as above, then imaged by SHG on the Leica SP8 (900-nm excitation, 420-nm SHG collection) using a 40× water objective (NA 1.1), acquiring SHG and fluorescence sequentially.

### Scanning electron microscopy (SEM)

Hydrogels (300 µL) were polymerized at 37°C for 1 h, fixed in 4% PFA (Carl Roth, P087.1) for 4 h at room temperature, and washed three times in PBS (Gibco, 10010-015). Samples were dehydrated in graded ethanol (20–100%; 10–16 h per step), then graded hexamethyldisilazane (HMDS; abcam, AB111175) in ethanol (20–100%; 1 h per step), and air-dried overnight. Samples were mounted with silver paint (Plano, G3692), carbon-coated (10 nm; Safematic CCU-010), and imaged on a Zeiss Merlin FE-SEM at 2.0 kV using an in-lens secondary electron detector (working distance ∼5–10 mm).

### Histology

Tissue and hydrogel droplets were fixed in 4% PFA for 20 min, washed three times in PBS, paraffin-embedded, sectioned at 10 µm, mounted (Eukitt; Sigma-Aldrich, 03989), and scanned on an Olympus Slideview VS200. Collagen was visualized by Picrosirius Red (Weigert’s iron hematoxylin 8 min; Sirius Red F3B/Direct Red 80, 0.1% w/v in picric acid 1 h; rinse in acidified water; ethanol dehydration; xylene clearing). Glycosaminoglycans were visualized by Safranin O/Fast Green (Weigert’s 5 min; differentiate in 1% acid alcohol; Fast Green 0.02% w/v 1 min; Safranin O 1% w/v 30 min; ethanol dehydration; xylene clearing). Elastic fibers were stained using Verhoeff’s Elastic Stain Kit (Sigma-Aldrich, HT25A-1KT) per manufacturer instructions (Table S4).

### Immunostaining and confocal microscopy

At endpoint, droplets were washed once in PBS. Collagen I/dECM organoids were fixed in-gel; Matrigel organoids were released as below, pelleted (500 RCF, 5 min), and handled with FBS-coated tips. All samples were fixed in 4% PFA (20 min, room temperature), washed in PBS (once for dissociated organoids; three times for in-gel organoids), blocked/permeabilized (blocking solution; 20 min), and incubated overnight at 4°C with primary antibodies in staining solution. Samples were washed (once or three times) and incubated 2–3 h at room temperature with secondary antibodies, DAPI (2:500; Thermo Fisher, H1399), and where indicated phalloidin-iFluor 555 (1:500; Abcam, ab176756) or phalloidin-iFluor 647 (1:500; Abcam, ab176759), then washed and imaged in PBS. Images were acquired on an Olympus FluoView4000 using 10× (NA ∼0.40) or 20× (NA ∼0.75) air objectives with 405/488/561/640-nm lasers; only linear brightness/contrast adjustments were applied. For β-casein and milk fat globule imaging, acquisition settings and post-processing were held constant across conditions/donors (Tables S4-S6).

### Image analysis

Morphology (spherical vs branched) was quantified in Fiji from merged 10× images (DAPI/CK8/CK14 or DAPI/F-actin/E-cadherin/Ki-67): images were converted to 8-bit, thresholded manually, and total organoid area measured. Spheres were identified by Analyze Particles using ≥50 µm² area and 0.40–1.00 circularity thresholds; branched area was computed as total minus spherical area and used to calculate spherical-to-branched area ratio (3 images/condition, averaged). Ki-67 was quantified in Fiji from three 20× images/condition: for thin stacks, maximum projections were used; for thick stacks, three non-overlapping optical slices were quantified and summed per replicate. DAPI nuclei were segmented by manual thresholding with watershed and corrected manually when required; Ki-67⁺ nuclei were defined by colocalization with DAPI using an AND operation. Particle analysis used ≥50 µm² (DAPI) and ≥35 µm² (Ki-67) size thresholds and 0.25–1.00 circularity; %Ki-67⁺ = (Ki-67⁺/DAPI⁺)×100 and averaged across replicates. Actin alignment was quantified with OrientationJ: for each matrix, 10 branches (collagen I/dECM) or spheres (Matrigel) were cropped from three replicate 20× maximum projections and analyzed using OrientationJ Distribution/Analysis; alignment was reported as the fraction of actin signal within ±15° of the dominant orientation (10 structures/condition, averaged).

### Matrix dissociation

For Matrigel imaging, RT-qPCR, and western blot, organoids were released enzymatically: Matrigel droplets were incubated in Dispase II (10 mg/mL in PBS; Sigma-Aldrich, D4693) for 20 min at room temperature and dissociated by gentle pipetting; collagen I/dECM droplets were incubated in collagenase (500 U/mL; Sigma-Aldrich, C9407) in PBS + 10 mM CaCl₂ for 15 min at 37°C and dissociated by gentle pipetting. Organoids were pelleted (500 RCF, 5 min), washed once in PBS, and fixed or stored at −80°C.

### Western blot

Milk MEC organoids were lysed in 100 µL RIPA buffer + protease inhibitors (1:1000; Sigma-Aldrich, P1860) (15 min on ice), then clarified (12,000 RCF, 10 min). Samples (16.25 µL lysate) were mixed with NuPAGE reducing agent (2.5 µL; Thermo Fisher, NP0009) and LDS sample buffer (2.5 µL; Thermo Fisher, NP0007), denatured (10 min, 80°C), and 20 µL loaded on NuPAGE 4–12% Bis-Tris gels (Thermo Fisher, NP0321BOX) with MOPS running buffer (200 V, ∼1 h). Proteins were transferred to 0.2-µm nitrocellulose (Amersham, 10600015) (25 V, ∼1 h) in transfer buffer, washed in PBST, blocked (5% milk in PBST, 30 min), incubated with primary antibodies overnight at 4°C, washed three times (10 min), incubated with HRP secondaries in 5% milk/PBST (1.5 h), washed three times (10 min), developed with WesternBright ECL (Advansta, K-12045-D20), and imaged on a FUSION FX6 EDGE system. Band intensities were quantified in ImageJ and normalized to GAPDH (Table S7, S8).

### RNA extraction and RT-qPCR

For each matrix, MCF10A and milk MEC donor sample 006.13 organoids were cultured as three biological replicates (each: pooled triplicate droplets). After dissociation, pellets were lysed in 250 µL NucleoZOL (Macherey-Nagel, 740404.200), mixed with 100 µL RNase-free water (Promega, P119E) (5 min, room temperature), centrifuged (12,000 RCF, 15 min), and 250 µL supernatant was precipitated with an equal volume of isopropanol (10 min, room temperature; 12,000 RCF, 10 min). Pellets were washed three times with 75% ethanol (8,000 RCF, 3 min) and resuspended in 15 µL RNase-free water; RNA was assessed by NanoDrop OneC. Reverse transcription used GoScript (Promega, A5003) with 1 µg RNA (MCF10A) or 100 ng (milk MEC); cDNA was diluted 1:5 (v/v). RT-qPCR used GoTaq qPCR Master Mix (Promega, A6002) on a QuantStudio 3 (Applied Biosystems) with two technical replicates per biological replicate; Ct values were normalized to GAPDH and reported as mean of three biological replicates; primer sequences are provided in the Supplementary Materials (Table S9).

### Biochemical assays (DNA, collagen, sGAG, elastin)

Residual DNA in randomly selected decellularized tissue pieces was quantified using the PureLink Genomic DNA Mini Kit (Invitrogen, K182001) with NanoDrop readout. Collagen, sulfated glycosaminoglycans (sGAGs), and elastin were measured using the QuickZyme Collagen assay (QuickZyme Biosciences), Blyscan sGAG assay (Biocolor, B3000), and Elastin assay (Biocolor, F4000), respectively, following the manufacturers’ protocols.

### Proteomics sample preparation

Lyophilized bovine dECM from n = 3 pooled batches (4 lactating udders/batch) and human breast tissue (n= 3 biological replicates) were fractionated into soluble and insoluble ECM. For each batch, 2 mg material was bead-homogenized (100 mg 3-mm glass beads) in 6 M guanidine-HCl, and 100 mM ammonium bicarbonate (200 µL/mg) (Bullet Blender BBX24; power 8, 1 min), vortexed overnight at 25°C, and centrifuged (18,000g, 15 min, 4°C) to collect the soluble fraction. Pellets were extracted with hydroxylamine buffer (1 M NH₂OH-HCl, 4.5 M guanidine-HCl, 0.2 M K₂CO₃; pH 9.0) at 200 µL/mg, homogenized (power 8, 1 min), incubated (45°C, 1000 rpm, 4 h), centrifuged (18,000g, 15 min), and the insoluble fraction was stored at −80°C. Both fractions were digested with trypsin (1:100 enzyme:protein, overnight) using filter-aided sample preparation and desalted during Evotip loading.

### LC–MS/MS

Tryptic peptides were loaded onto Evotips and analyzed on an Evosep One using the default “30 samples/day” method with a Pepsep column (150 µm ID × 15 cm; ReproSil C18, 1.9 µm, 120 Å) coupled to a timsTOF Pro (Bruker) via Captive Spray. Data were acquired in PASEF mode (ramp 100 ms; 10 PASEF MS/MS scans per TopN cycle; m/z 100–1700; ion mobility 0.7– 1.50 Vs/cm²), with precursor isolation ±1 Th and ion mobility–dependent collision energy (20–59 eV). Precursors >500 counts were considered, with resequencing low-abundance ions (<20,000 target) and dynamic exclusion of 0.4 min.

### Database search and label-free quantification

Raw files were searched with MSFragger via FragPipe v21.1 against UniProt Bos taurus plus contaminants (37,606 sequences) for bovine samples and SwissProt *Homo sapiens* plus contaminants (20,466 sequences) for human samples using ±15 ppm precursor and ±0.08 Da fragment tolerances, semispecific trypsin, and 1% FDR at peptide and protein levels. Carbamidomethyl (C) was fixed; oxidation (M), oxidation/hydroxylation (P), dioxidation (P), deamidation (N/Q), N-terminal pyroglutamate (Q), and peptide N-terminal acetylation were variable. Label-free quantification used IonQuant v1.10.12 with match-between-runs enabled; soluble and insoluble fractions were searched separately and merged post-search. Human and bovine datasets were merged based on gene name.

### Proteomics data analysis

Proteomic data analysis was performed in Excel and proteomics intensity tables were analyzed in MetaboAnalyst (https://www.metaboanalyst.ca). Data were log-transformed and sum-normalized, then used for PCA and PLS-DA analyses. Differential abundance was assessed by univariate testing (fold change from group means), and results were visualized with volcano plots using the thresholds indicated in the figure legends.

### Flow cytometry staining and analysis

Single-cell suspensions were blocked with Fc receptor–blocking reagent (Human BD Fc Block, 1:50; BD Pharmingen, 564219) for 10 min at room temperature, followed by staining with the antibody panel (Table S10) prepared in FACS buffer (PBS, 2% fetal bovine serum, 0.5 mM EDTA) for 30 min at 4°C. Cell viability was assessed using Zombie NIR Fixable Viability dye (1:1000; BioLegend, 423105) in PBS. After staining, cells were fixed with 4% paraformaldehyde for 10 min, washed, and resuspended in FACS buffer for acquisition on a BD LSRFortessa (BD FACSDiva v8.0.1). Flow cytometry data were processed in FlowJo v10. Gating hierarchies are described in the Supplementary Materials (Fig. S21)

### Bulk RNA Barcoding and sequencing

BRB-seq count data were aligned to the human reference genome (hg38) using STAR, after which gene-level count matrices were imported into R for downstream analysis. Sample metadata were annotated by donor and culture condition, and lowly expressed genes were filtered prior to normalization. Variance-stabilizing transformation was performed with DESeq2 for unsupervised analyses. For lineage- and state-associated analyses, curated luminal, basal, progenitor, mammary epithelial, estrogen receptor target, and lactation-related gene sets were evaluated using gene-wise z-scored expression values and summarized at the sample level. For differential expression between Matrigel and Human dECM in luminal/progenitor donors, raw counts were analyzed using the limma-voom workflow following TMM normalization in edgeR, with donor included in the design matrix to account for paired donor structure. Effect-size-aware differential expression testing was additionally performed using limma-TREAT with a log2 fold-change threshold of 0.5. Gene set enrichment analysis was performed on ranked differential expression results using clusterProfiler with MSigDB-derived GO Biological Process gene sets. Visualizations were generated using ggplot2.

## Acknowledgements

We thank the participating mothers and their children, without whom this research would not have been possible. We acknowledge the medical staff at University Hospital Zürich for assistance with consent and sample collection. We thank Prof. H. Gehart and members of his laboratory, S. Moser, M. Winkelbauer, Prof. S. Kelleher, Dr. E.-H. Ervin, and members of the TEB laboratory for helpful discussions and input. We thank G. Silvestrelli and S. Ulbrich for helping with the procurement of bovine tissue and for sharing reagents. We thank C. Bigosch for the help with the ethics approval. We acknowledge ScopeM for support and assistance. Lightsheet imaging was performed with equipment maintained by the Center for Microscopy and Image Analysis, University of Zurich. We acknowledge the Flow Cytometry Core Facility at ETH Zurich for support, with special thanks to R. Antonialli and F. Mair. AI-assisted proofreading was used for linguistic refinement of the manuscript and all content was verified and approved by the authors.

## Ethics approval

Written informed consent was obtained from all milk donors prior to sample collection. The study involving human milk-derived mammary epithelial cells was approved by the Kantonale Ethikkommission (approval no. 2022-02012). Human breast tissue was collected after written informed consent under ethics approval from the Kantonale Ethikkommission (approval no. 2022-01844).

## Data and materials availability

All data needed to evaluate the conclusions in the paper are present in the paper and/or the Supplementary Materials. Additional data is available in the ETH Zurich Research Collection (https://doi.org/10.3929/ethz-c-000798071) under the terms of the repository’s data-sharing policies and sequencing data has been deposited in the NCBI Gene Expression Omnibus (GEO) under accession number GSE327550.

## References

1. Macias H, Hinck L (2012) Mammary Gland Development. Wiley Interdiscip Rev Dev Biol 1:533–557. 10.1002/wdev.35

2. Levental KR, Yu H, Kass L, et al (2009) Matrix Crosslinking Forces Tumor Progression by Enhancing Integrin Signaling. Cell 139:891–906. 10.1016/j.cell.2009.10.027

3. Yang H, Yang J, Zheng X, et al (2024) The Hippo Pathway in Breast Cancer: The Extracellular Matrix and Hypoxia. International Journal of Molecular Sciences 25:12868. 10.3390/ijms252312868

4. Lee S, Kelleher SL (2016) Biological underpinnings of breastfeeding challenges: the role of genetics, diet, and environment on lactation physiology. Am J Physiol Endocrinol Metab 311:E405–E422. 10.1152/ajpendo.00495.2015

5. Bjerre LB, Chalmers SB, Davis FM (2025) Little women: the relevance and reliance on mouse models for mammary gland research and next steps for translation. J Mammary Gland Biol Neoplasia 30:16. 10.1007/s10911-025-09590-8

6. Donovan SM, Aghaeepour N, Andres A, et al (2023) Evidence for human milk as a biological system and recommendations for study design—a report from “Breastmilk Ecology: Genesis of Infant Nutrition (BEGIN)” Working Group 4. The American Journal of Clinical Nutrition 117:S61–S86. 10.1016/j.ajcnut.2022.12.021

7. Rauner G, Jin DX, Miller DH, et al (2021) Breast tissue regeneration is driven by cell-matrix interactions coordinating multi-lineage stem cell differentiation through DDR1. Nat Commun 12:7116. 10.1038/s41467-021-27401-6

8. Weaver VM, Lelièvre S, Lakins JN, et al (2002) β4 integrin-dependent formation of polarized three-dimensional architecture confers resistance to apoptosis in normal and malignant mammary epithelium. Cancer Cell 2:205–216. 10.1016/S1535-6108(02)00125-3

9. Fata JE, Werb Z, Bissell MJ (2004) Regulation of mammary gland branching morphogenesis by the extracellular matrix and its remodeling enzymes. Breast Cancer Res 6:1–11. 10.1186/bcr634

10. Schedin P (2006) Pregnancy-associated breast cancer and metastasis. Nat Rev Cancer 6:281–291. 10.1038/nrc1839

11. Yu W, Datta A, Leroy P, et al (2005) Beta1-integrin orients epithelial polarity via Rac1 and laminin. Mol Biol Cell 16:433–445. 10.1091/mbc.e04-05-0435

12. Klinowska TCM, Soriano JV, Edwards GM, et al (1999) Laminin and β1 Integrins Are Crucial for Normal Mammary Gland Development in the Mouse. Developmental Biology 215:13–32. 10.1006/dbio.1999.9435

13. Nerger BA, Jaslove JM, Elashal HE, et al (2021) Local accumulation of extracellular matrix regulates global morphogenetic patterning in the developing mammary gland. Current Biology 31:1903–1917.e6. 10.1016/j.cub.2021.02.015

14. Gjorevski N, Nelson CM (2010) Endogenous patterns of mechanical stress are required for branching morphogenesis. Int Bio (Cam) 2:424–434. 10.1039/c0ib00040j

15. Varner VD, Nelson CM (2014) Cellular and physical mechanisms of branching morphogenesis. Development 141:2750–2759. 10.1242/dev.104794

16. Lyons TR, O’Brien J, Borges V, et al (2011) Postpartum mammary gland involution drives DCIS progression through collagen and COX-2. Nat Med 17:1109–1115. 10.1038/nm.2416

17. Simian M, Bissell MJ (2016) Organoids: A historical perspective of thinking in three dimensions. J Cell Biol 216:31–40. 10.1083/jcb.201610056

18. Yang J, Richards J, Bowman P, et al (1979) Sustained growth and three-dimensional organization of primary mammary tumor epithelial cells embedded in collagen gels. Proc Natl Acad Sci U S A 76:3401–3405. 10.1073/pnas.76.7.3401

19. Buchmann B, Engelbrecht LK, Fernandez P, et al (2021) Mechanical plasticity of collagen directs branch elongation in human mammary gland organoids. Nat Commun 12:2759. 10.1038/s41467-021-22988-2

20. Sumbal J, Chiche A, Charifou E, et al (2020) Primary Mammary Organoid Model of Lactation and Involution. Front Cell Dev Biol 8:. 10.3389/fcell.2020.00068

21. Yuan L, Xie S, Bai H, et al (2023) Reconstruction of dynamic mammary mini gland in vitro for normal physiology and oncogenesis. Nat Methods 20:2021–2033. 10.1038/s41592-023-02039-y

22. Dontu G, Ince TA (2015) Of Mice and Women: A Comparative Tissue Biology Perspective of Breast Stem Cells and Differentiation. J Mammary Gland Biol Neoplasia 20:51–62. 10.1007/s10911-015-9341-4

23. Caruso M, Saberiseyedabad K, Mourao L, Scheele CLGJ (2024) A Decision Tree to Guide Human and Mouse Mammary Organoid Model Selection. Methods Mol Biol 2764:77–105. 10.1007/978-1-0716-3674-9_7

24. Pavlovich AL, Manivannan S, Nelson CM (2010) Adipose Stroma Induces Branching Morphogenesis of Engineered Epithelial Tubules. Tissue Engineering Part A 16:3719–3726. 10.1089/ten.tea.2009.0836

25. Zhang Y, Tang C, Span PN, et al (2020) Polyisocyanide Hydrogels as a Tunable Platform for Mammary Gland Organoid Formation. Advanced Science 7:2001797. 10.1002/advs.202001797

26. Hasenauer A, Bevc K, McCabe MC, et al (2025) Volumetric printed biomimetic scaffolds support in vitro lactation of human milk-derived mammary epithelial cells. Sci Adv 11:eadu5793. 10.1126/sciadv.adu5793

27. Martin Carli JF, Trahan GD, Jones KL, et al (2020) Single Cell RNA Sequencing of Human Milk-Derived Cells Reveals Sub-Populations of Mammary Epithelial Cells with Molecular Signatures of Progenitor and Mature States: a Novel, Non-invasive Framework for Investigating Human Lactation Physiology. J Mammary Gland Biol Neoplasia 25:367–387. 10.1007/s10911-020-09466-z

28. Twigger A-J, Engelbrecht LK, Bach K, et al (2022) Transcriptional changes in the mammary gland during lactation revealed by single cell sequencing of cells from human milk. Nat Commun 13:562. 10.1038/s41467-021-27895-0

29. Hassiotou F, Beltran A, Chetwynd E, et al (2012) Breastmilk Is a Novel Source of Stem Cells with Multilineage Differentiation Potential. Stem Cells 30:2164–2174. 10.1002/stem.1188

30. Hasenauer A, Zenobi-Wong M (2026) A 3D printed model of human lactation. 2026.01.30.702762

31. Liu H, Gong Y, Zhang K, et al (2023) Recent Advances in Decellularized Matrix-Derived Materials for Bioink and 3D Bioprinting. Gels 9:195. 10.3390/gels9030195

32. Fernández-Pérez J, Ahearne M (2019) The impact of decellularization methods on extracellular matrix derived hydrogels. Sci Rep 9:14933. 10.1038/s41598-019-49575-2

33. Guo X, Liu B, Zhang Y, et al (2024) Decellularized extracellular matrix for organoid and engineered organ culture. J Tissue Eng 15:20417314241300386. 10.1177/20417314241300386

34. Gadre M, Kasturi M, Agarwal P, Vasanthan KS (2024) Decellularization and Their Significance for Tissue Regeneration in the Era of 3D Bioprinting. ACS Omega 9:7375–7392. 10.1021/acsomega.3c08930

35. Mollica PA, Booth-Creech EN, Reid JA, et al (2019) 3D bioprinted mammary organoids and tumoroids in human mammary derived ECM hydrogels. Acta Biomaterialia 95:201–213. 10.1016/j.actbio.2019.06.017

36. Blanco-Fernandez B, Rey-Vinolas S, Bağcı G, et al (2022) Bioprinting Decellularized Breast Tissue for the Development of Three-Dimensional Breast Cancer Models. ACS Appl Mater Interfaces 14:29467–29482. 10.1021/acsami.2c00920

37. Rosenbluth JM, Schackmann RCJ, Gray GK, et al (2020) Organoid cultures from normal and cancer-prone human breast tissues preserve complex epithelial lineages. Nat Commun 11:1711. 10.1038/s41467-020-15548-7

38. Sokol ES, Miller DH, Breggia A, et al (2016) Growth of human breast tissues from patient cells in 3D hydrogel scaffolds. Breast Cancer Res 18:19. 10.1186/s13058-016-0677-5

39. Qu Y, Han B, Yu Y, et al (2015) Evaluation of MCF10A as a Reliable Model for Normal Human Mammary Epithelial Cells. PLoS One 10:e0131285. 10.1371/journal.pone.0131285

40. Park K-S, Dubon MJ, Gumbiner BM (2015) N-cadherin mediates the migration of MCF-10A cells undergoing bone morphogenetic protein 4-mediated epithelial mesenchymal transition. Tumour Biol 36:3549–3556. 10.1007/s13277-014-2991-9

41. Jenkins Jr. EC, Debnath S, Varriano S, et al (2014) Na+/H+ exchanger 1 (NHE1) function is necessary for maintaining mammary tissue architecture. Developmental Dynamics 243:229–242. 10.1002/dvdy.24032

42. Linnemann JR, Miura H, Meixner LK, et al (2015) Quantification of regenerative potential in primary human mammary epithelial cells. Development 142:3239–3251. 10.1242/dev.123554

43. Paavolainen O, Peurla M, Koskinen LM, et al (2024) Volumetric analysis of the terminal ductal lobular unit architecture and cell phenotypes in the human breast. Cell Reports 43:114837. 10.1016/j.celrep.2024.114837

44. Hughes K (2021) Comparative mammary gland postnatal development and tumourigenesis in the sheep, cow, cat and rabbit: Exploring the menagerie. Seminars in Cell & Developmental Biology 114:186–195. 10.1016/j.semcdb.2020.09.010

45. Goddard ET, Hill RC, Barrett A, et al (2016) Quantitative extracellular matrix proteomics to study mammary and liver tissue microenvironments. Int J Biochem Cell Biol 81:223–232. 10.1016/j.biocel.2016.10.014

46. Brownfield DG, Venugopalan G, Lo A, et al (2013) Patterned Collagen Fibers Orient Branching Mammary Epithelium through Distinct Signaling Modules. Curr Biol 23:703–709. 10.1016/j.cub.2013.03.032

47. Majood M, Rao R (2025) Human milk: insights on cell composition, organoids and emerging applications. Pediatr Res 1–12. 10.1038/s41390-025-04458-3

