## Supplementary Material for "Extracellular matrix context shapes morphogenesis and lactation-associated states in human milk-derived mammary organoids"

Amelia Hasenauer *et al.*

*Marcy Zenobi-Wong,

**This PDF file includes:**

Figures S1 to S31

Tables S1 to S10

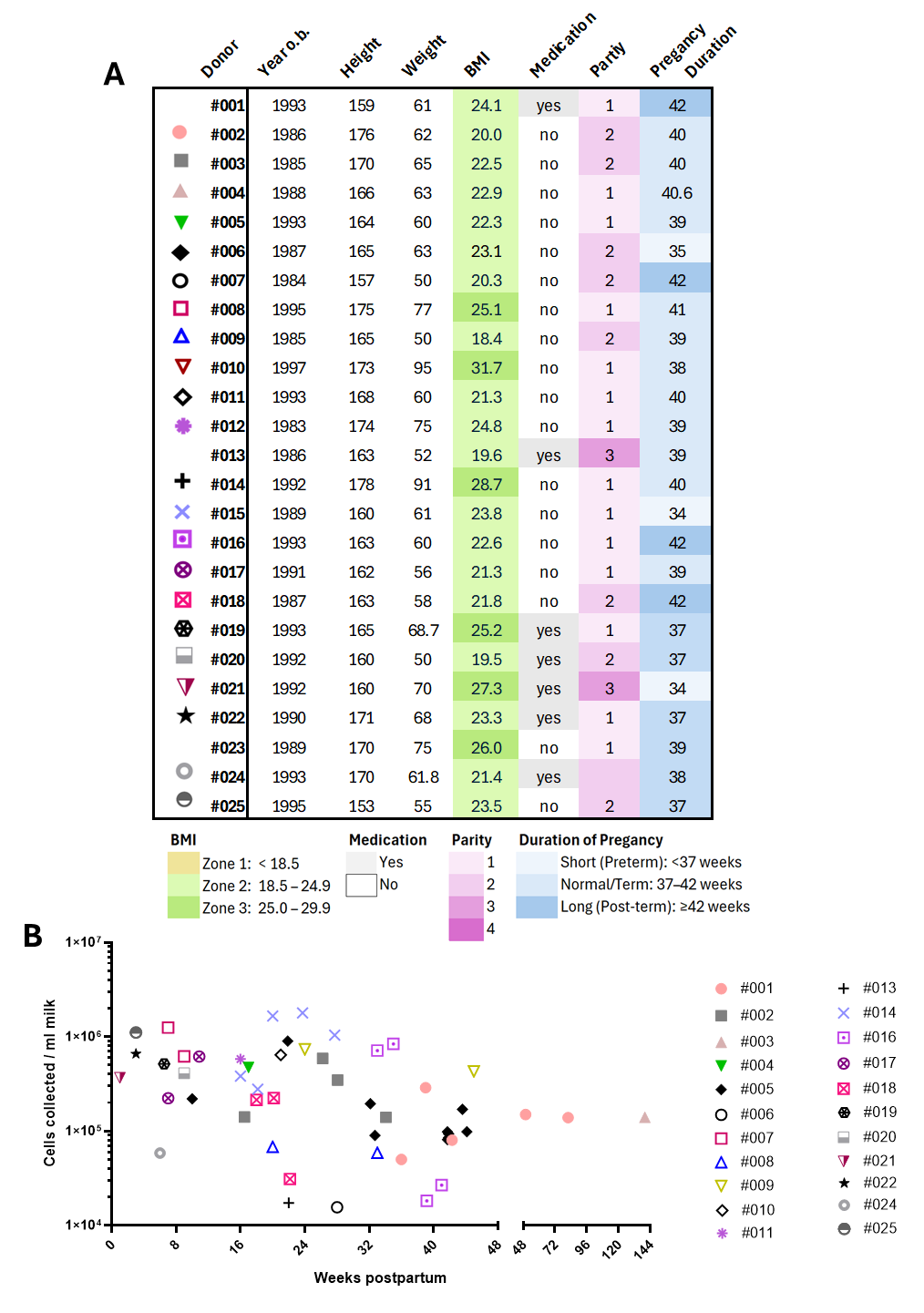

**Figure S1: Milk Donor characteristics**. Each row represents one donor; the leftmost symbol denotes the donor ID. Columns report year of birth, height (cm), weight (kg), and body mass index (BMI, kg/m²). BMI values are color-coded by zone: Zone 1, <18.5; Zone 2, 18.5–24.9; Zone 3, 25.0–29.9. Medication status is indicated as yes or no. Parity is encoded by a pink gradient (1–4). Pregnancy duration (weeks) is shown with a blue gradient and categorized as Short (preterm, <37 weeks), Normal/Term (37–42 weeks), and Long (post-term, ≥42 weeks). **B)** Cell yield from milk samples plotted against weeks postpartum; each point represents one milk sample, and symbol/color correspond to the donor number as in panel A.

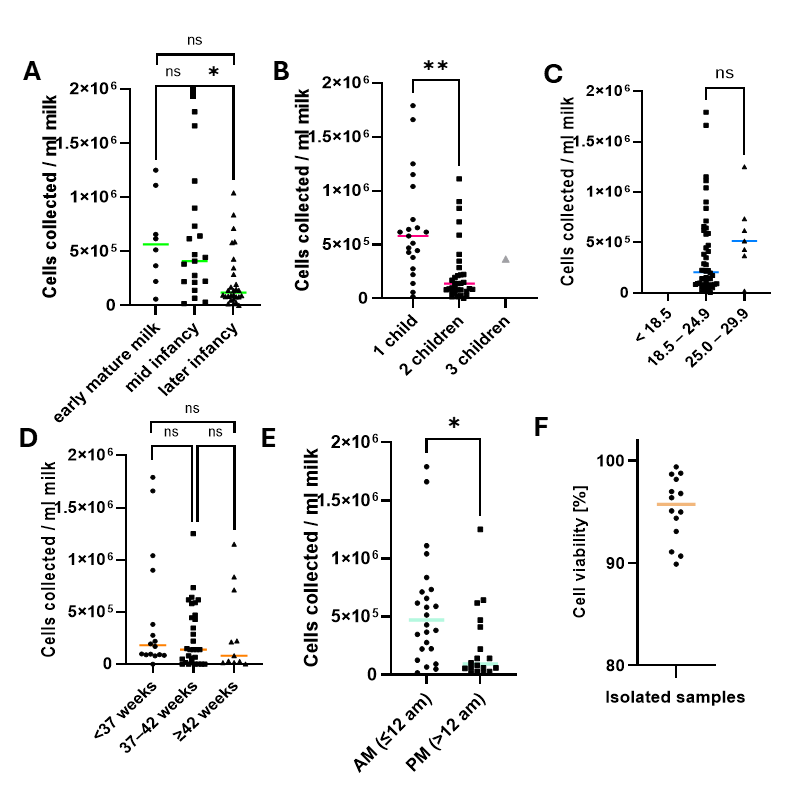

**Figure S2: Milk cell yield stratified by demographic and perinatal variables and viability post isolation. A–D)** Scatter plots show the number of cells collected per mL of milk for each sample (points), with the group mean indicated by the horizontal colored line. Statistical significance is annotated above comparisons (*p < 0.05; **p < 0.01; ns, not significant). **A)** Lactation stage: early mature milk (2–8 weeks; n = 8), mid infancy (9–24 weeks; n = 19), later infancy (≥25 weeks; n = 28). **B)** Parity (number of children): 1 child (n = 21), 2 children (n = 33), 3 children (n = 1, was not included in statistical analysis due to low sample size). **C)** Maternal BMI category: <18.5 (n = 0), 18.5–24.9 (n = 48), 25.0–29.9 (n = 7). **D)** Gestational age at delivery: <37 weeks (n = 8), 37–42 weeks (n = 19), >42 weeks (n = 28). **E)** Pumping time; morning (AM, n= 24) vs. afternoon (PM, n=18). **F)** Cell viability post isolation (n=14 samples), the viability is above 90%.

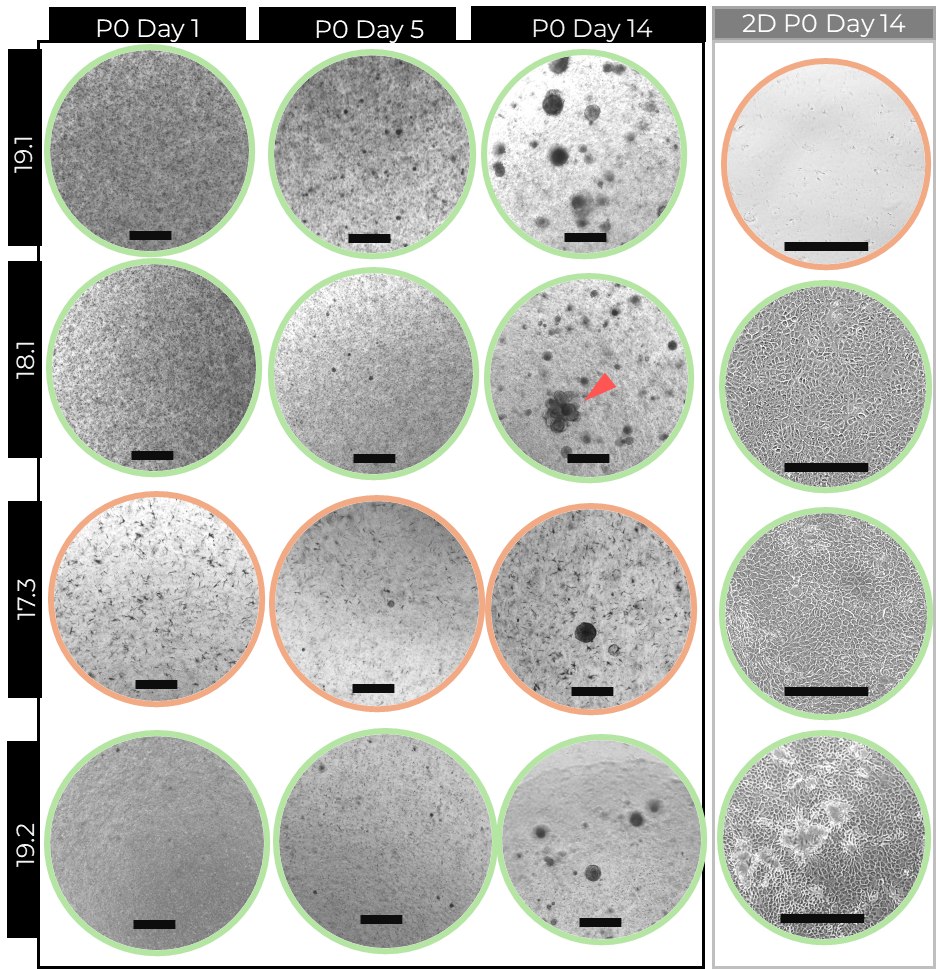

**Figure S3: Human milk MEC in culture.** Representative brightfield images of Passage 0 (P0) human milk–MECs from individual donor samples (19.1, 18.1, 17.3, 19.2, and 19.3) over time in culture. Images were acquired on Day 1, Day 5, and Day 14. The rightmost column shows the corresponding 2D culture on Day 14. Colored outlines indicate growth (green) or no growth (orange). Red arrowhead indicate branched organoid. Scale bars: 750 μm.

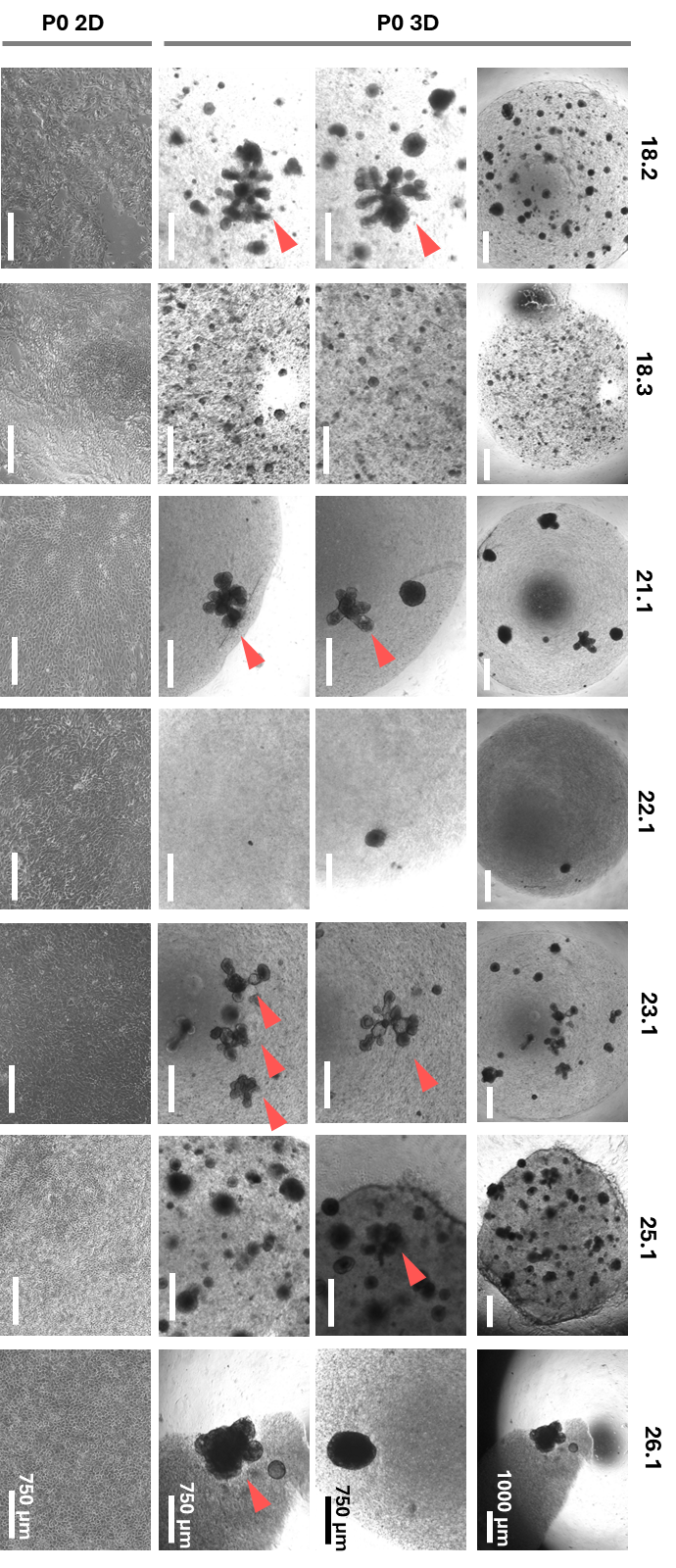

**Figure S4. Milk-derived MEC in 2D and 3D culture**. Representative brightfield images of mammary epithelial cells (MECs) isolated from milk and maintained at passage 0 (P0, post isolation) for 14–20 days in culture. Red arrowheads indicate branched organoids.

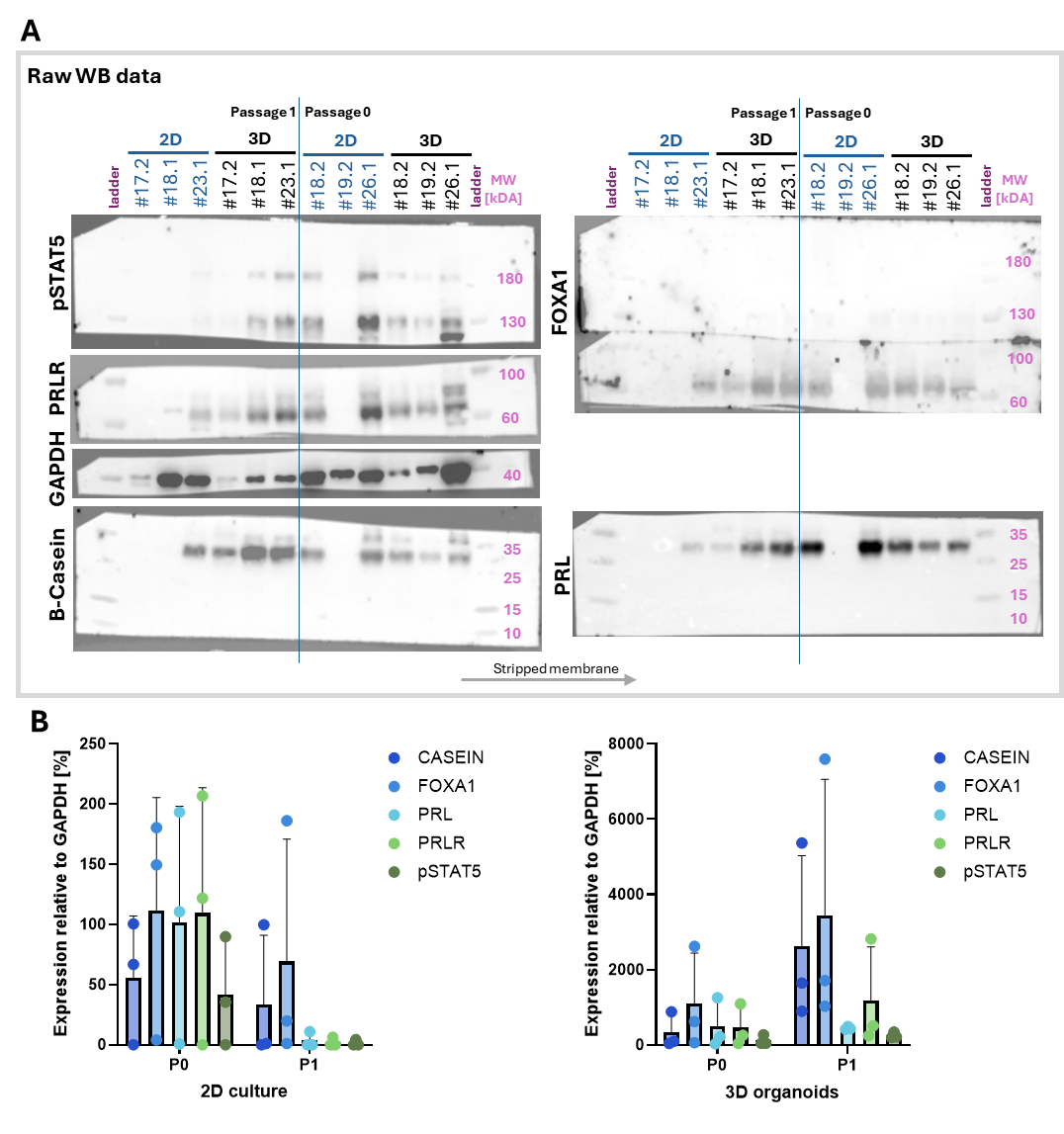

**Figure S5** **Western blot analysis of milk-derived MECs cultured in 2D and 3D at passage 0 and passage 1. (A)** Representative Western blots of milk-derived mammary epithelial cells (Milk MECs) cultured in 2D or 3D at passage 0 (P0) and passage 1 (P1) from independent donors samples (#17.2, #18.1, #23.1, #18.2, #19.2, #26.1). Blots were probed for pSTAT5, PRLR, β-casein, FOXA1 and KRT8; GAPDH served as a loading control. Molecular mass markers (kDa) are indicated at right. **(B)** Quantification of the indicated proteins in 2D and 3D cultures at P0 and P1, normalized to GAPDH. Dots represent individual donors; bars indicate mean values.

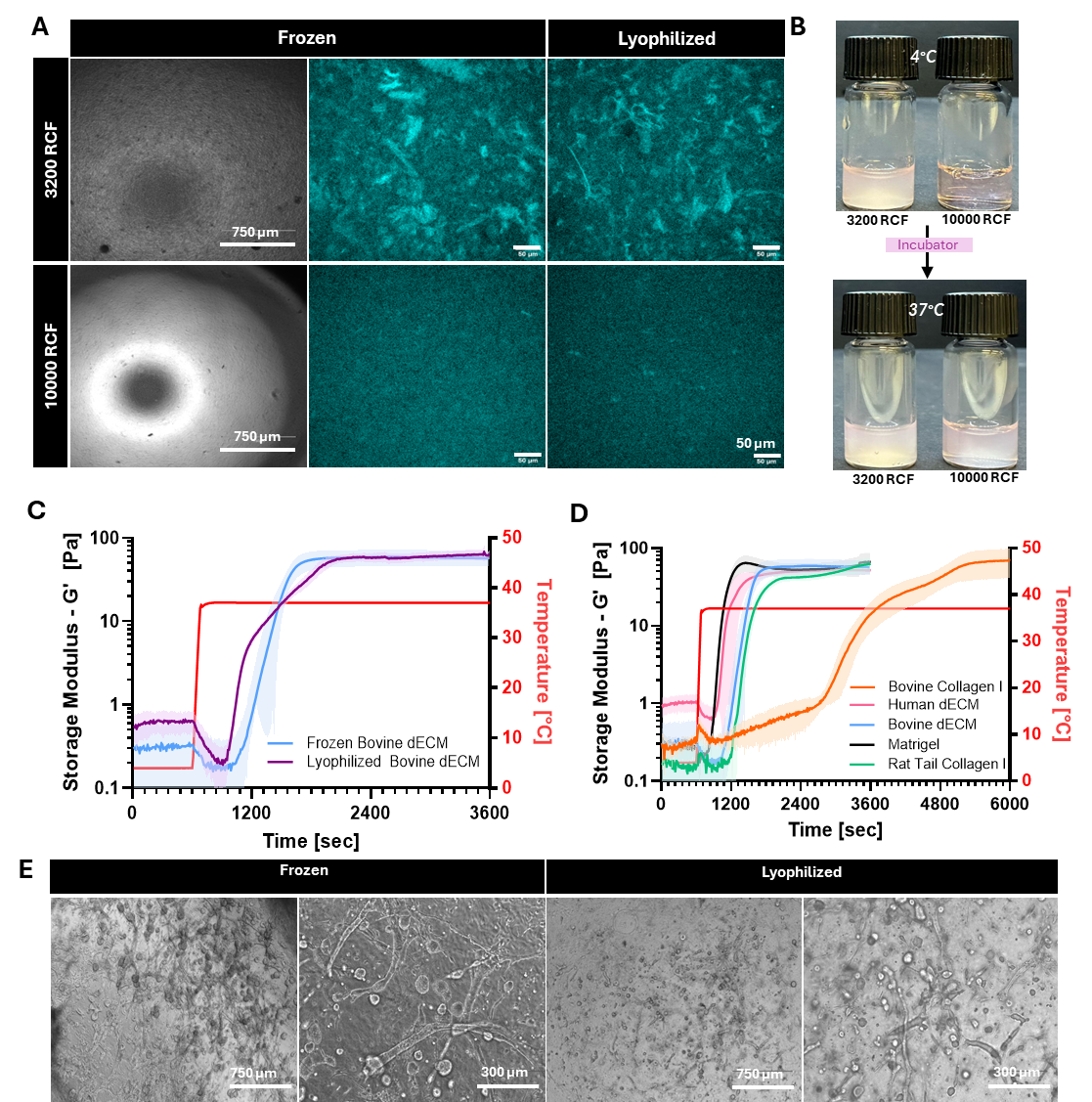

**Figure S6: Optimization of dECM hydrogels. A)** Brightfield and SHG images of bovine dECM hydrogels demonstrate that higher centrifugation speed efficiently removed large particles. **B)** Images showing increased bovine dECM gel clarity at higher centrifugation speed both before and after thermal gelation (4 °C and 37 °C). **C)** Rheological analysis reveals differences in gelation kinetics between lyophilized and frozen bovine dECM. **D)** Rheological analysis of bovine collagen I. Rheology of bovine collagen I hydrogels (2.6 mg/mL) reveals delayed gelation compared to other hydrogel materials, and therefore rat tail collagen was used. **E)** Comparison of MCF10A organoids cultured for five days in frozen versus lyophilized bovine dECM shows increased growth and branching in the frozen gels. Scale bars as indicated.

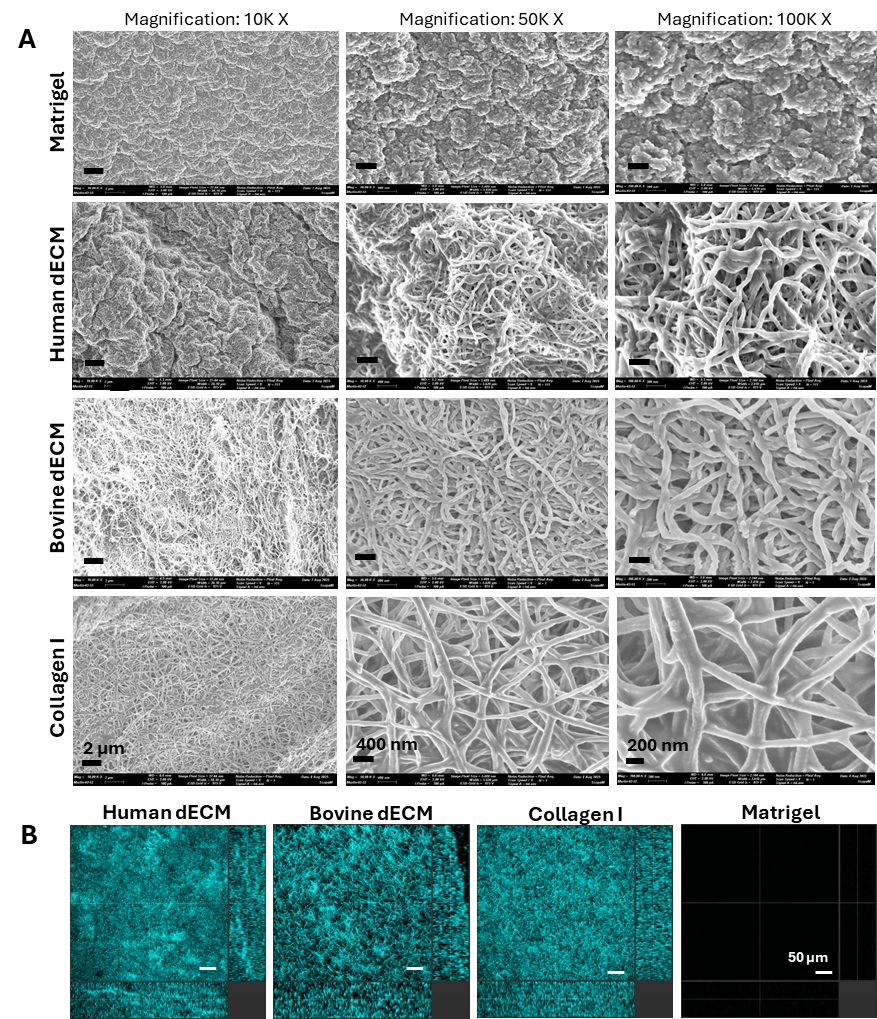

**Figure S7. Ultrastructural and fibrillar organization of thermally gelled matrices.** **A)** Scanning electron microscopy (SEM) and second harmonic generation (SHG) imaging of decellularized extracellular matrix (dECM), collagen, and Matrigel gels following thermal gelation. SEM micrographs (top panels) show representative surface morphology across increasing magnification (rows), highlighting differences in pore/lamellar architecture and fibrillar network organization between matrices (top to bottom: Matrigel, human dECM, bovine dECM, Collagen I). **B)** SHG images (bottom panels; cyan) visualize SHG-active fibrillar collagen content/organization within each gel; the rightmost image is a no-signal/blank reference. Scale bars are indicated in each panel (including 2 µm, 400 nm, 200 nm for SEM examples, and 50 µm for SHG).

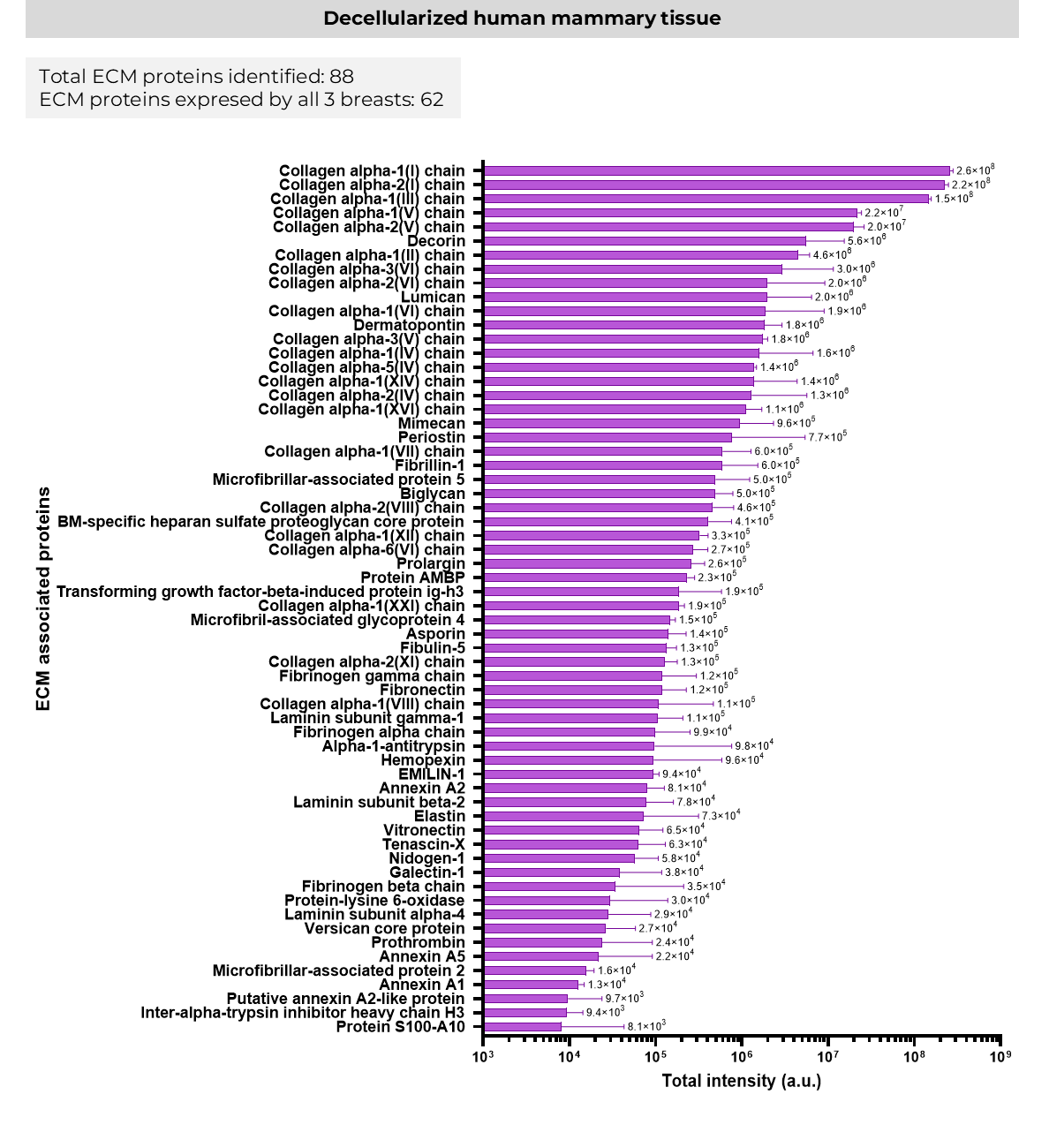

**Figure S8: Relative abundance of the human breast ECM proteins identified by mass spectrometry**

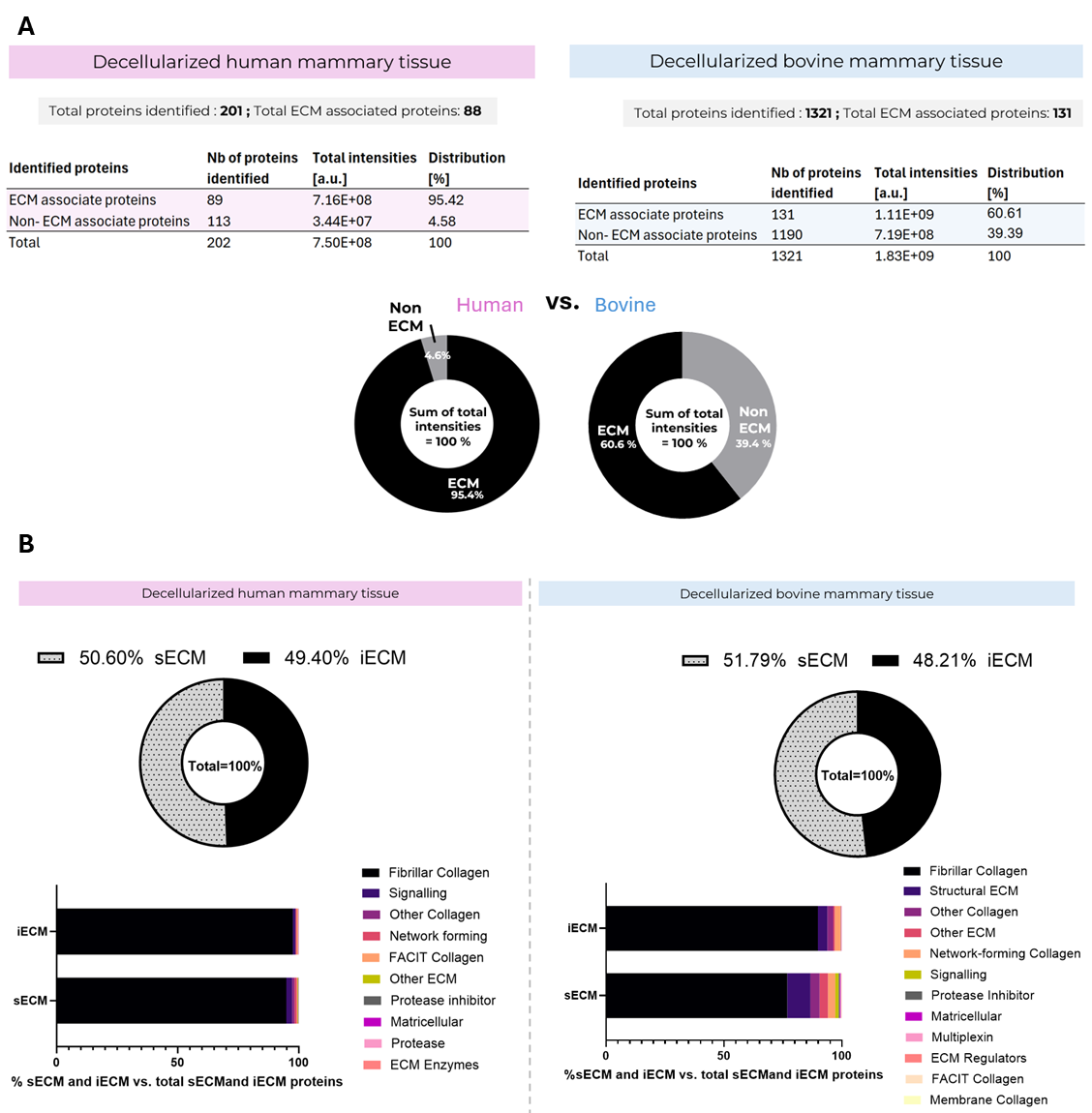

**Figure S9: Proteomic overview of decellularized mammary matrices. A)** Proteomic partitioning of ECM versus non-ECM: Summary of proteins identified by LC–MS/MS in decellularized human mammary tissue (left) and decellularized bovine mammary tissue (right), reporting the number of proteins and summed signal intensities (a.u.) assigned to ECM-associated versus non-ECM proteins. Donut plots depict the corresponding fraction of total protein intensity attributable to ECM or non-ECM components for each source. **B)** ECM proteins were further partitioned into insoluble ECM (iECM) and soluble ECM (sECM) fractions; donut plots show the relative contribution of sECM and iECM to total ECM signal. Horizontal stacked bars summarize the distribution of ECM functional groups (DAVID GO categories; e.g., fibrillar collagen, structural ECM, network-forming collagens, signaling and regulatory components) within iECM and sECM for each tissue source. Overall, human dECM is highly enriched for ECM-associated signal, whereas bovine dECM contains a larger non-ECM fraction; both sources exhibit broadly similar sECM/iECM partitioning with source-dependent differences in ECM sub-class composition.

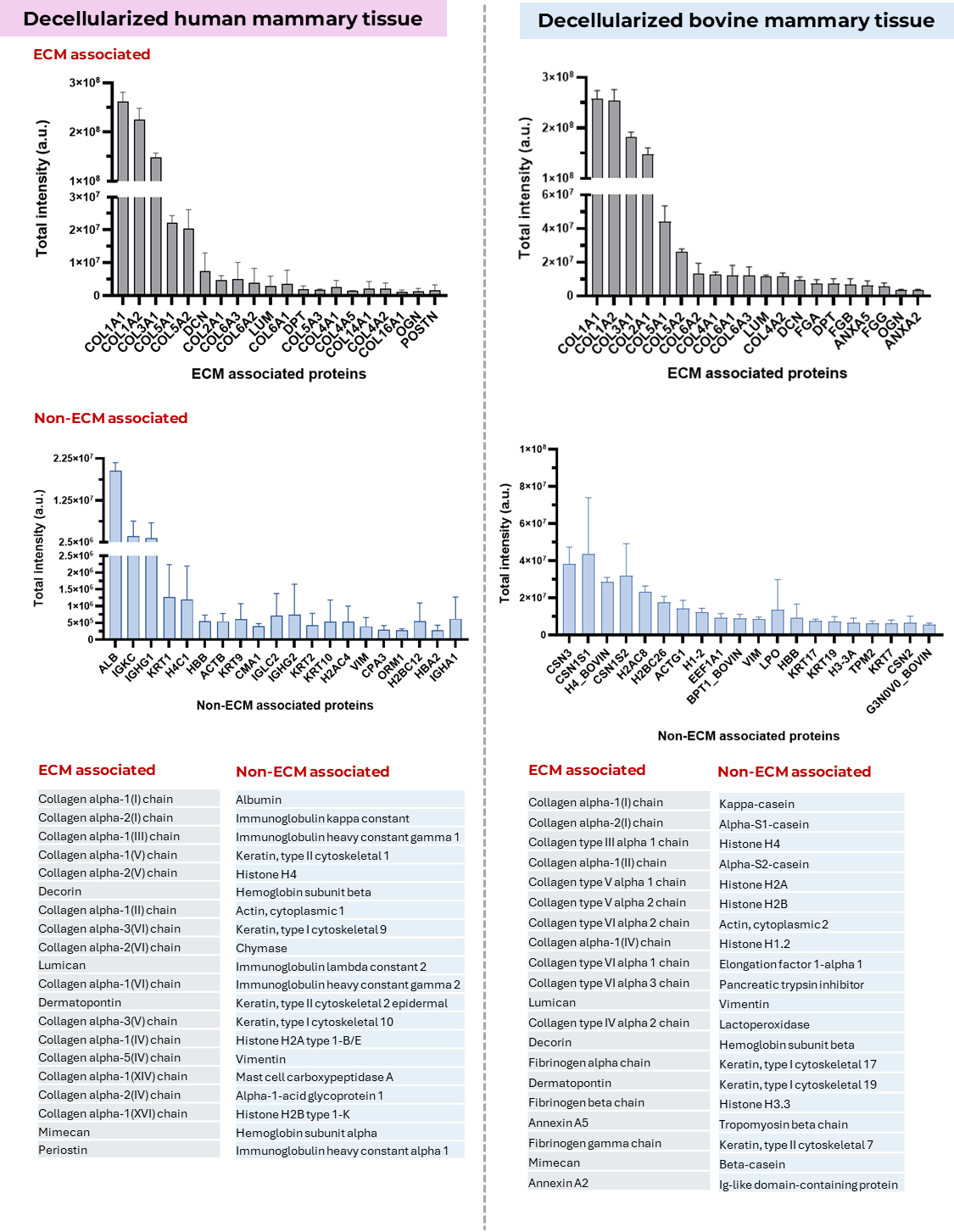

**Figure S10:** **Most abundant ECM- and non-ECM–associated proteins in human and bovine mammary dECM.** Protein abundance is plotted as total intensity (a.u.); bars represent mean +/- SD with n = 3. Tables list the corresponding top-ranked proteins for each category.

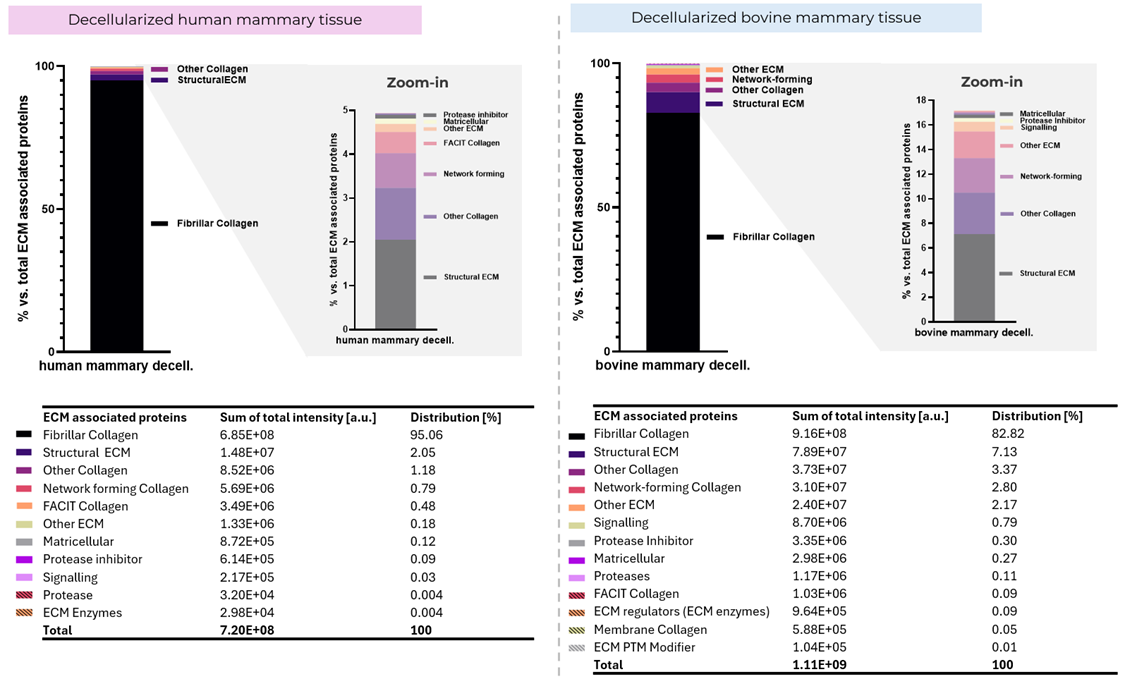

**Figure S11:** **Composition of decellularized human and bovine mammary matrices.** Relative abundance of ECM-associated proteins identified by proteomics in decellularized human mammary tissue (left) and decellularized bovine mammary tissue (right), grouped using DAVID Gene Ontology functional categories. Stacked bars show the percent contribution of each ECM group to the total ECM-associated protein signal; zoom-in panels expand low-abundance categories to facilitate comparison. Tables below report the corresponding summed intensities (a.u.) and distributions (%) for each category. In both matrices, fibrillar collagens comprise the dominant fraction, with bovine dECM showing a comparatively larger contribution from additional ECM classes (e.g., structural ECM, other collagen, and network-forming components) relative to human dECM.

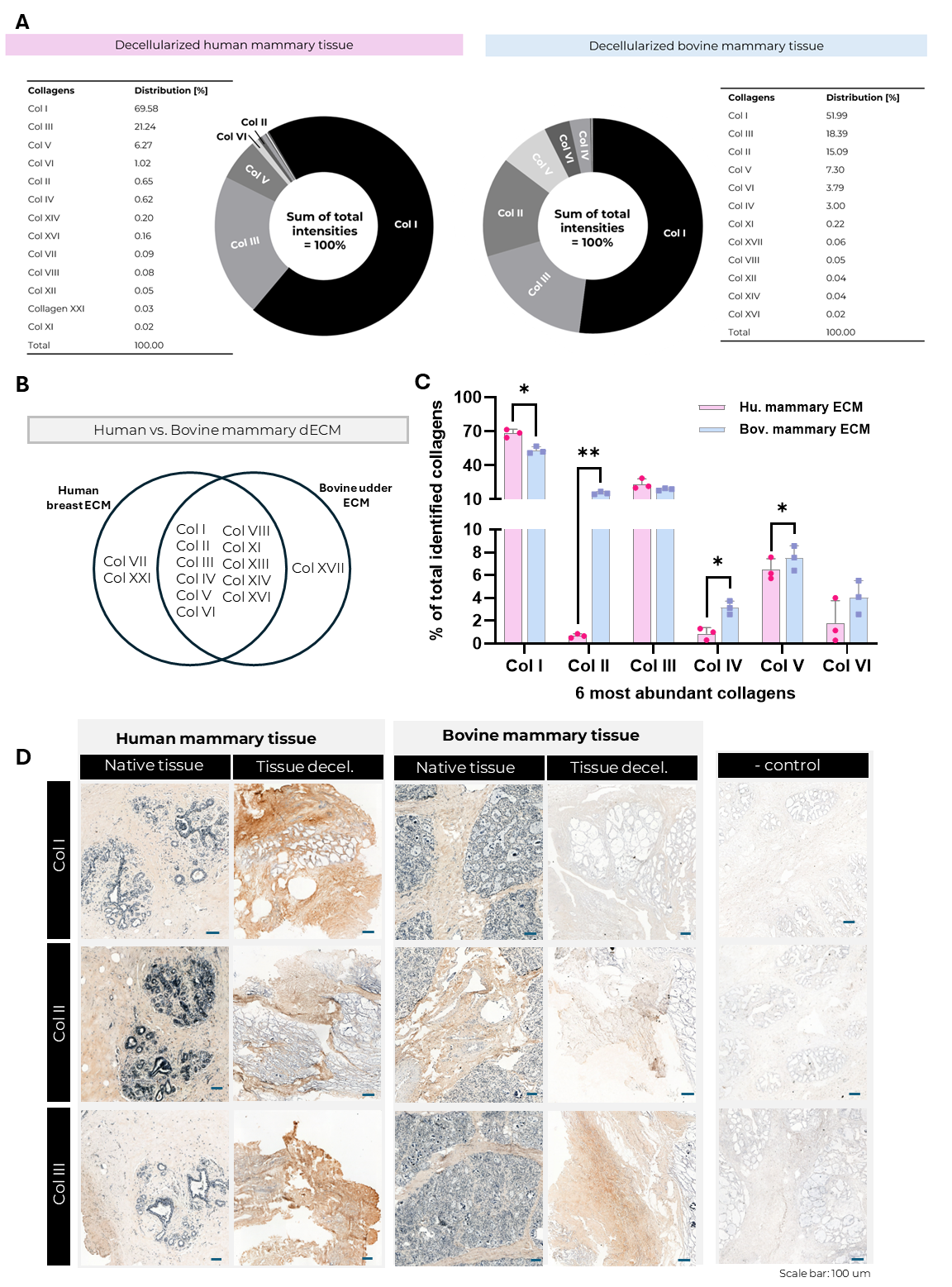

**Figure S12. Collagen composition in human and bovine mammary dECM.** **A)** Proteomics-based distribution of collagen subtypes in decellularized human mammary tissue (left) and decellularized bovine mammary tissue (right). Donut charts show the relative contribution of each collagen type to the summed collagen intensity (total = 100%), with accompanying tables listing percent distributions. **B)** Comparison of collagen species detected in human versus bovine mammary dECM (Venn diagram) and **C)** relative abundance of selected collagen classes/types. **D)** Representative immunohistochemistry for collagen I (Col I), collagen II (Col II), and collagen III (Col III) in native tissue and after decellularization for human and bovine mammary samples, with negative control shown at right.

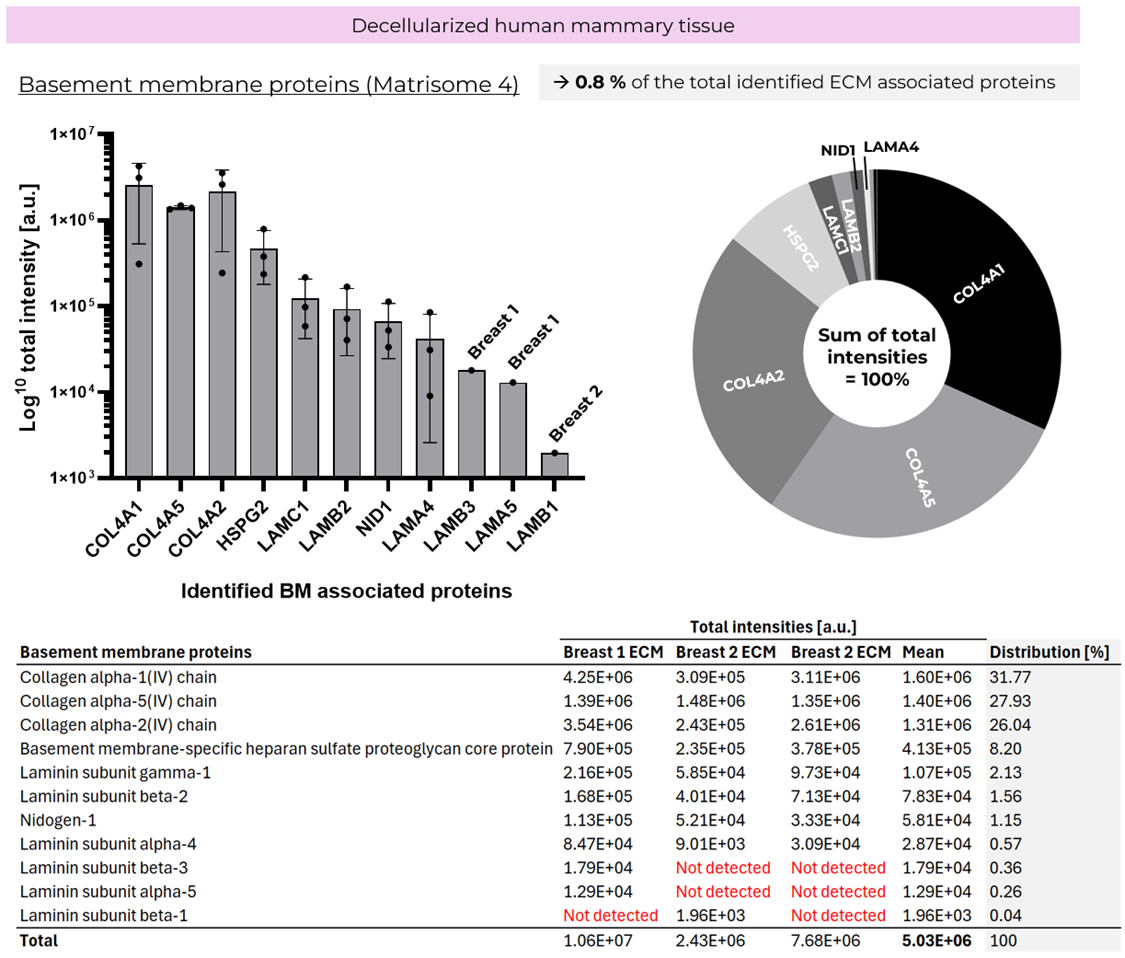

**Figure S13. Basement membrane proteins in human mammary dECM.** LC–MS/MS quantification of basement membrane (BM) proteins in decellularized human mammary tissue (~0.8% of total ECM-associated signal). Left, log10(total intensity, a.u.) for identified BM proteins (points = individual samples; bars/error bars = summary across samples). Right, relative contribution of each BM protein to total BM intensity. Bottom, per-sample intensities, mean, and distribution; “Not detected” indicates below detection.

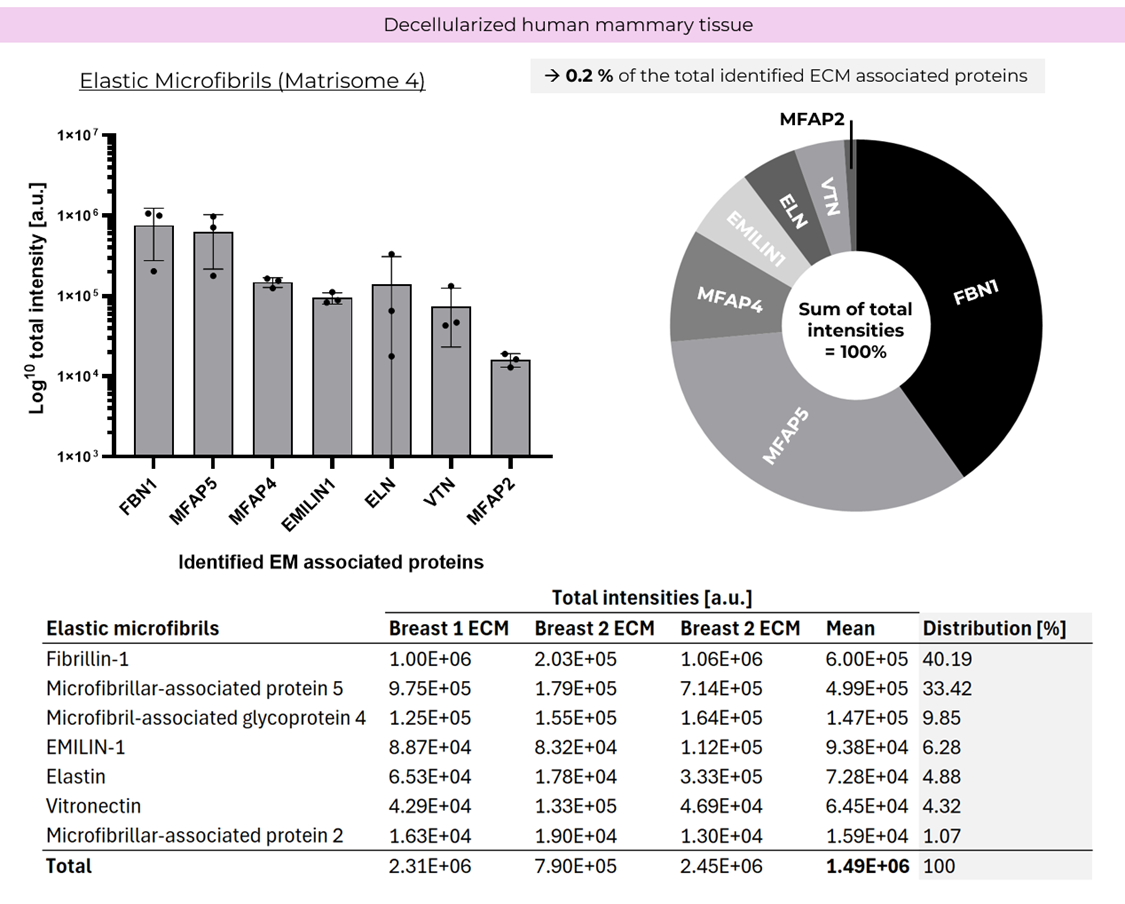

**Figure S14. Elastic microfibril proteins in human mammary dECM.** LC–MS/MS quantification of elastic microfibril–associated proteins in decellularized human mammary tissue (~0.2% of total ECM-associated signal). Left, log10(total intensity, a.u.) for identified elastic microfibril components (points = individual samples; bars/error bars = summary across samples). Right, relative contribution of each protein to total elastic microfibril intensity. Bottom, per-sample intensities, mean, and distribution.

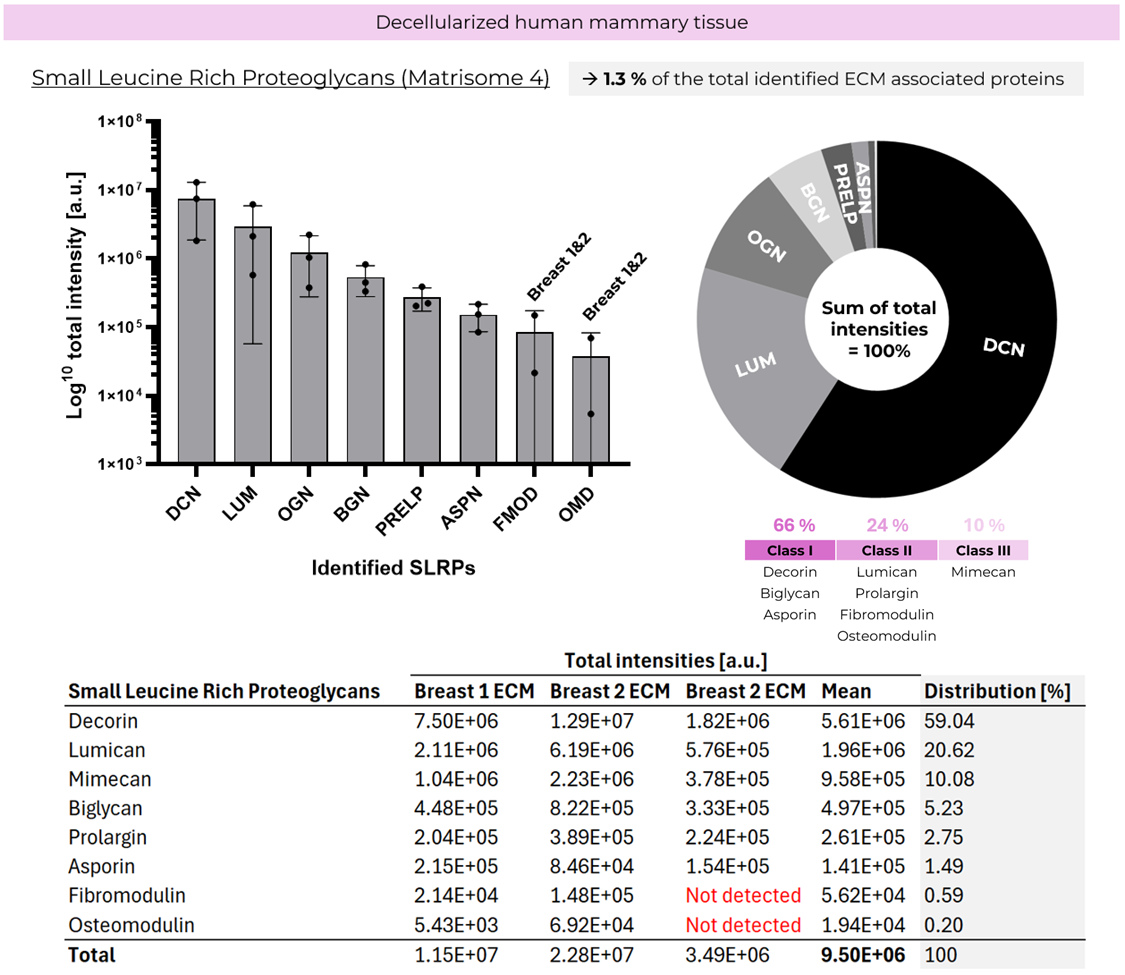

**Figure 15. Small leucine-rich proteoglycans (SLRPs) in decellularized human mammary dECM.** Bar plot of SLRP abundances shown as log10(total intensity, a.u.) and points indicate individual samples/replicates. Right: Donut chart showing the relative contribution of each SLRP to the summed SLRP signal (total = 100%). SLRP class assignment. Identified SLRPs are grouped into Class I, Class II, and Class III based on established SLRP subfamilies defined by protein core features (including leucine-rich repeat organization), N-terminal cysteine cluster pattern, and typical glycosylation type: Class I (canonical dermatan/chondroitin sulfate SLRPs): Decorin (DCN), Biglycan (BGN), Asporin (ASPN); Class II (keratan sulfate–associated / fibromodulin subfamily): Lumican (LUM), Prolargin (PRELP), Fibromodulin (FMOD), Osteomodulin (OMD); Class III (atypical SLRPs): Mimecan/Osteoglycin (OGN). Bottom: Table reporting per-sample intensities, mean intensities, and percentage distribution; “Not detected” indicates proteins below detection in the indicated sample.

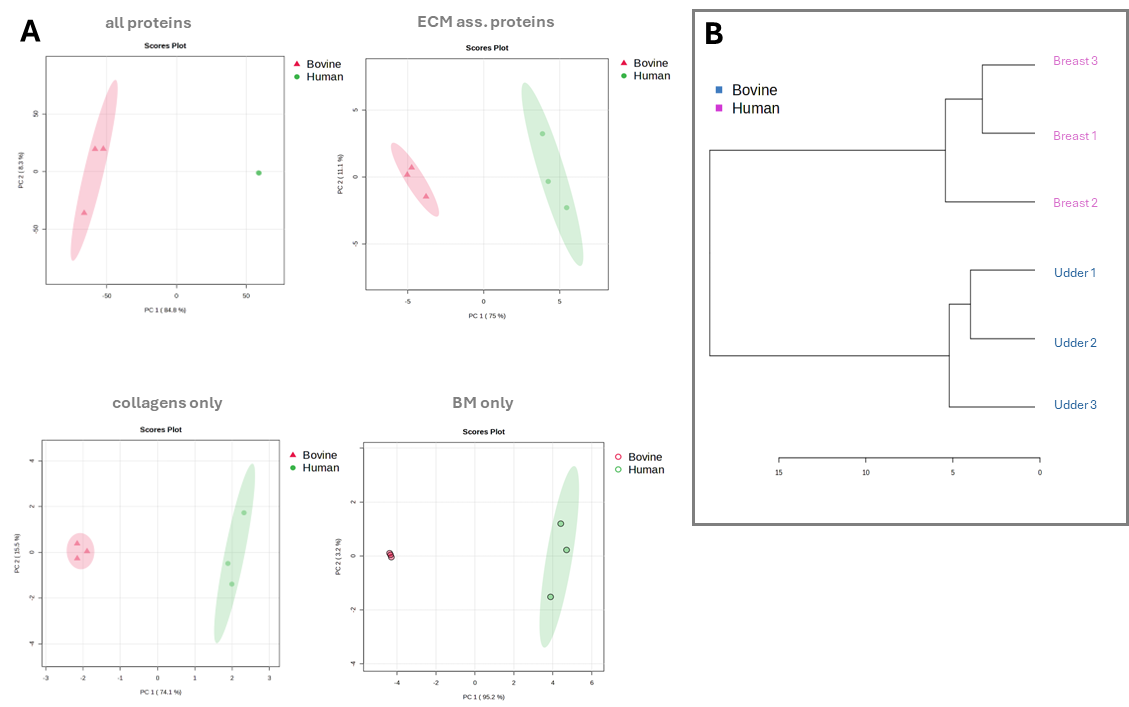

**Figure S16. Proteomic separation of human breast and bovine udder dECM. A)** Principal component analysis (PCA) score plots comparing decellularized human mammary tissue (green) and decellularized bovine mammary tissue (pink) of all proteins, ECM-associated proteins, collagens only, and basement-membrane (BM) proteins only. **B)** Unsupervised hierarchical clustering of samples based on ECM-associated proteomic profiles shows robust grouping by tissue source, separating human breast from bovine udder decellularized matrices.

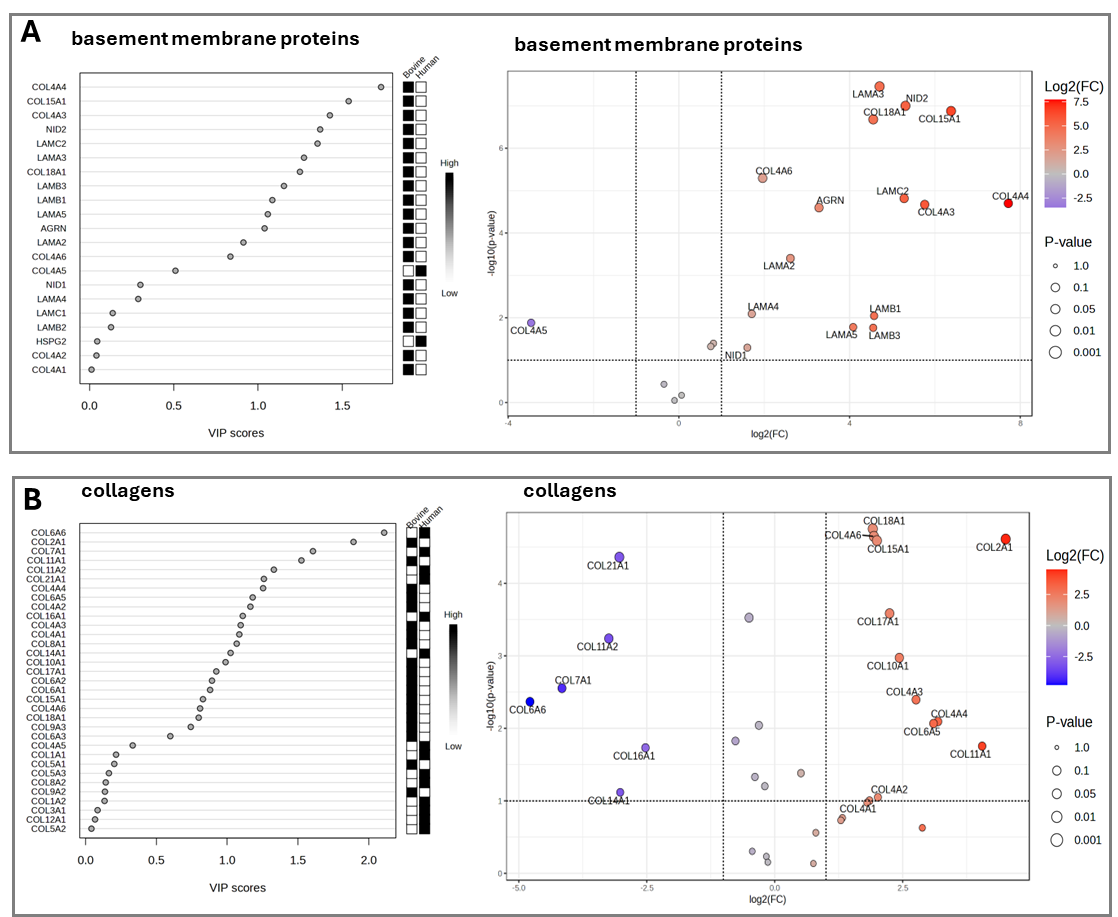

**Figure S17. Differential basement-membrane and collagen signatures distinguish human and bovine mammary dECM**. Proteomics-based multivariate and univariate analyses comparing decellularized human mammary tissue and decellularized bovine mammary tissue.
**A)** Basement membrane proteins. Left, variable-importance-in-projection (VIP) scores from PLS-DA identify BM proteins that most strongly contribute to separation between tissue sources; the adjacent heat strip indicates relative abundance per source (low–high). Right, volcano plot showing log2(fold change) (human vs bovine, as defined in the analysis) versus –log10(p value) for BM proteins; point color encodes log2(FC) and point size reflects significance. **B)** Collagens. Left, VIP ranking for collagen proteins contributing to group separation with corresponding relative-abundance heat strip. Right, volcano plot of collagen proteins highlighting differentially abundant collagen chains between human and bovine dECM. Vertical and horizontal dashed lines denote fold-change and p-value thresholds used for significance.

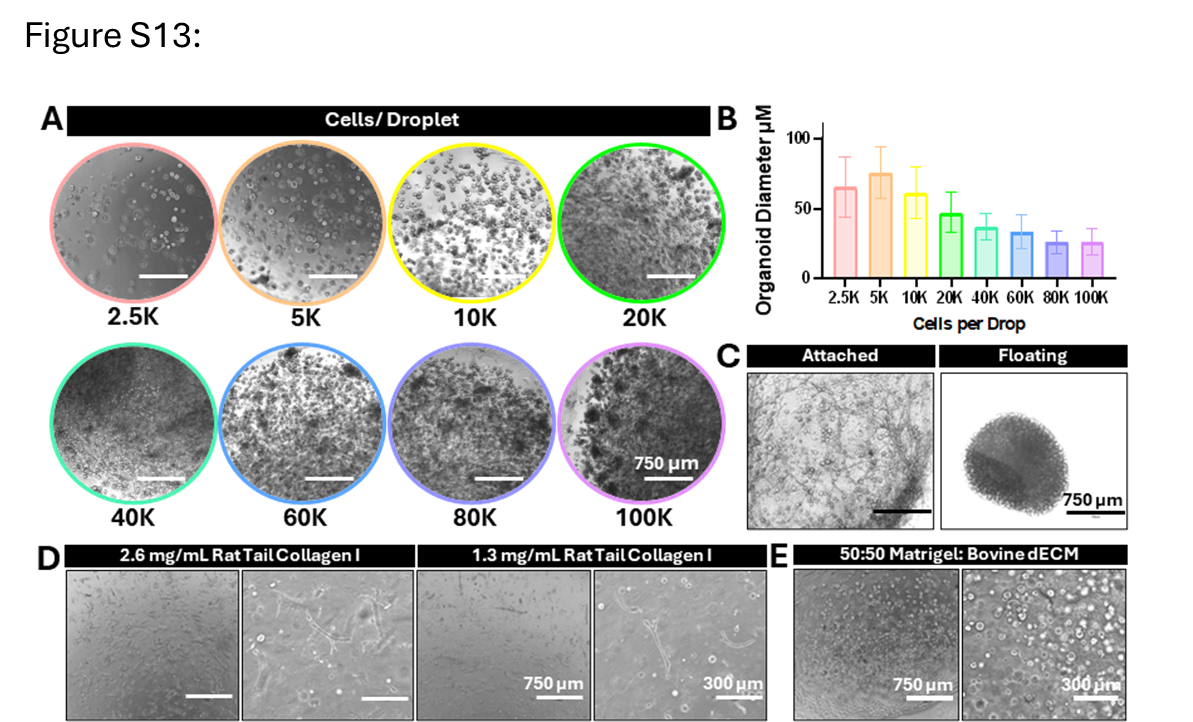

**Figure S18. Optimization of MCF10A organoid culture. A)** Representative images of organoids formed at seeding densities of 2.5 K–100 K cells per 30 μL Matrigel droplet. **B)** Quantification of average organoid diameter across seeding densities (mean ± SD, n = 50 organoids per condition). **C)** Comparison of organoids grown in attached versus floating bovine dECM droplets showing significant shrinkage in the floating droplet. **D)** Representative images of organoids cultured in stiffness-matched collagen I (2.6 mg/mL) and lower-concentration collagen I (1.3 mg/mL) where no appreciable difference in organoid size or branching morphology was observed. **E)** Representative images of MCF10A cultured in 50:50 Matrigel:bovine dECM gels showing organoids with predominantly spherical morphology. Scale bars as indicated.

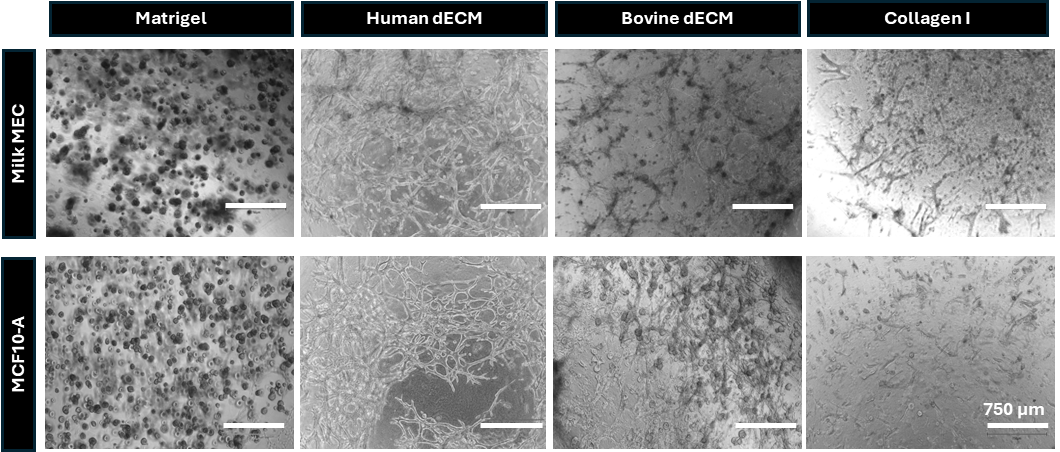

**Figure S19. Brightfield overview of organoid morphology.** Representative 4× brightfield images of milk MEC and MCF10A organoids cultured in Matrigel, human dECM, bovine dECM, or collagen I. Morphological differences are apparent even at low magnification: organoids in Matrigel formed exclusively spherical structures, human dECM supported the most extensive cell growth and thickest branches, bovine dECM produced intermediate branching, and collagen I generated sparse networks with the smallest branches. Scale bars 750 μm.

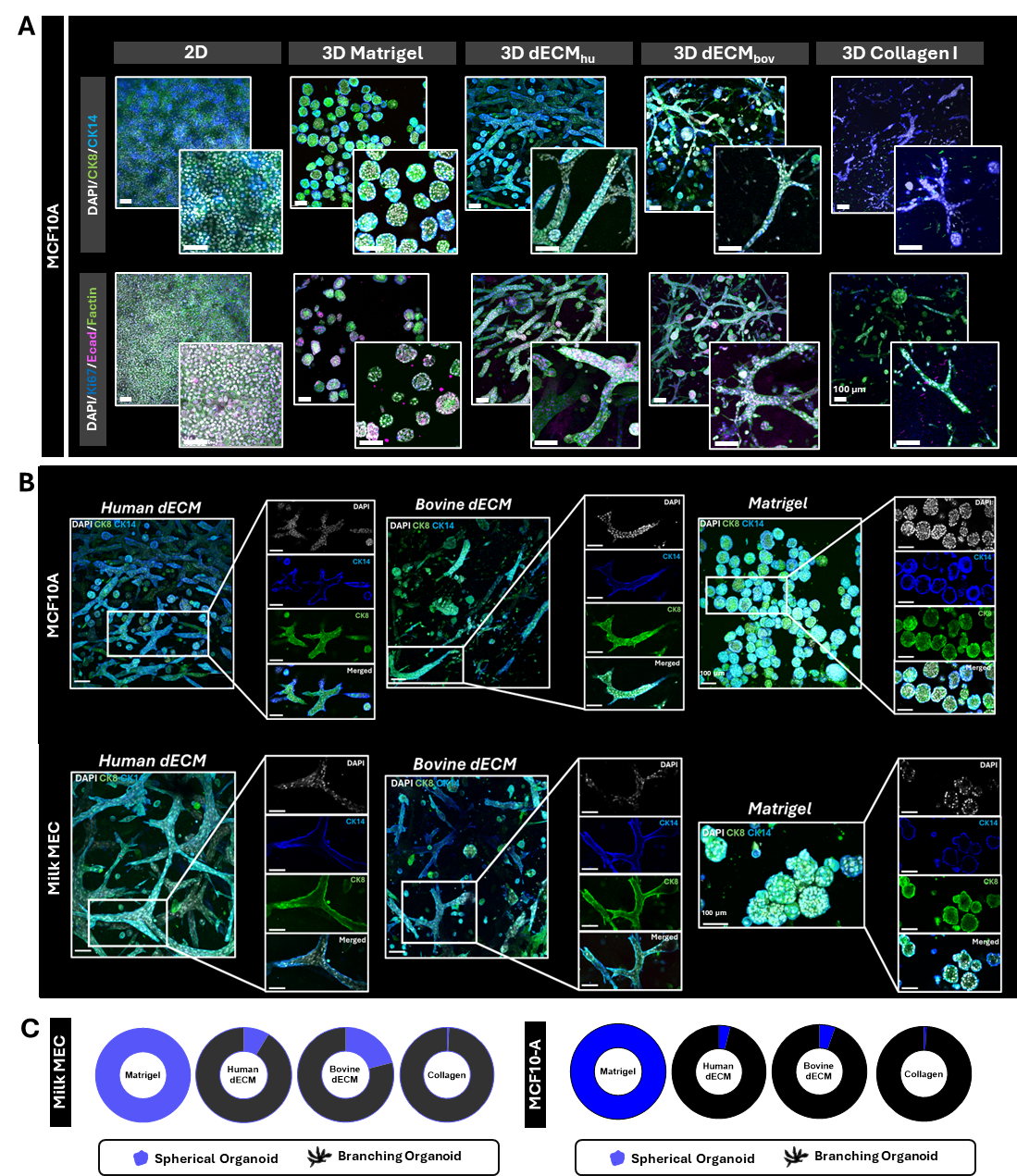

**Figure S20. Matrix-dependent architecture of MCF10A and Milk MEC organoids.** **A-B)** Representative confocal immunofluorescence images of MCF10A and human milk–derived mammary epithelial cells (Milk MECs) cultured in 2D, 3D Matrigel, 3D human mammary dECM (dECM_hu_), 3D bovine mammary dECM (dECM_bov_), or 3D collagen I. Cells are stained for DAPI (nuclei) and epithelial/basal markers CK8 and CK14, as well as F-actin and Ki67. Matrigel predominantly yields rounded spheroids, whereas stromal/fibrillar matrices (dECM and collagen I) support elongated/branched structures with CK14-enriched basal-like cells preferentially localized to the periphery. **C)** Schematics summarize branched and spherical morphologies conditions. Scale bars, 100 µm (unless noted).

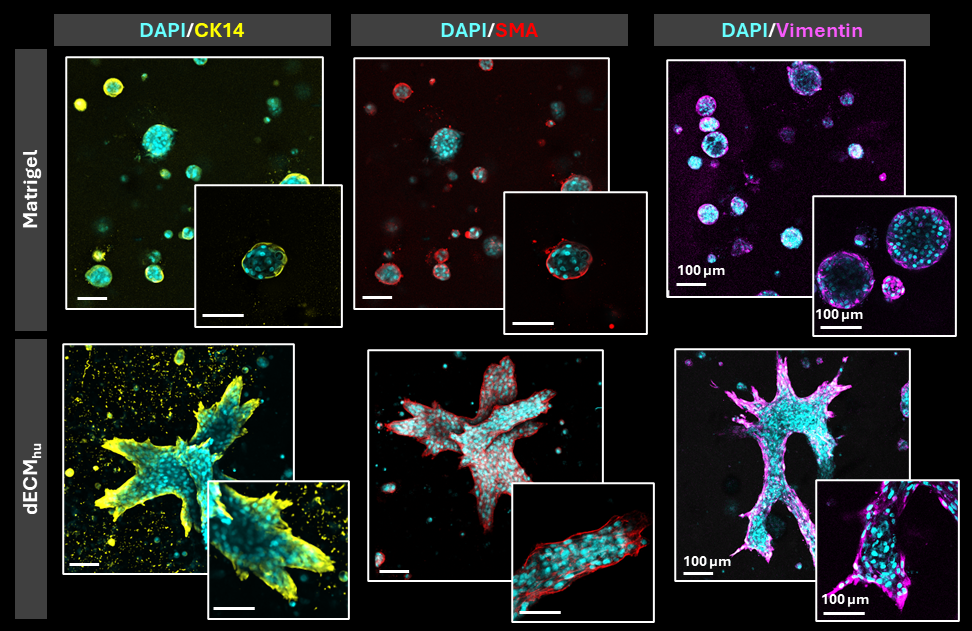

**Figure S21. Matrix-dependent basal/myoepithelial marker localization in Milk MEC organoids.** Representative immunofluorescence images of human milk–derived mammary epithelial cells (Milk MECs) cultured in Matrigel (top row) or human mammary dECM (dECM_hu_) (bottom row). Nuclei are labeled with DAPI (cyan). Columns show staining for KRT14/CK14 (yellow; basal-lineage marker), αSMA (red; myoepithelial/contractile marker), and Vimentin (magenta; mesenchymal-associated intermediate filament that can be enriched in basal-like/myoepithelial or partial EMT/stromal-like states). Matrigel cultures form predominantly rounded spheroids, whereas dECM_hu_ supports elongated/branched multicellular structures. In both matrices, CK14 and αSMA are preferentially localized to the outer cell layer, consistent with basal/myoepithelial-like positioning at the organoid periphery. Insets show higher-magnification views. Scale bars, 100 µm.

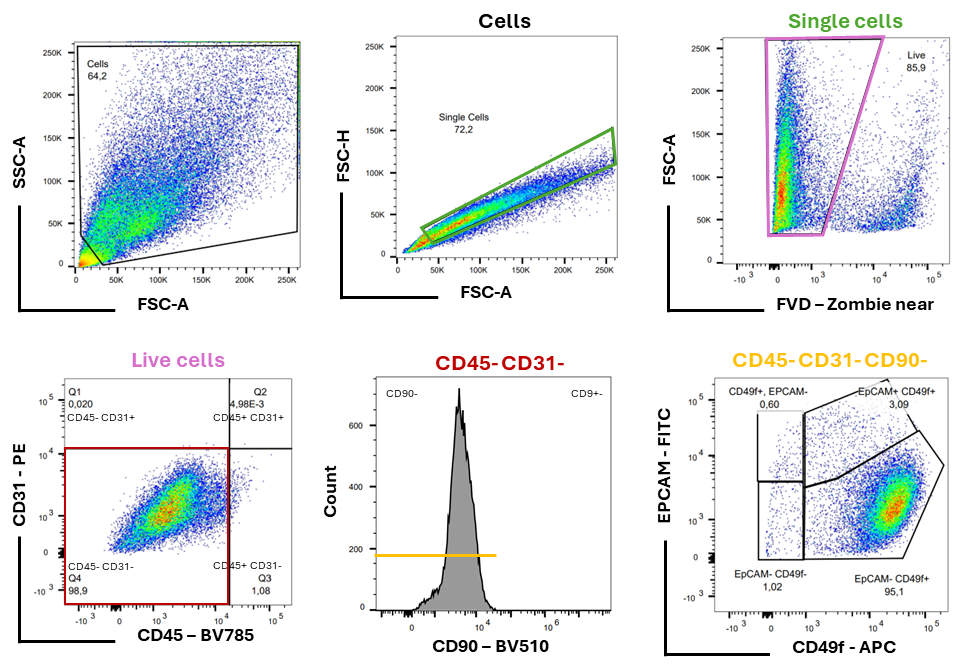

**Figure 22. Flow Cytometry gating strategy for milk MEC grown in 3D and 2D conditions.** Flow cytometric panel was designed using previously established protocols to identify basal and luminal mammary epithelial cells. (The example shown corresponds to milk MEC organoids cultured in Matrigel).

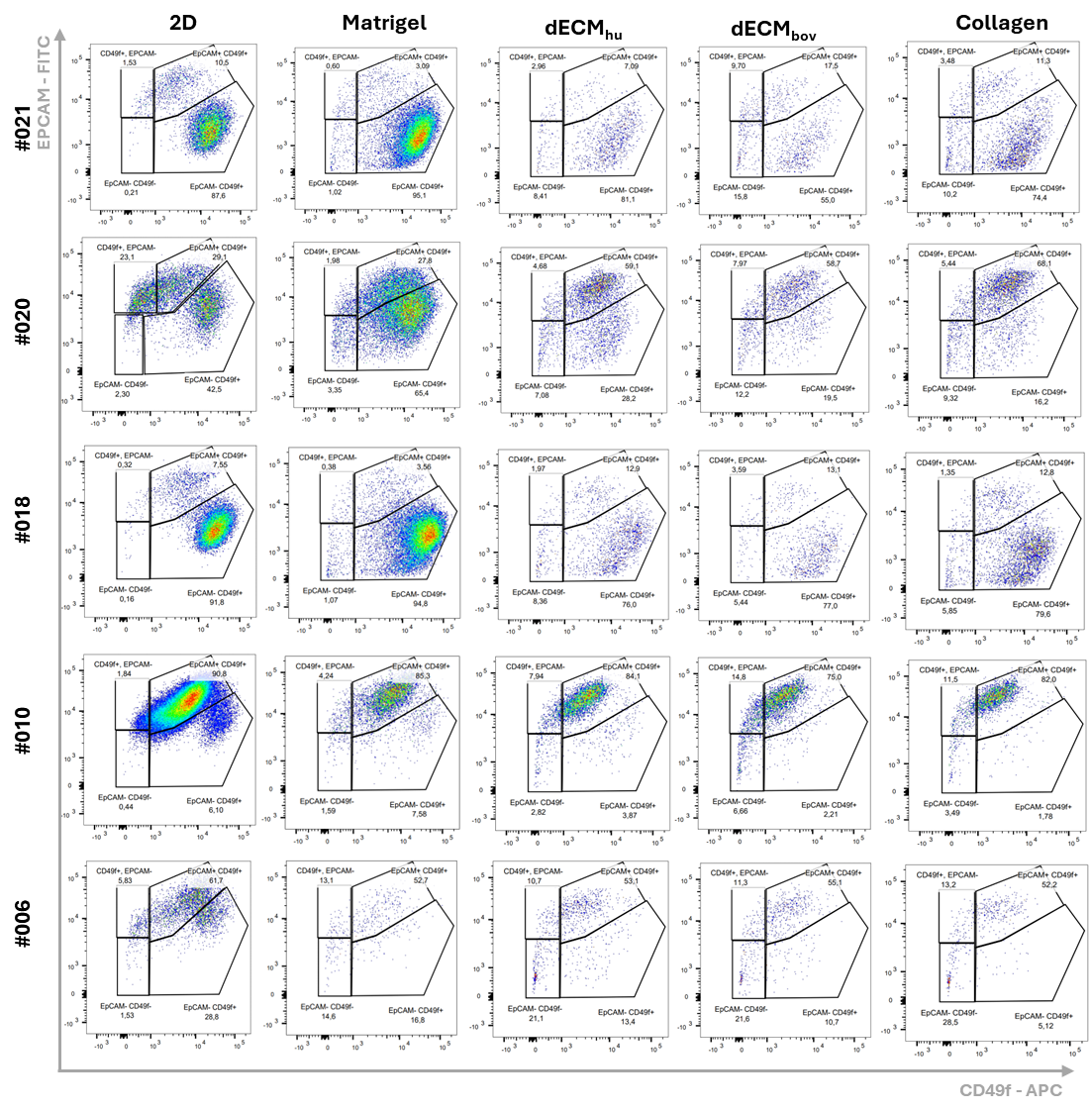

**Figure S23. Mammary epithelial cell populations in different culture conditions analyzed by Flow cytometry.** Milk derived cells were cultured in vitro for 8 days and split at 85-90 % confluency. The cells display mature luminal MEC (EPCAM+ CD49f-), luminal progenitor MEC (EPCAM+ CD49f+) and basal-like MEC phenotypes (EPCAM- CD49f+) (n= 5 donors).

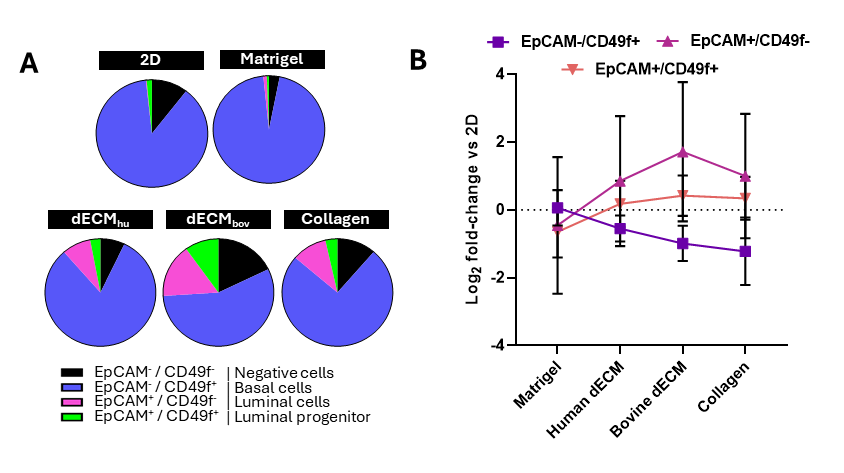

**Figure S24. Flow cytometry summary of human milk–derived mammary epithelial cells across culture conditions.** Human milk–derived mammary epithelial cells from 5 independent donors were cultured under the indicated conditions in different materials :2D, Matrigel, human dECM (dECM_hu_), bovine dECM (dECM_bov_), or collagen; and analyzed by flow cytometry after dissociation to single cells. **A)** Pie charts show the mean fraction of cells in each EpCAM/CD49f gate across donors: EpCAM⁻/CD49f⁻ (double negative, black), EpCAM⁻/CD49f⁺ (CD49f⁺, blue), EpCAM⁺/CD49f⁻ (EpCAM⁺, pink), and EpCAM⁺/CD49f⁺ (double positive, green). **B)** Line plot shows log₂ fold-change in gate frequency relative to 2D across ECM/3D conditions (Mean +/- SD, n = 5).

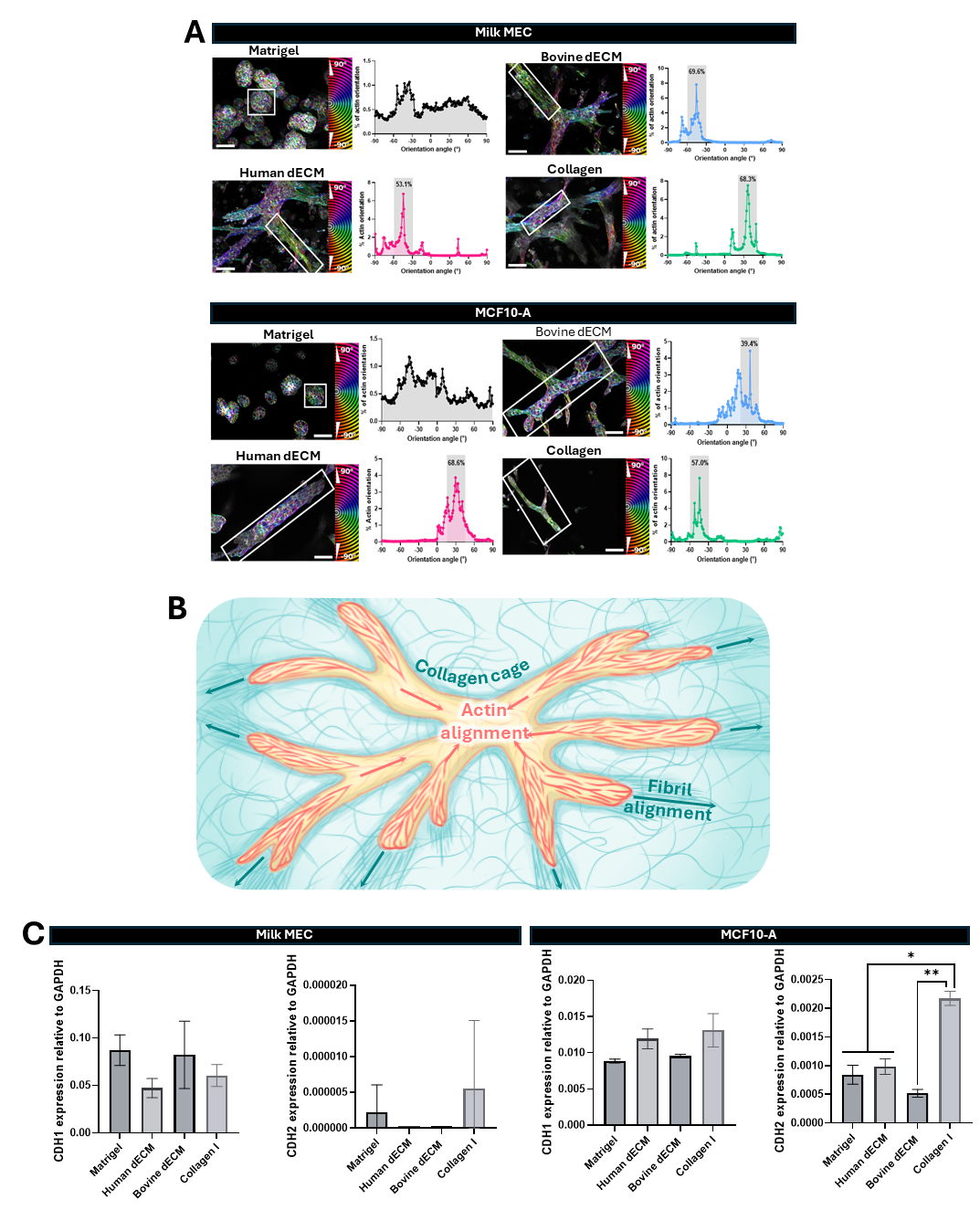

**Figure S25. Collagen-rich matrices promote cytoskeletal alignment and matrix-dependent cadherin expression in milk MEC.** **A)** Representative images of F-actin organization in human Milk MEC and MCF10A cultured in different matrix conditions. Outlined regions were used for orientation analysis. Plots show the distribution of actin orientations for each corresponding region. **B)** Schematic illustrating the proposed response to collagen-rich matrices, in which a surrounding “collagen cage” constrains cell shape and is associated with actin alignment and collagen fibril alignment along the same principal axis. **C)** RT–qPCR quantification of CDH1 and CDH2 in Milk MEC and MCF10A cultured in Matrigel, human dECM, bovine dECM, or collagen I. Expression is shown relative to GAPDH. (Mean +/- SD, *p < 0.05, *p < 0.01)

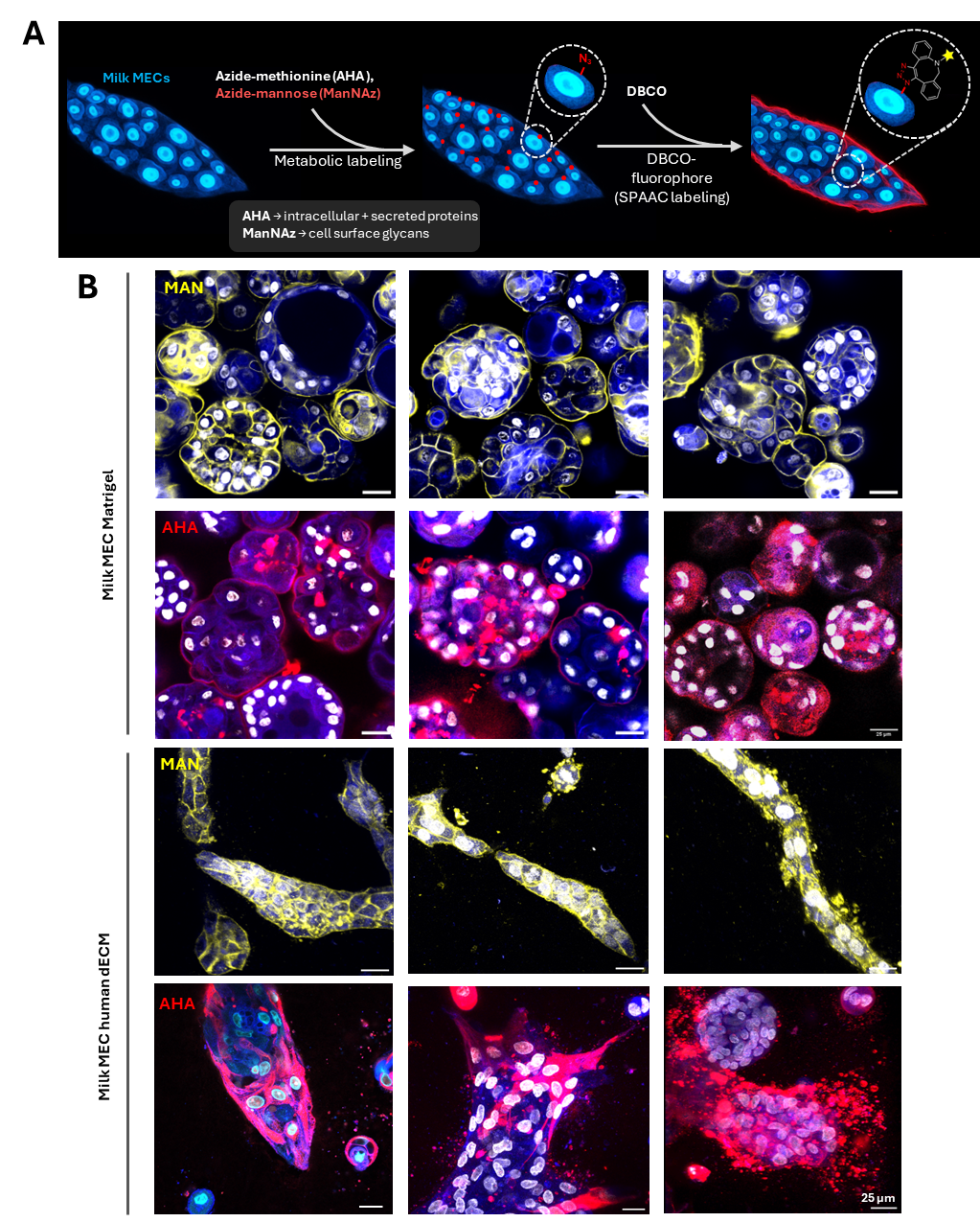

**Figure S26. Metabolic labeling shows nascent ECM deposition by milk MECs in Matrigel and in human dECM.** **A)** Schematic of the dual metabolic labeling strategy to visualize newly synthesized extracellular matrix (ECM). Azide–methionine (AHA) incorporates into nascent proteins, whereas azide–mannose (MAN) labels newly synthesized glycoproteins and proteoglycans (nascent proteoglycans), which are subsequently detected by click-chemistry–based fluorescent tagging to highlight secreted ECM. **B)** Representative confocal images of milk MEC organoids cultured in Matrigel showing MAN signal (yellow; top row) and AHA signal (red; bottom row). Nuclei are counterstained (blue/white). **C)** Representative confocal images of milk MECs cultured on/in human dECM showing MAN signal (yellow; top row) and AHA signal (red; bottom row). Scale bars, 25 µm.

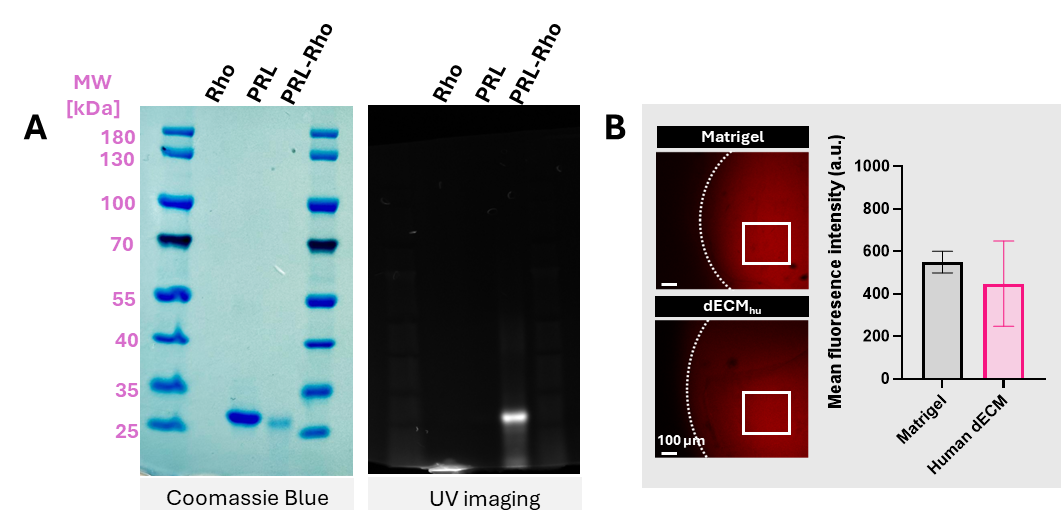

**Figure S27. Fluorescent labeling and hydrogel diffusion of prolactin.** **A)** Recombinant prolactin (PRL) was fluorescently labeled with NHS–rhodamine and labeling was verified by SDS–PAGE. Coomassie Blue staining shows total protein and UV imaging confirms rhodamine fluorescence associated with PRL at the expected molecular weight. **B)** Labeled PRL was then pipetted onto Matrigel or human dECM (dECM_hu_) hydrogels and its diffusion was assessed by fluorescence imaging (representative fields shown; dashed line indicates gel boundary and boxed regions indicate the quantification area). Right, bar graph quantifies mean fluorescence intensity (a.u.) within the indicated regions. Data are shown as mean ± SD, n = 3, and no significant difference was detected between Matrigel and dECM_hu_.

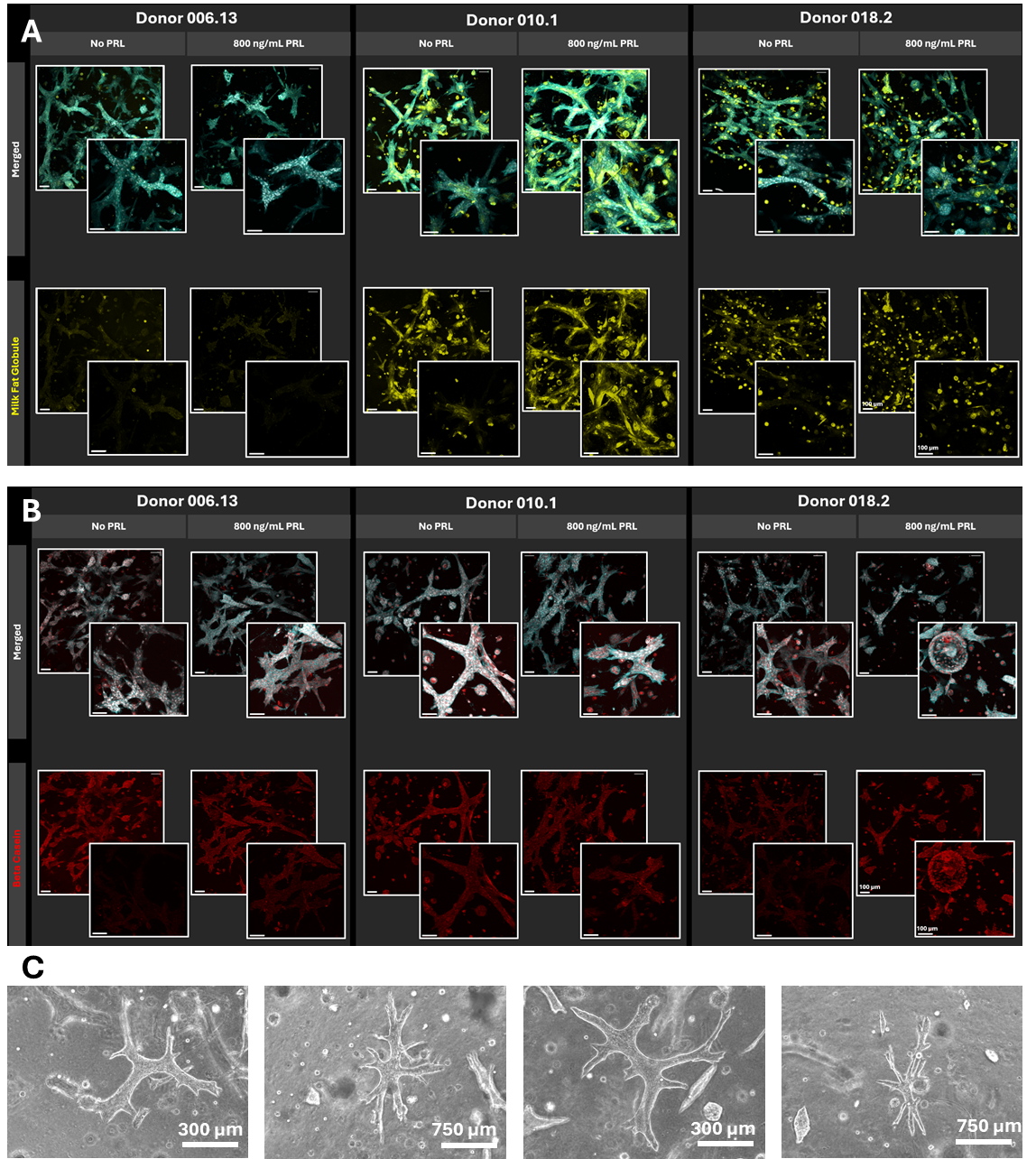

**Figure S28. Milk MEC phenotype and morphology in Matrigel and human dECM.** **A–B)** Representative immunofluorescence images of milk mammary epithelial cells (MECs) cultured in Matrigel or in human decellularized extracellular matrix (dECM) and stained for milk-associated proteins to assess lactation-associated differentiation (markers as indicated in the figure). Nuclei are counterstained with DAPI.

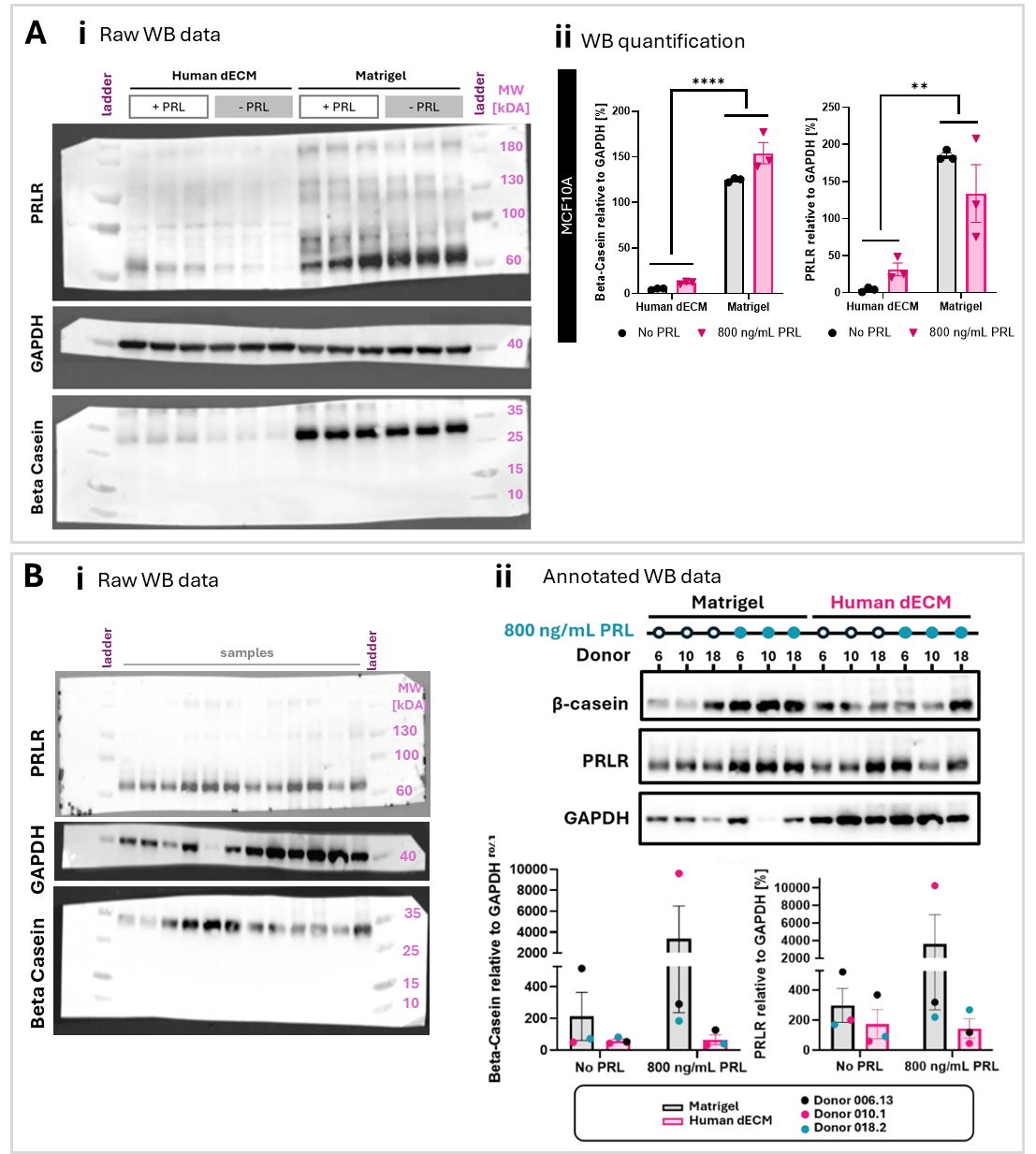

**Figure S29: Western blot (WB) of MCF10A and Milk MECs cultured in Matrigel or human dECM ± prolactin. A) i)** Uncropped, unannotated membranes (raw data) from MCF10A cells cultured in Matrigel or human dECM (dECM_hu_) with or without prolactin (PRL; 800 ng/mL), showing protein bands at the indicated molecular weights. **ii)** Quantification of β-casein (left) and prolactin receptor (PRLR) (right) protein in MCF10A cells cultured in 3D within human decellularized ECM (dECM) or Matrigel, in the absence (No PRL) or presence of 800 ng/mL prolactin (PRL). Signals are normalized to GAPDH and expressed as % relative to GAPDH. (Mean ± SD, ** p < 0.01; **** p < 0.0001, n = 3 biological replicates).**B)** Milk MECs from independent donors were cultured in Matrigel or human dECM (dECM_hu_) with or without prolactin (PRL; 800 ng/mL). **i)** Uncropped, unannotated membranes (raw data) showing total protein loading and protein bands at the indicated molecular weights. **ii)** Cropped and annotated blots from the same membranes for β-casein, PRLR, and GAPDH, with donor IDs indicated. Quantification (β-casein/GAPDH; PRLR/GAPDH, %) is shown with individual donor values color-coded (n=3; mean ± SD. No statistically significant differences were detected between conditions)

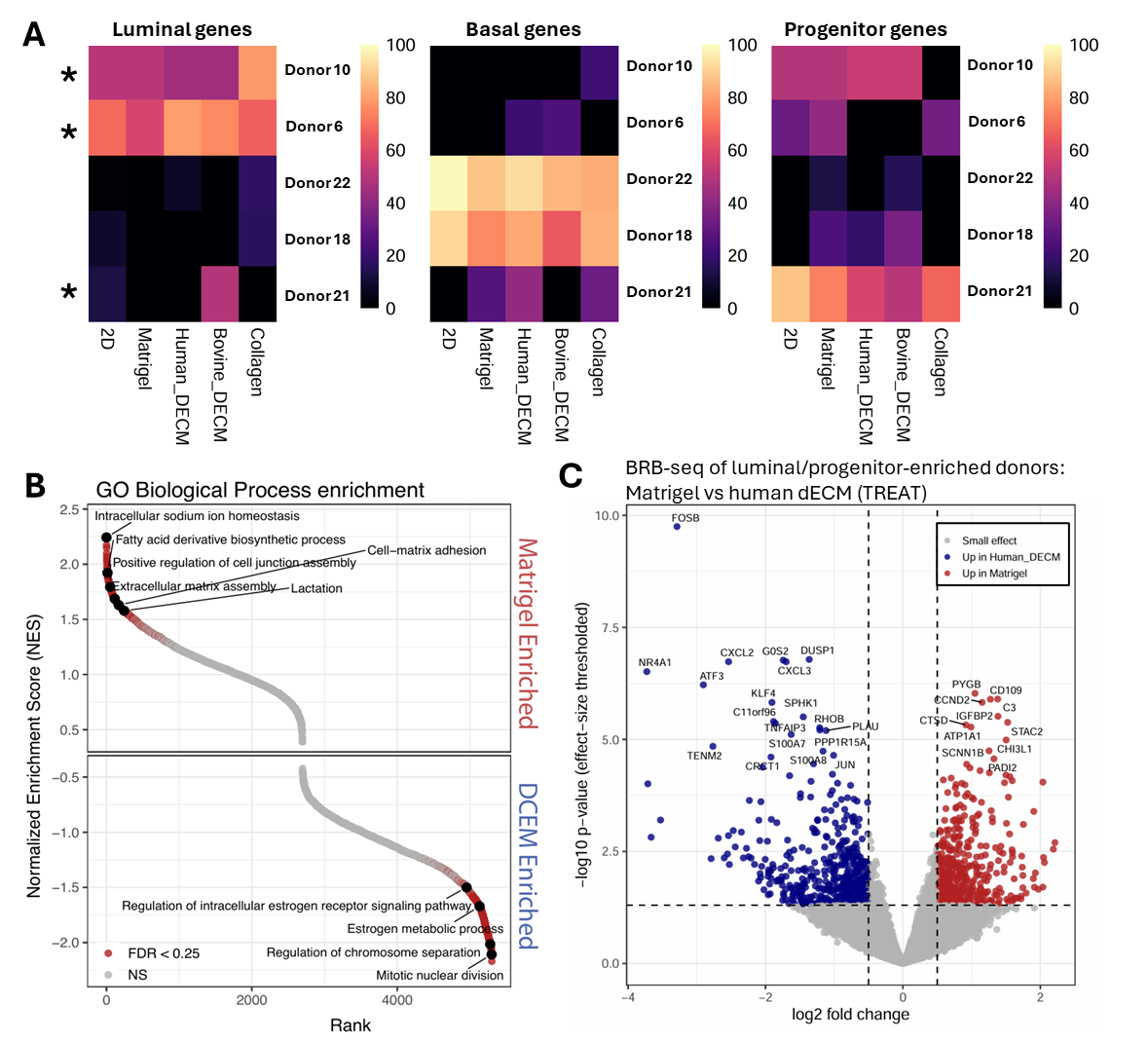

**Figure S30. BRB-seq lineage-signature scoring and differential expression analyses in luminal/progenitor-enriched donors across ECM conditions. A)** Heatmaps showing relative luminal, basal, and progenitor epithelial state scores across ECM conditions (columns: 2D, Matrigel, Human dECM, Bovine dECM, and Collagen) for each donor (row). Scores were derived from curated lineage marker sets using variance-stabilized expression data, gene-wise z-scoring across samples, and per-sample normalization so that luminal, basal, and progenitor program scores summed to 100%. Higher values indicate a stronger relative contribution of the corresponding epithelial program within a given sample. For consequent analyses, the three donors (10, 6 and 21 marked with an asterisk) with the strongest luminal/progenitor-like epithelial compositionwere used (luminal_genes <- c( "ESR1", "PGR", "FOXA1", "GATA3", "GREB1", "TFF1", "TFF3", "XBP1", "AGR2", "MUC1", "CA12","BCL2","SCGB2A2") ; prog_genes <- c("ELF5", "KIT", "ALDH1A3", "SOX9", "NOTCH1", "NOTCH3", "JAG1", "PROM1", "CD24", "BCL11A"); basal_genes <- c("KRT5", "KRT14", "KRT17", "TP63", "ITGA6", "ITGB4", "LAMB3", "LAMC2", "COL17A1", "EGFR","CDH3")) **B)** Gene set enrichment analysis of Gene Ontology (GO) Biological Process terms in luminal/progenitor donors comparing Matrigel with Human dECM, showing Matrigel-enriched and dECM-enriched pathways ranked by normalized enrichment score (NES); highlighted terms passed the displayed FDR threshold of 0.25. **C)** Volcano plot showing limma-TREAT differential expression results for luminal/progenitor donors cultured in Matrigel versus Human dECM. Genes are displayed by log2 fold change and effect-size-thresholded p-value, with positive values indicating Matrigel-enriched expression and negative values indicating Human dECM-enriched expression. Dashed lines mark the TREAT fold-change threshold (|log2FC| = 0.5) and nominal p-value cutoff (p = 0.05); labeled genes represent selected differentially expressed transcripts.

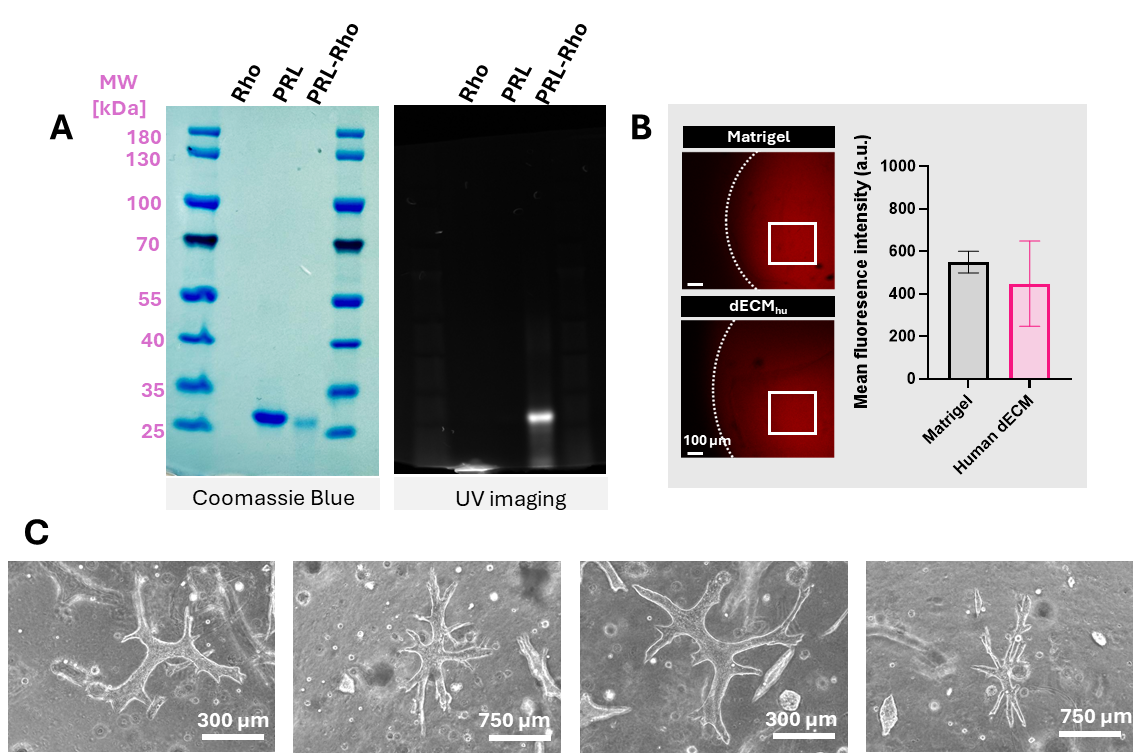

**Figure S31. Milk MEC at lower seeding density with PRL stimulation.** Representative brightfield images of milk MECs cultured in human dECM at lower seeding density and stimulated with prolactin (PRL) to evaluate emergence of terminal duct lobular unit (TDLU)-like morphologies/structures. No TDLUs were observed.

**Table S1. Milk sample information** (red crosses indicate missing data)

| **Sample Number** | **Infant age** | **Time** | **V Start** | **Nb of cells collected/ml** |
| --- | --- | --- | --- | --- |
| 1.1 | 15.0 | x | 15 | x |
| 1.2 |  | x | 10 | 1.15E+06 |
| 2.1 | 36.0 | x | 35 | 5.00E+04 |
| 2.2 | 39.0 | x | 40 | 2.88E+05 |
| 2.3 | 42.3 | x | 35 | 8.00E+04 |
| 2.4 | 49.9 | x | 30 | 1.50E+05 |
| 2.5 | 82.0 | x | 35 | 1.39E+05 |
| 3.1 | 16.5 | x | 5 | 1.41E+05 |
| 3.2 | 26.2 | 10:30 | 8 | 5.89E+05 |
| 3.3 | 28.1 | 10:00 | 7 | 3.47E+05 |
| 3.4 | 31.2 | 14:00 | x | x |
| 3.5 | 34.1 | 14:00 | 5 | 1.40E+05 |
| 4.1 | 140.0 | 14:00 | 4 | 1.40E+05 |
| 5.1 | 17.0 | 12:20 | 40 | 4.71E+05 |
| 6 | 10.0 | 13:00 | 120 | 2.20E+05 |
| 6.2 | 21.9 | x | 30 | 9.00E+05 |
| 6.3 | 32.1 | x | 35 | 1.95E+05 |
| 6.4 | 32.7 | x | 30 | 9.00E+04 |
| 6.7 | 41.7 | 11:00 | 25 | 4.86E+04 |
| 6.8 | 41.7 | 15:00 | 25 | 5.88E+04 |
| 6.9 | 41.9 | 12:00 | 30 | 9.33E+04 |
| 6.1 | 41.9 | 15:00 | 50 | 1.02E+05 |
| 6.13 | 43.6 | x | 30 | 1.17E+05 |
| 6.14 | 44.1 | 15:00 | 25 | 5.57E+04 |
| 7.1 | 28.0 | 15:30 | 30 | 1.56E+04 |
| 7.2 |  | x | 10 | 3.20E+03 |
| 8.1 | 7.0 | 12:30 | 30 | 1.25E+06 |
| 8.2 | 9.0 | 12:15 | 40 | 6.18E+05 |
| 9.1 | 20.0 | 10:03 | 5 | 6.80E+04 |
| 9.2 | 33.0 | 14:00 | 18 | 5.89E+04 |
| 10.1 | 24.0 | 11:30 | 60 | 7.34E+05 |
| 10.2 | 28.6 | 10:00 | 50 | x |
| 10.3 | 45.0 | 08:15 | 70 | 4.29E+05 |
| 10.4 | 47.1 | 09:45 |  | x |
| 11.1 | 21.0 | 12:05 | 180 | 6.42E+05 |
| 11.2 |  | x | 250 | 4.45E+05 |
| 12.1 | 16.0 | 11:50 | 30 | 5.80E+05 |
| 12.2 | 20.1 | 09:30 | x | x |
| 13.1 | 24.0 | 12:40 | x | x |
| 14.1 | 22.0 | 10:00 | 10 | 1.73E+04 |
| 15.1 | 16.0 | 10:00 | 6 | 3.83E+05 |
| 15.2 | 18.1 | 09:30 | 9 | 2.78E+05 |
| 15.3 | 20.0 | 10:30 | 10 | 1.66E+06 |
| 15.4 | 23.7 | 11:15 | 9.5 | 1.79E+06 |
| 15.5 | 27.7 | 10:45 | 13 | 1.04E+06 |
| 16.1 |  | x | x | x |
| 17.1 | 33.0 | 09:45 | 31 | 7.13E+05 |
| 17.2 | 35.0 | 10:00 | 17 | 8.38E+05 |
| 17.3 | 39.1 | 14:15 | 36 | 1.82E+04 |
| 17.4 | 41.0 | 14:15 | 26 | 2.67E+04 |
| 17.5 |  | 14:15 | 20 | 8.24E+04 |
| 18.1 | 7.0 | 09:00 | 20 | 2.23E+05 |
| 18.2 | 10.9 | 08:00 | 20 | 6.17E+05 |
| 19.1 | 18.0 | 10:20 | 10.5 | 1.25E+05 |
| 19.2 | 20.1 | 11:00 | 19 | 2.24E+05 |
| 19.3 | 22.1 | 12:30 | 65 | 3.10E+04 |
| 20.1 | 6.5 | 10:04 | 56.5 | 5.14E+05 |
| 21.1 | 9.0 | 13:40 | 41.5 | 4.10E+05 |
| 22.1 | 1.0 | 10:10 | 321 | 3.68E+05 |
| 23.1 | 3.0 | 12:00 | 47 | 6.58E+05 |
| 24.1 | 4.0 | 14:00 | x | x |
| 25.1 | 6.0 | 15:45 | 45 | 5.84E+04 |
| 26.1 | 3.0 | 09:00 | 80 | 1.11E+06 |

**Table S2. Cell culture media composition**

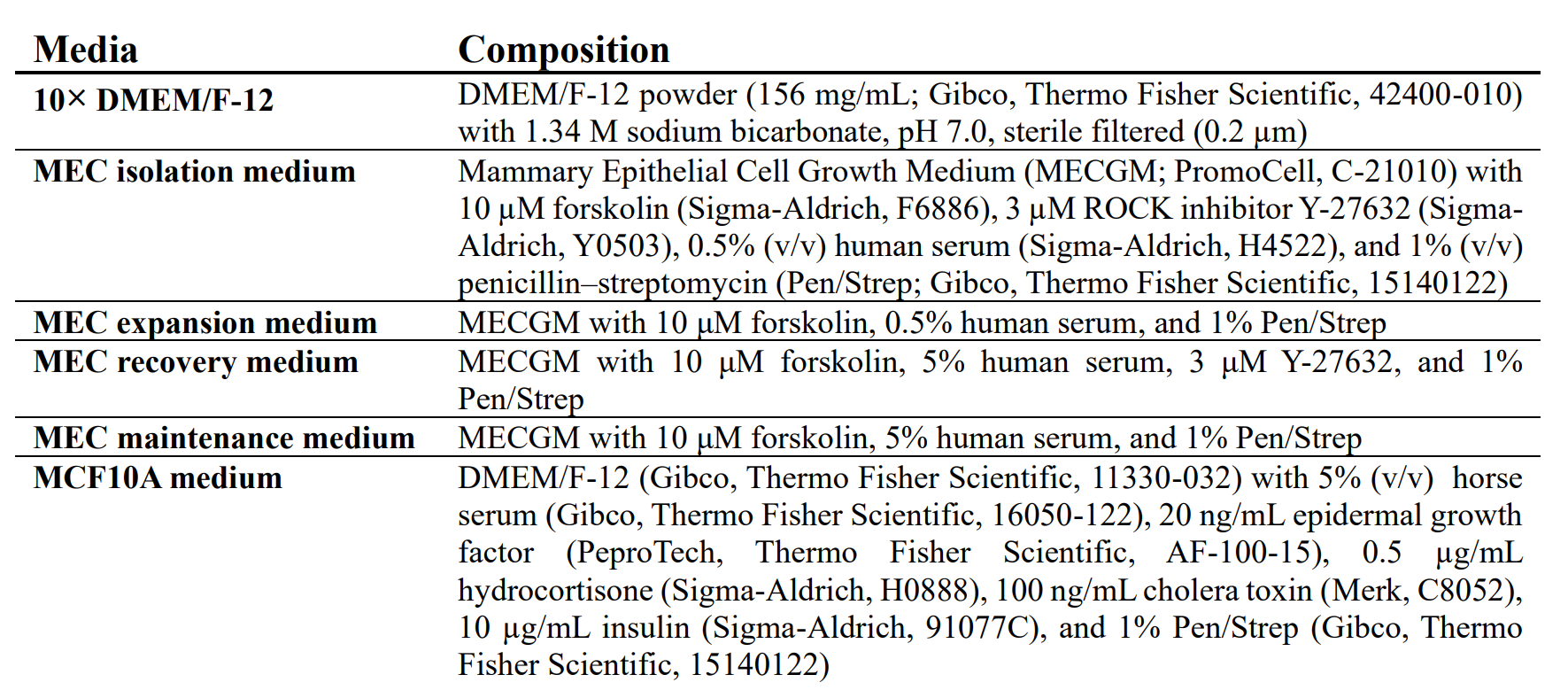

**Table S3. Organoid seeding densities**

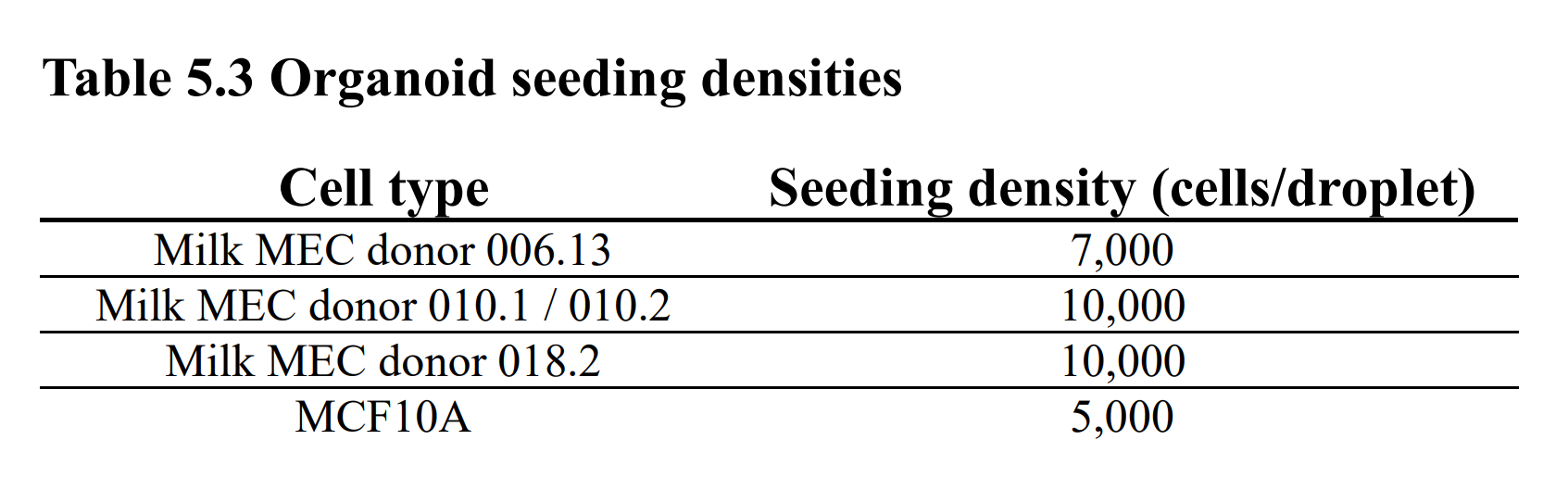

**Table S4. Solution and buffer composition**

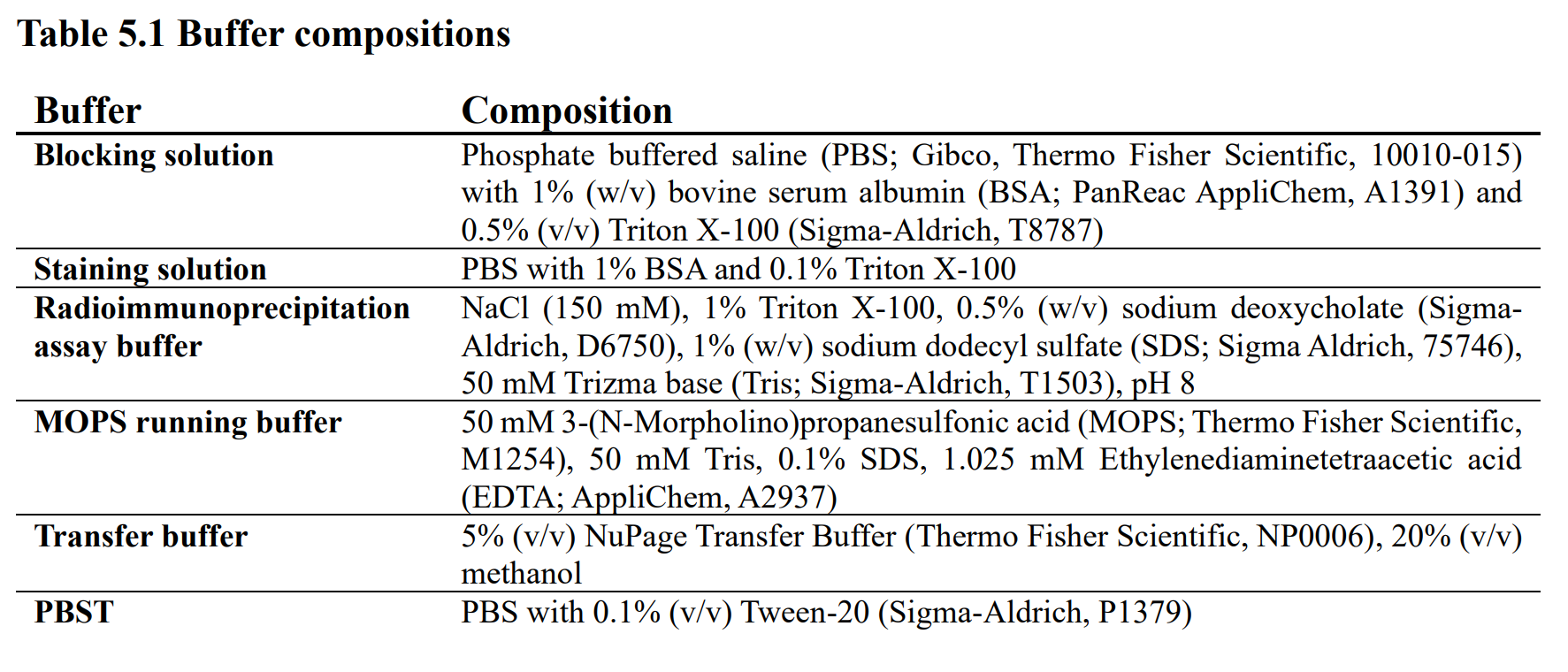

**Table S5. Primary antibodies**

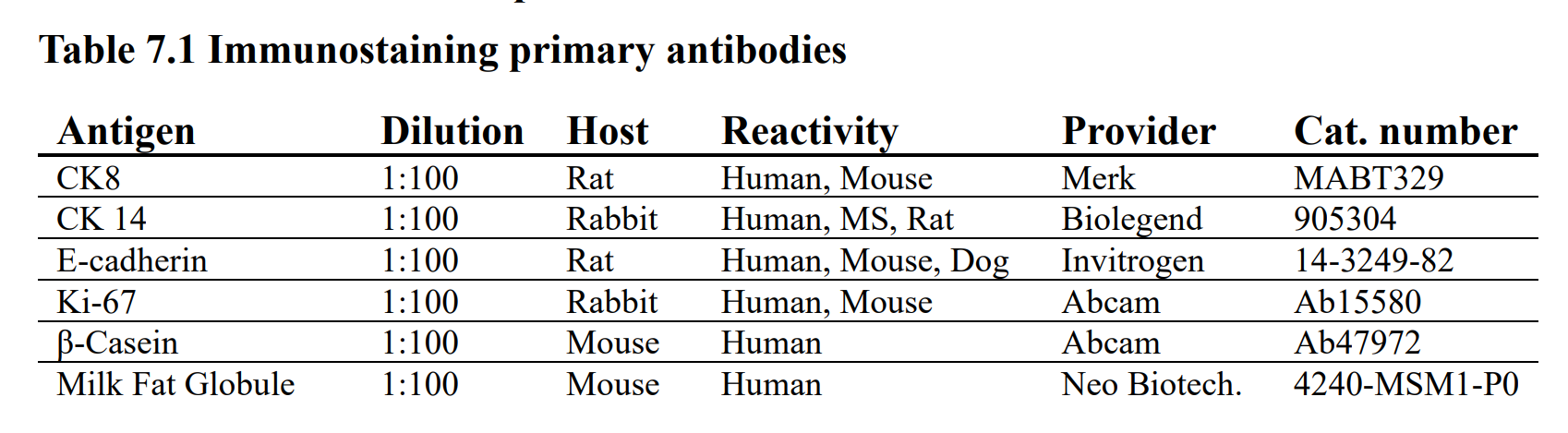

**Table S6. Secondary antibodies**

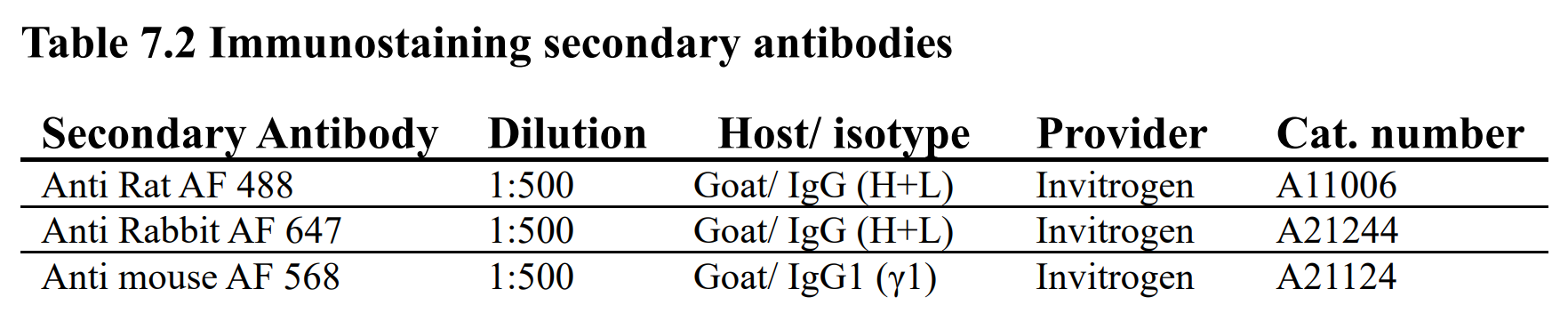

**Table S7. Western blot primary antibodies**

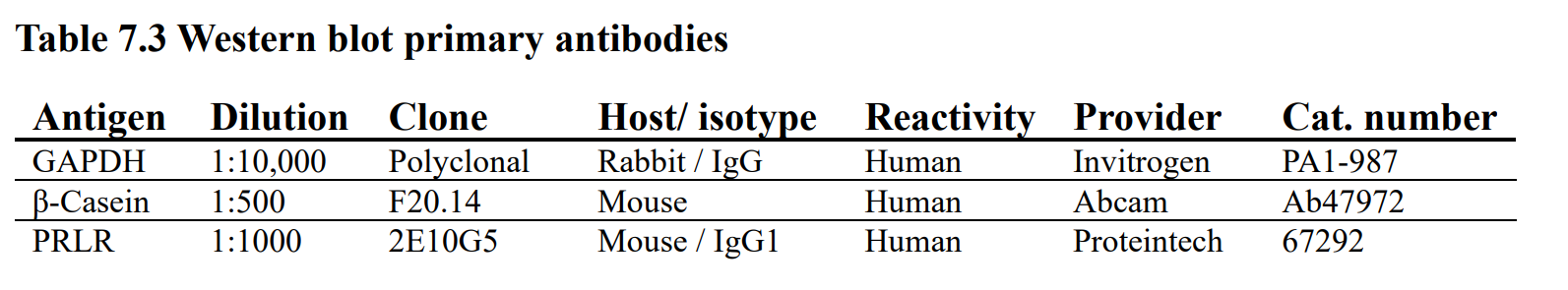

**Table 8. Western blot secondary antibodies**

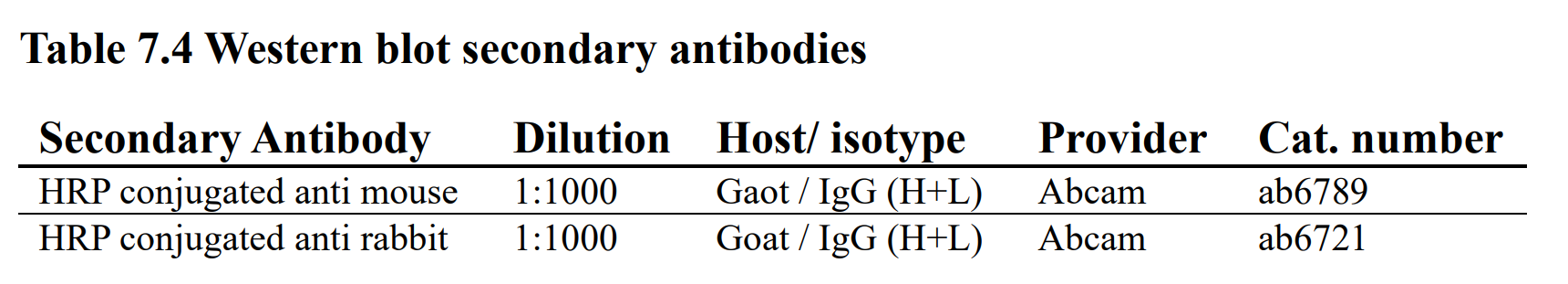

**Table 9. RT-qPCR Primers**

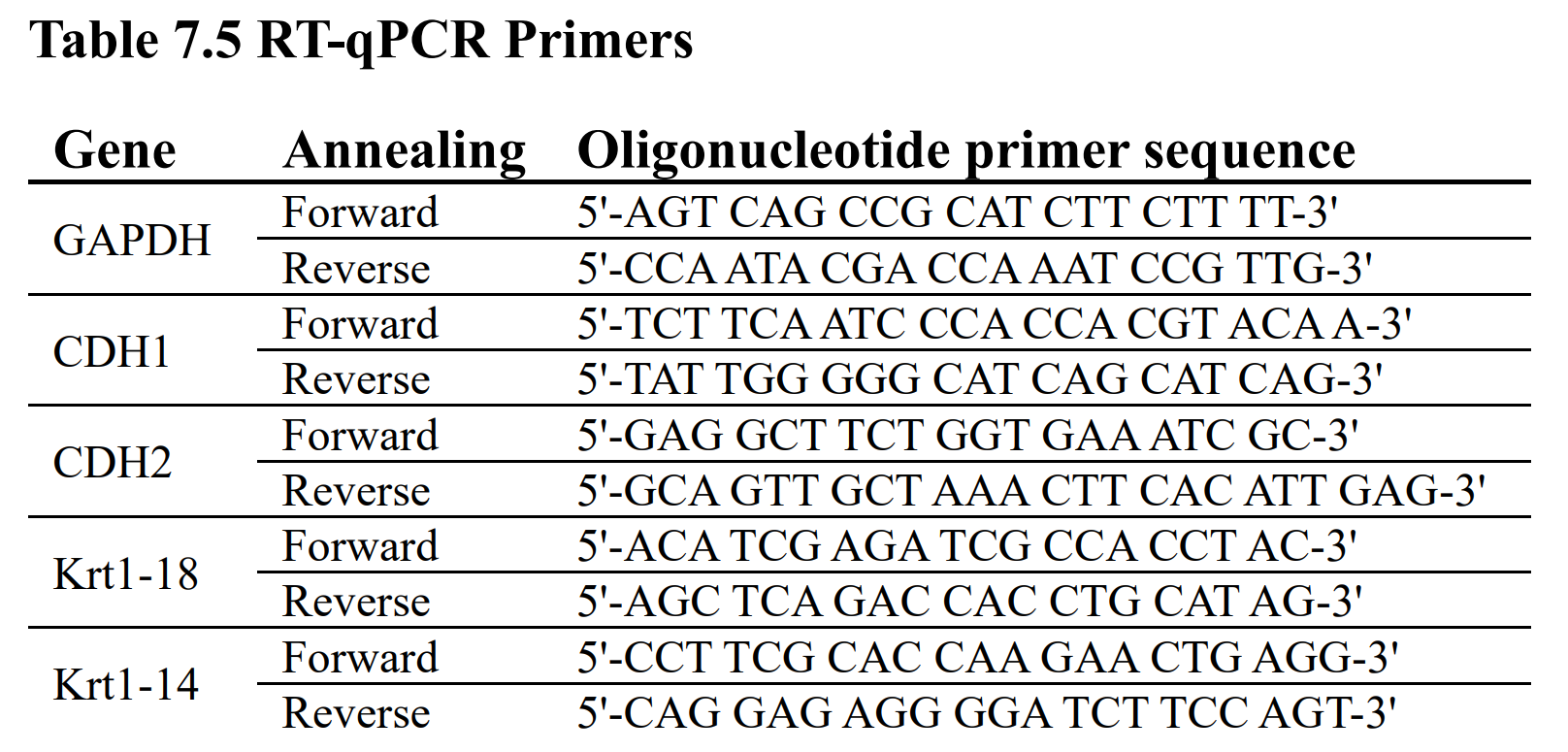

**Table S10. Flow cytometry antibodies**

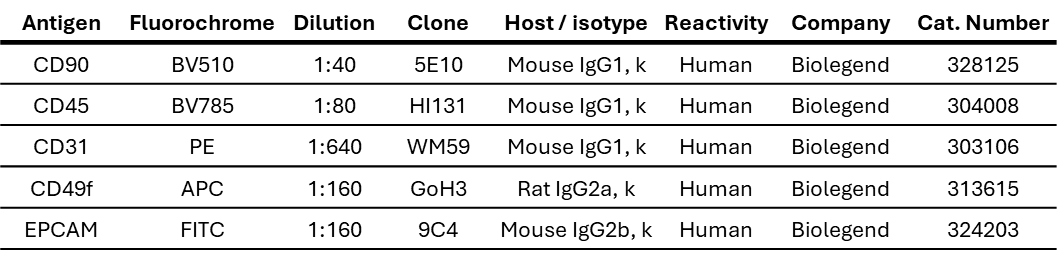
